# Single-particle studies of the effects of RNA-protein interactions on the self-assembly of RNA virus particles

**DOI:** 10.1101/2022.05.09.488235

**Authors:** Rees F. Garmann, Aaron M. Goldfain, Cheylene R. Tanimoto, Christian E. Beren, Fernando F. Vasquez, Charles M. Knobler, William M. Gelbart, Vinothan N. Manoharan

**Author notes:** These authors contributed equally to this work (Rees F. Garmann, Aaron M. Goldfain, Cheylene R. Tanimoto). Corresponding author, Rees F. Garmann, **Email:**.

## Abstract

Understanding the pathways by which simple RNA viruses self-assemble from their coat proteins and RNA is of practical and fundamental interest. Although RNA-protein interactions are thought to play a critical role in the assembly, our understanding of their effects is limited because the assembly process is difficult to observe directly. We address this problem by using interferometric scattering microscopy, a sensitive optical technique with high dynamic range, to follow the *in vitro* assembly kinetics of over 500 individual particles of brome mosaic virus (BMV)—for which RNA-protein interactions can be controlled by varying the ionic strength of the buffer. We find that when RNA-protein interactions are weak, BMV assembles by a nucleation-and-growth pathway in which a small cluster of RNA-bound proteins must exceed a critical size before additional proteins can bind. As the strength of RNA-protein interactions increases, the nucleation time becomes shorter and more narrowly distributed until the assembly kinetics become indistinguishable from diffusion-limited adsorption. In contrast, the time to grow a capsid after nucleation varies weakly with both salt and protein concentration. These results show that the nucleation rate is controlled by RNA-protein interactions, while the growth process is driven less by RNA-protein interactions and more by protein-protein interactions and intra-protein forces. The nucleated pathway observed with the plant virus BMV is strikingly similar to that previously observed with bacteriophage MS2, a phylogenetically distinct virus with a different host kingdom. These results raise the possibility that nucleated assembly pathways might be common to other RNA viruses.

RNA viruses first inspired the term “self-assembly.” Yet much is still not understood about how even the simplest such viruses assemble or if different viruses assemble in similar ways. Theoretical models suggest many possible assembly pathways, with many different roles for the RNA, but until recently measuring these pathways has not been possible. We use a sensitive microscopy technique to follow the assembly of individual particles of BMV, a plant virus. We find evidence of an RNA-mediated nucleation-and-growth pathway that is strikingly similar to that of MS2, a bacterial virus. The last common ancestor of BMV and MS2 existed only in ancient times, suggesting that their assembly pathway might be evolutionarily conserved and other viruses might follow a similar pathway.

## Introduction

Since the 1950s, the question of how RNA viruses self-assemble has inspired theoretical and experimental work in many fields of basic and applied science. Simple RNA viruses, which consist of a single-stranded RNA genome inside an ordered capsid made up of multiple copies of a single protein (**Figure 1A**), have served as model systems for studying the physical principles of structural virology involving virus particles of all shapes and sizes (1, 2). But the mechanisms and pathways by which these viruses assemble into the correct structure, while avoiding the many possible malformed structures, are not yet understood.

**Figure 1.**
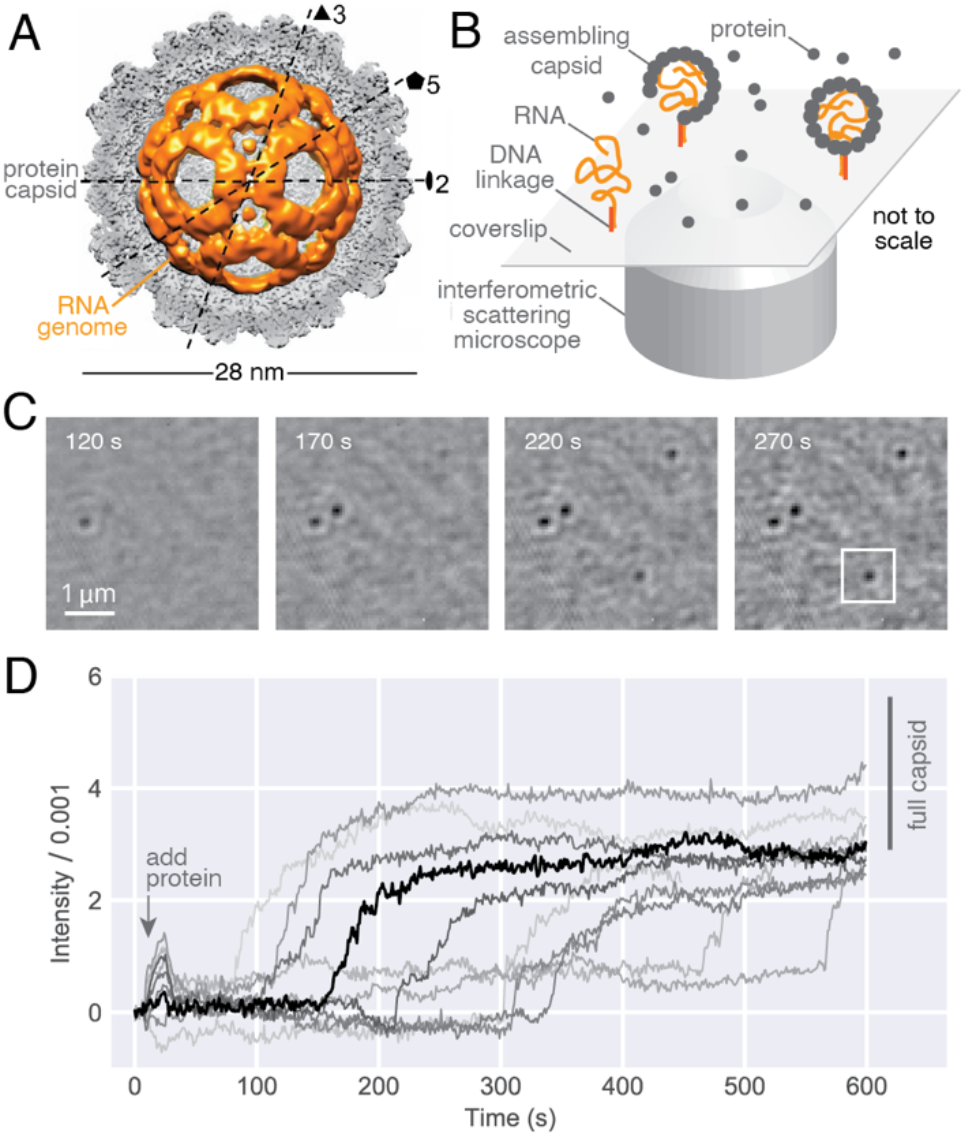
Overview of the system and the measurement. (A) A 3D model of BMV reconstructed from cryo-electron microscopy data (26) shows the protein capsid (gray) surrounding the RNA (gold). The model reveals most of the icosahedral capsid but only a small portion of the RNA, the rest of which adopts a disordered arrangement within the capsid. (B) A cartoon of the experiment shows viral coat proteins assembling around RNA strands that are tethered by DNA linkages to the surface of a functionalized glass coverslip. (C) The assembling proteins are imaged at 1000 Hz for 600 s using iSCAT microscopy. Each dark spot that appears in the images corresponds to proteins bound to an individual RNA strand. The darkness, or intensity, of each spot is proportional to the number of proteins bound to that RNA. The displayed images are the average of 1000 consecutive frames. (D) Traces of the intensity as a function of time (1000 frame moving average) reveal the assembly kinetics for each particle. Experimental conditions are 0.135 μmol/L protein and 250 mmol/L NaCl. The initial spike in intensity present in many of the traces is associated with vibrations introduced into the system as coat protein is injected. The thick, black trace corresponds to the boxed particle in (C). We compare the final intensities of the traces to the estimated intensity range of full capsids, which is shown as a vertical bar to the right of the traces.

Many different RNA viruses self-assemble, though the most well-studied come from two families: *Bromoviridae*, a family of plant-infecting viruses that includes brome mosaic virus (BMV) and cowpea chlorotic mottle virus (CCMV); and *Fiersviridae* (previously *Leviviridae*), a family of bacteria-infecting viruses that includes MS2 and Qβ. These families are as distinct phylogenetically as any two RNA virus families can be, having a last common ancestor that is thought to predate the emergence of eukaryotic cells (3). Accordingly, there are many well-established physical and biological differences among viruses in these families. Yet the four most studied members—BMV, CCMV, MS2, and Qβ—do have some structural commonalities: they are icosahedral viruses with a triangulation number (*T*) of 3, they have no lipid envelope, and each capsid surrounds about 3000-4000 nucleotides of single-stranded RNA.

The assembly of such structures is a nontrivial process. Identical coat proteins must adopt non-equivalent positions to make a *T*=3 capsid, with some arranging in pentagonal configurations and others in hexagonal configurations (2, 4, 5). Furthermore, these configurations must form in the correct proportions and positions for the capsid to close. Despite these challenges, assembly of virus-like particles of CCMV (6–8), BMV (7, 8), and MS2 (9) occurs in high yield even *in vitro* and in the absence of host-cell factors. The ability of viruses to avoid the many possible metastable states en route to complete assembly has been likened to the Levinthal paradox of protein folding (10, 11).

But unlike proteins, RNA viruses have a template for assembly: their own RNA. Current theoretical models of RNA virus self-assembly posit markedly different roles for the RNA, depending on the relative strengths of RNA-protein and protein-protein interactions, sequence-dependent RNA-protein interactions, RNA-mediated protein-protein interactions, and several other factors (12). Although specific interactions between RNA substructures and coat proteins have been hypothesized to help the virus avoid malformed configurations (11), viruses from different families differ greatly in their RNA structures and RNA-protein interactions. It is therefore unclear whether there are common features of the assembly process for different *T*=3 viruses or if there are distinct assembly pathways that depend on RNA-protein interactions.

Recent measurements of assembly kinetics suggest the latter: that assembly of viruses from different families follows different pathways. Fluorescence correlation spectroscopy (FCS) experiments (13, 14) of the kinetics of binding of MS2 coat protein and RNA indicate that assembly starts with a small cluster of RNA-bound proteins that trigger a change in the hydrodynamic radius of the RNA. In contrast, cryo-electron microscopy (15) and small-angle X-ray scattering (SAXS) (16) experiments of the assembly of CCMV coat protein and RNA show that disordered RNA-protein complexes formed at neutral pH anneal over several thousand seconds into well-formed capsids when the pH drops below 6.

But because these experiments involve different assembly conditions and different measurement techniques, their outcomes might not reflect fundamental differences in the assembly pathways of these viruses, but rather technical differences in the methods and protocols used to study them. Furthermore, most of the techniques that have been used do not measure the assembly process directly at the scale of individual particles because—one way or the other—they involve averaging over many particles. Such averaging can obscure the mechanisms and pathways that underpin stochastic assembly processes like viral assembly, in which each individual particle can follow its own unique sequence of intermediate states. Thus, it remains an open question whether a common assembly pathway might exist between these viruses.

We recently demonstrated that interferometric scattering (iSCAT) microscopy (17) can resolve the assembly kinetics of individual virus-like particles (18), providing a method to directly measure and compare the assembly pathways of different viruses. To perform the iSCAT experiment, we first tether viral RNA molecules to the surface of a functionalized glass coverslip under the desired buffer conditions (19) (**Figure 1B**). Next, we begin collecting iSCAT images of the RNA-decorated coverslip as we inject viral coat proteins at the desired concentration and in the appropriate buffer. As the proteins bind to the surface-tethered RNA, dark spots appear in the iSCAT images (**Figure 1C**). Subtracting the intensity associated with the RNA then yields images in which the intensity of each dark spot is proportional to the number of proteins that have accrued onto each individual RNA. Accordingly, plotting the trace of the intensity of a spot as a function of time reveals the assembly kinetics for that particle, and plotting the collection of traces reveals the assembly kinetics for the ensemble of particles (**Figure 1D**).

In our previous work (18) we examined the assembly of bacteriophage MS2. We found that the assembly kinetics were consistent with a nucleation-and-growth pathway in which a small cluster of RNA-bound proteins must exceed a critical size before the binding of additional proteins becomes favorable. Despite an apparently small critical nucleus size (a few coat-protein dimers), we found that MS2 capsids grow monotonically to full or nearly full size with high yield.

Although this previous study highlighted the importance of the RNA in the assembly process, the strong and specific RNA-protein interactions in MS2 (20–22) make it difficult to systematically address the central question of how the RNA affects the pathway. By contrast, the RNA in BMV interacts with the coat proteins through non-specific electrostatic interactions (23). As a result, the strength of RNA-protein interactions can be tuned by changing the ionic strength of the buffer solution (15, 24, 25). BMV therefore offers not only an interesting comparison to MS2—it is phylogenetically distinct but structurally similar—but also the means to understand the role of RNA-protein interactions.

In this study, we infer the assembly pathways of BMV from iSCAT measurements under different RNA-protein interaction strengths, allowing us to critically assess competing models of the assembly process. We follow the assembly trajectories of more than 500 individual virus particles under different assembly conditions, and we correlate the results with the absence and presence of ordered nucleocapsids as detected with electron microscopy. We find that BMV can assemble by a nucleation-and-growth process that is qualitatively similar to that of MS2. We show that the strength of RNA-protein interactions strongly affects the nucleation time, but only weakly affects the growth time, suggesting that RNA plays a central role in nucleating the viral capsid, but a relatively minor role in its growth. We discuss these observations in the context of recent models and hypotheses of RNA virus self-assembly.

## Results

### RNA-protein interactions can be controlled independently of protein-protein interactions

We first demonstrate that we can control the strength of RNA-protein interactions. By measuring the yield of RNA-protein binding between 20 nucleotide-long poly-U RNA and BMV coat-protein dimers (CP_2_) using a nitrocellulose binding assay (27), we find that increasing the concentration of salt decreases the amount of RNA-protein binding (**Figure 2A**). This result is consistent with binding being driven by electrostatic interactions between the negatively charged phosphate backbone of the RNA and the positively charged N-terminus of the coat protein (23).

**Figure 2.**
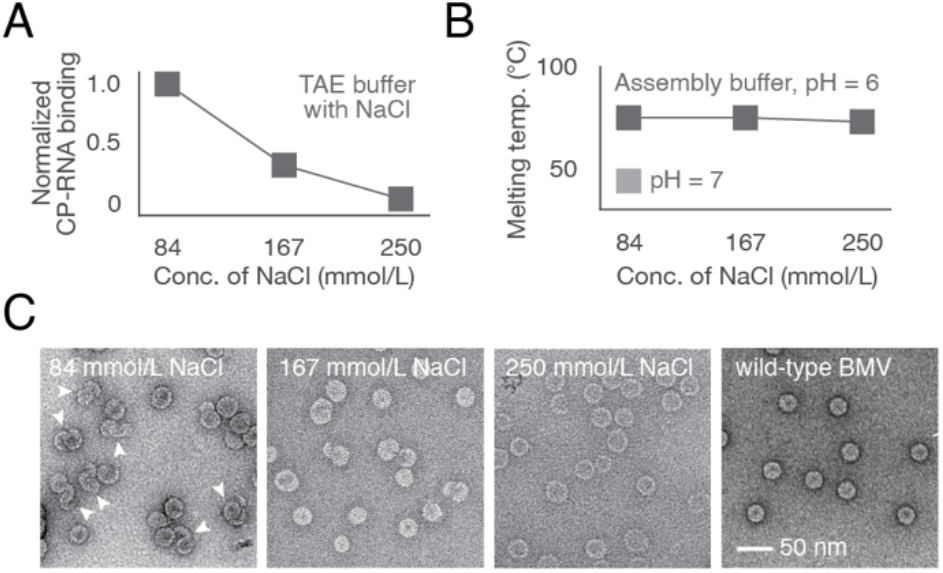
Ionic strength affects RNA-protein binding and the assembly process but does not affect the thermal stability of assembled particles. (A) Nitrocellulose binding measurements show that the yield of RNA-protein binding depends strongly on ionic strength. Measurements are performed at 4 nmol/L RNA and 0.043 μmol/L CP_2_ in TAE buffer with varying amounts of NaCl. The binding yield measurements are normalized with respect to the amount of binding at 84 mmol/L NaCl. (B) Differential Scanning Fluorimetry (DSF) measurements of the melting temperature of wild-type BMV in assembly buffer at pH 6 with varying NaCl, and at pH 7 with fixed NaCl, show that the capsid stability depends weakly on ionic strength, relative to pH, across the conditions tested. (C) Uranyl acetate negative-stain transmission electron microscope images of BMV particles assembled in assembly buffer at varying ionic strengths. We carry out assembly reactions using 7.5 nmol/L BMV RNA1 and 0.86 μmol/L BMV CP_2_, and assembly buffers at pH 6 and either 84 mmol/L, 167 mmol/L, or 250 mmol/L NaCl. The 84 mmol/L NaCl sample is notably heterogeneous, presenting many more malformed particles, which are denoted by arrowheads, compared with the higher-salt assemblies, which are more monodisperse. Wild-type BMV particles are shown as a reference.

To test whether varying ionic strength also affects protein-protein interactions, we perform Differential Scanning Fluorimetry (DSF) on wild-type BMV particles to study their thermal stability, a measure of lateral interactions between proteins in the capsid. We find that the thermal stability is approximately constant over the range of NaCl concentrations tested (**Figure 2B**). In contrast, the stability of BMV drops sharply when the pH is increased from 6 to 7, consistent with previous results (28, 29) showing that increasing pH weakens protein-protein interactions in BMV (**Figure 2B**). The absence of a change in stability with increasing salt is important because it suggests that ionic strength only weakly affects protein-protein interactions, and therefore that ionic strength can be used to specifically modify RNA-protein interactions.

Next, we use negative-stain electron microscopy to test for capsid formation as we vary ionic strength. In assembly reactions involving BMV CP_2_ and BMV RNA, we find that 84 mmol/L NaCl leads to heterogeneous assembly products, with some well-formed particles and many malformed particles, whereas 167 mmol/L and 250 mmol/L NaCl gives rise to more homogeneous spherical particles with roughly the same size and curvature as wild-type BMV (**Figure 2C**).

Taken together, these measurements show that we can control the strength of RNA-protein interactions, and that changing these interactions changes the assembly products. These results set the stage for iSCAT single-particle measurements that probe how the assembly pathways might change as RNA-protein interactions are varied.

### iSCAT measurements show that BMV assembles by a nucleation-and-growth pathway under certain conditions

We use iSCAT to measure the self-assembly kinetics of individual BMV particles as a function of the RNA-protein interaction strength, as controlled by the salt concentration (84 mmol/L, 167 mmol/L, and 250 mmol/L NaCl). We also vary the protein concentration (0.043 μmol/L, 0.135 μmol/L, and 0.427 μmol/L CP_2_) for a total of 9 possible experimental conditions. However, we did not test one of these combinations (250 mmol/L NaCl and 0.043 μmol/L CP_2_) because, given the results for the other 8 conditions, we expect the assembly timescales to be longer than our 600 s measurement time. In each of the 8 experimental conditions tested, we performed measurements in duplicate. In total, we measured and analyzed the assembly kinetics of 511 particles, 72 of which are shown in **Figure 3**.

**Figure 3.**
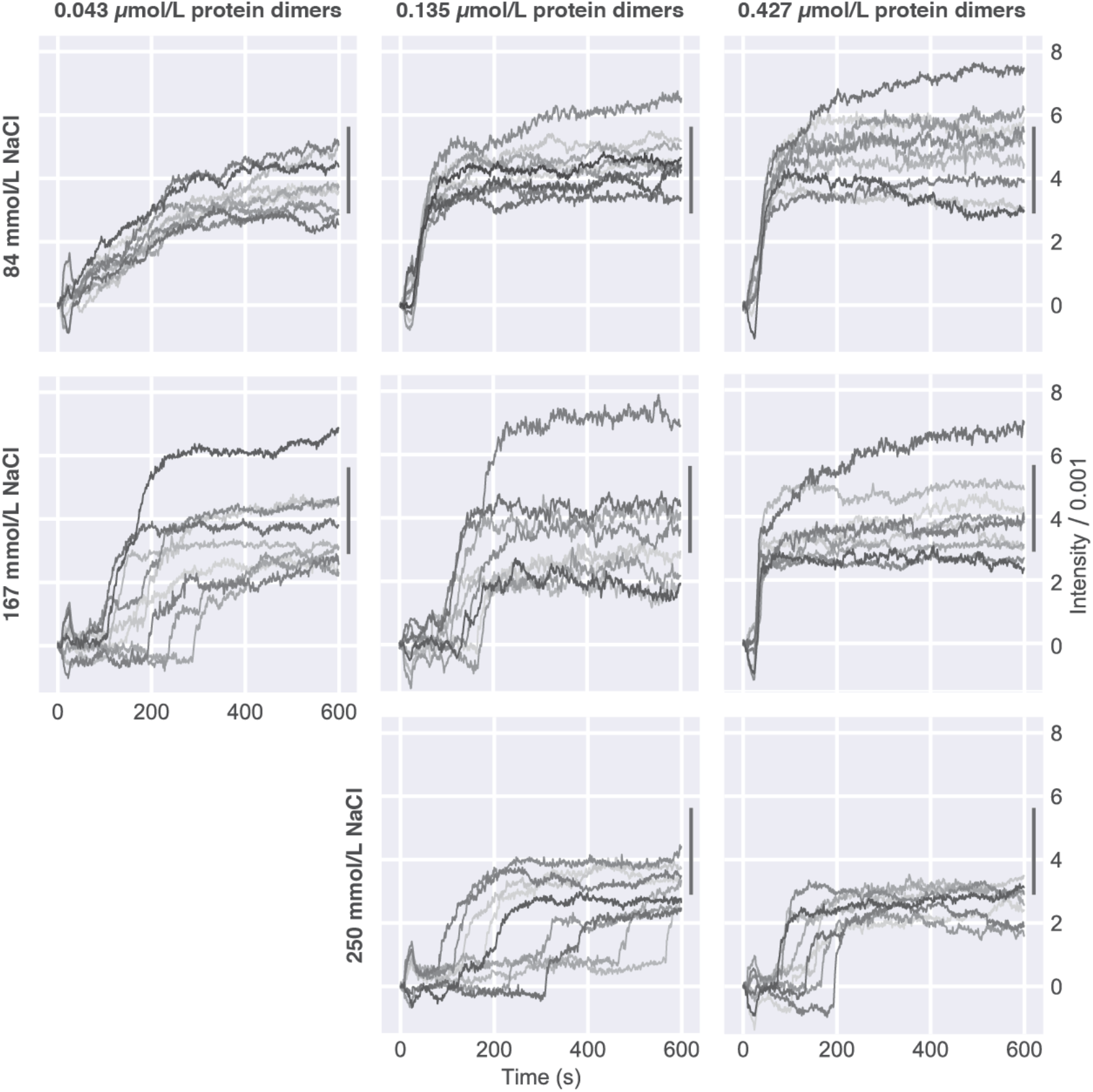
Single-particle measurements reveal qualitative differences in the assembly kinetics with varying ionic strength and protein concentration. Plots of traces showing intensity as a function of time from iSCAT experiments. Each trace represents the assembly of a single particle consisting of BMV coat protein and BMV RNA. We show 9 randomly selected traces at each condition (all 511 traces are shown in *Supporting Information*). The bars to the right of the traces show the range of intensities that are consistent with a full capsid.

We see qualitative differences among these single-particle traces for different RNA-protein interactions, most notably in the time at which each trace begins to increase rapidly (the “start time”). Consider the middle column in **Figure 3**, which corresponds to 0.135 μmol/L CP_2_. When RNA-protein interactions are strongest (corresponding to 84 mmol/L NaCl), each trace begins increasing immediately after the protein is introduced. In contrast, for weaker RNA-protein interactions (corresponding to 167 mmol/L NaCl), each trace remains at a low intensity for a variable amount of time before increasing, with some traces having start times greater than 100 s at the lowest protein concentration. When RNA-protein interactions are weaker still (corresponding to 250 mmol/L NaCl), the distribution of start times extends to longer values, with some traces having start times as large as 500 s at the intermediate protein concentration. Remarkably, one of the two experiments for 250 mmol/L NaCl did not result in any traces increasing above their initial value over the 600 s experiment.

These experiments allow us to rule out diffusion-limited accretion of proteins on the RNA as an assembly pathway for at least some of these conditions. If assembly were purely diffusion-limited (30), we would expect the onset of assembly to vary only with the concentration of protein. Instead, our results show that the start times vary with ionic strength at the same protein concentration.

The variation in start times points to a free-energy barrier to protein accretion on the RNA—in other words, a nucleation barrier (18). In classical nucleation theory, the barrier is associated with an initially unstable cluster—here, of proteins—becoming large enough that subsequent proteins bind favorably. Because nucleation is a stochastic process, the time required to form a sufficiently large cluster varies from particle to particle. These timescales are reflected in our start-time measurements. The observation that the distribution of start times broadens and shifts to larger values as RNA-protein interactions are weakened (but protein-protein interactions are not significantly changed) reveals that nucleation in BMV is a heterogeneous process, driven at least in part by attractive interactions between the RNA and the assembling proteins.

For the strongest RNA-protein interactions there is no obvious evidence of a nucleation barrier: At 84 mmol/L NaCl, the traces have nearly identical start times, as shown in the top row of **Figure 3**. At these conditions, we see that the final intensities of the traces, a measure of the number of proteins attached to the RNA, increase with increasing protein concentration. Moreover, the final intensities tend to decrease with decreasing RNA-protein interaction strength. This result is qualitatively consistent with our TEM measurements, which show heterogeneous, malformed, larger-than-wild-type particles for the strongest RNA-protein interactions (see structures highlighted by arrowheads in **Figure 2C**), and increasingly homogeneous, spherical, roughly wild-type-size particles as the RNA-protein interaction strength decreases. Understanding why the assembly products vary with RNA-protein interaction strength requires a more quantitative analysis of the assembly kinetics, and in particular how nucleation rates compare to growth rates.

### The shapes of the assembly traces reveal differences between nucleation and growth phases

To extract quantitative information from each recorded trace, we analyze the traces as shown in **Figure 4** to determine three kinetic parameters: the final intensity (*I*_*f*_), the start time (*t*_*s*_), and the growth time (*τ*_*g*_). The values of these parameters for experiments with 0.135 μmol/L CP and varying ionic strength are shown as histograms in **Figures 5A–C**.

**Figure 4.**
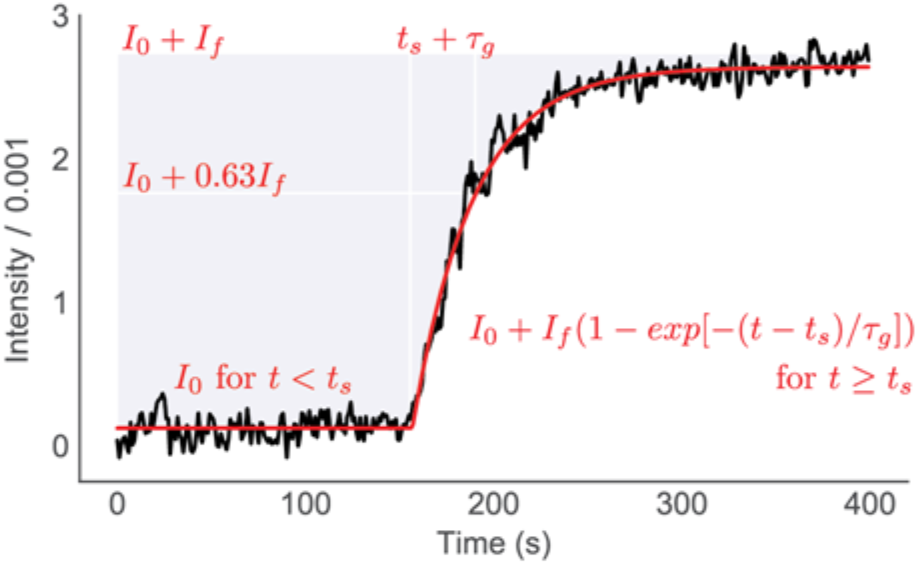
Extracting quantitative information from the assembly traces: *I*_*f*_, *t*_*s*_, and *τ*_*g*_. An intensity trace of the assembly of a single particle (black curve, corresponding to the boxed particle in Figure 1C) can be fit (red curve) to a piecewise function *I*(*t*) = *I*_0_ for *t* < *t*_*s*_; *I*(*t*) = *I*_0_ + *I*_*f*_(1 − exp[− (*t* − *t*_*s*_)/*τ*_*g*_]) *for t* ≥ *t*_*s*_, where *I*_0_ is the initial intensity (which can be offset from zero because of microscope drift), *I*_*f*_ is the final intensity, *t*_*s*_ is the start time (the time at which the trace begins to rise from its initial value), and *τ*_*g*_ is the growth time (the time for the trace to reach (1-1/*e*) = 0.63 of its final value once it has started increasing). The exponential function is not intended to imply a particular assembly mechanism (such as Langmuir adsorption) but rather is chosen because it is a simple function that fits the data well for most traces, allowing us to extract a characteristic growth timescale. In this way, we determine a set of three kinetic parameters, *I*_*f*_, *t*_*s*_, and *τ*_*g*_, for each assembling particle.

**Figure 5.**
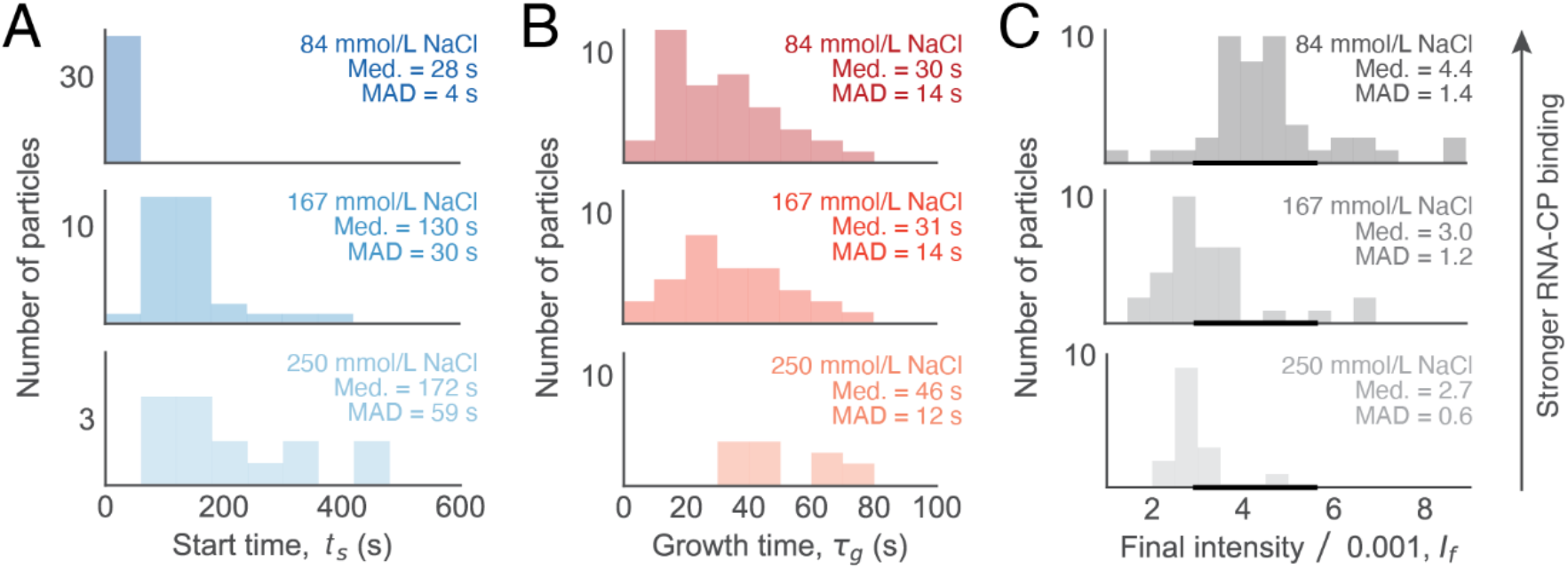
RNA-protein interactions strongly affect nucleation but only weakly affect growth. Histograms of start time (*t*_*s*_), growth time (*τ*_*g*_), and final intensity (*I*_*f*_) are plotted for three experiments with 0.135 μmol/L CP_2_ and either 84 mmol/L, 167 mmol/L, or 250 mmol/L NaCl: (A) Start times *t*_*s*_ increase and broaden with decreasing RNA-protein interaction strength; (B) growth times *τ*_*g*_ are less affected by RNA-protein interactions; and (C) the final intensity *I*_*f*_ decreases with decreasing RNA-protein interaction strength. The bold portion of the *x*-axis in (C) shows the size range for full capsids. In panels A–C, the median (Med) and median absolute deviation (MAD) of each fit parameter are listed. Both the median and MAD are robust to outliers that arise from the few traces that are fit poorly by the piecewise function (see *Supporting Information*).

Consistent with our qualitative analysis of the traces, the start-time histograms show that decreasing the strength of RNA-protein interactions increases the median start times and broadens their spread, as quantified by the median absolute deviation or MAD (**Figure 5A**). We find that the median start times increase by more than a factor of 6 and the MAD of start times increases by more than a factor of 14 when we weaken the RNA-protein interactions by increasing the salt concentration from 84 mmol/L to 250 mmol/L (**Figure 5A**). By contrast, the growth times and final intensities are more narrowly distributed and less affected by changes in RNA-protein interactions, showing an increase by a factor of roughly 1.5 in the median growth times and no increase in the MAD of the growth times (**Figure 5B**). The median final intensities show a corresponding decrease in particle size by factor of over 1.5 (**Figure 5C**).

These results show that RNA-protein interactions primarily affect the nucleation phase of the assembly pathway and only weakly affect the growth phase, if at all. Specifically, stronger RNA-protein interactions reduce the characteristic nucleation time until, at sufficiently large interaction strengths, it is no longer possible to tell if the assembly pathway is nucleated because all particles begin to assemble in apparent synchrony. We discuss this point in more detail in the *Discussion*.

To gain further insights into the nucleation kinetics, we examine the effect of protein concentration on the start-time distribution. We plot the MAD of start times, *t*_MAD_, for each protein concentration and each RNA-protein interaction strength (corresponding to values of salt concentration) in **Figure 6A**. In principle, the slope, *α*, of the fitted line represents the exponent in a power-law scaling *t*_*MAD*_ ∝ *c*^−*α*^, where *c* is the protein concentration. We find that for intermediate RNA-protein interaction strength (corresponding to 167 mmol/L NaCl), the measured start times become much more narrowly distributed as the protein concentrations increase, scaling with an exponent *α* = 1.5 ± 0.2. These results are once again consistent with assembly being a nucleated process—at higher protein concentrations the law of mass action drives more proteins onto the RNA, favoring larger clusters and shortening the time needed to form a nucleus.

**Figure 6.**
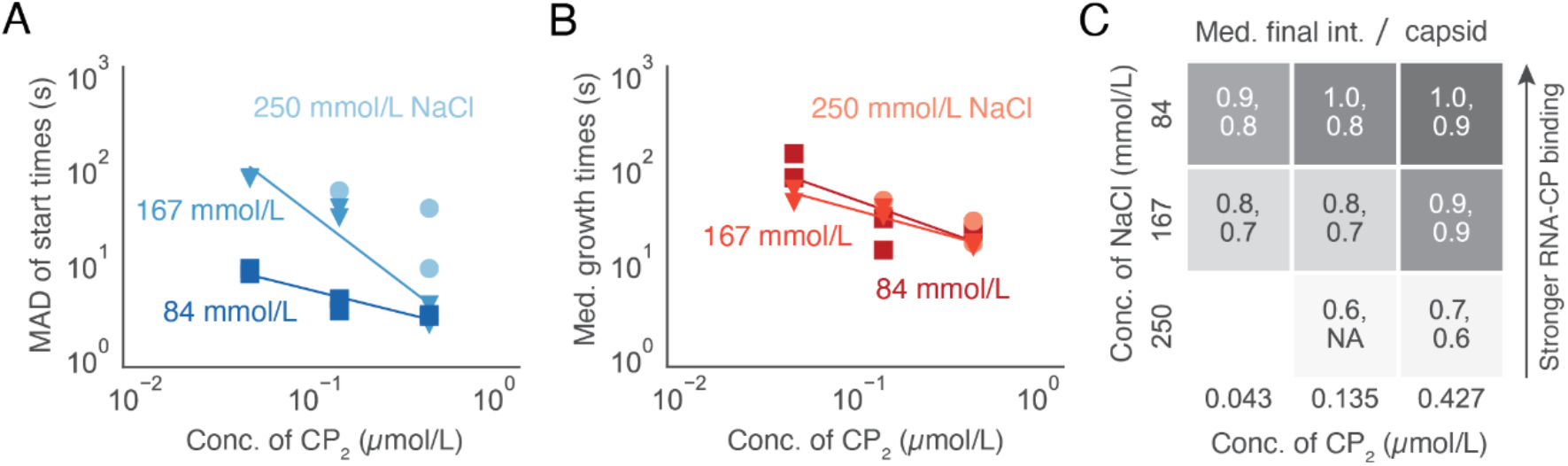
Experiments at varying protein concentration suggest that nucleation occurs by a collective process involving multiple proteins, while growth occurs by a lower-order process. (A) A log-log plot of the spread of start times as a function of protein concentration and RNA-protein interaction strength (squares: 84 mmol/L salt; triangles: 167 mmol/L; circles: 250 mmol/L salt). Duplicate measurements are plotted for each set of conditions except for 250 mmol/L salt and 0.135 μmol/L CP_2_, in which one of the duplicates did not yield assembly during the 600 s measurement time, and 250 mmol/L salt and 0.043 μmol/L CP_2_, which was not tested. Lines have been fit to the data from the 84 mmol/L and 167 mmol/L salt experiments to show how the dependence on protein concentration becomes steeper at higher salt (no line is fitted to the 250 mmol/L data because only two protein concentrations were tested at that salt concentration). (B) Log-log plot of the median growth times shows a concentration dependence that is roughly the same for all ionic strengths. (C) Heat map of the median final intensity, normalized by the intensity of a full capsid, of each duplicate measurement. The darkness of the gray color represents the magnitude of the intensity, which is proportional to the size of the assembled particle. While the intensities increase with increasing protein and decreasing salt, many of the assembled particles do not reach the size of a full capsid, indicating that they are missing proteins. These smaller particles might have partial capsids that fail to completely close around the tethered RNA strand. Duplicate measurements are separated by commas.

In contrast, for strong RNA-protein interactions (corresponding to 84 mmol/L NaCl), the start times remain narrowly distributed for all protein concentrations tested. Because the experimental uncertainties in our start-time measurements are on the order of seconds, small absolute differences in MAD values are not statistically significant. Thus, the line between nucleation and diffusion-limited aggregation becomes blurred. While it is possible that nucleation is occurring on a time scale too fast for us to measure, it is also possible that strengthening RNA-protein interactions qualitatively changes the assembly pathway such that it is no longer nucleated. We discuss these results further in the *Discussion*.

The growth kinetics are much less affected by varying either protein or salt concentration, as shown in **Figure 6B**. For all but one of the conditions tested, the median growth times differ by a factor of 2.5 at most. Only for 0.043 μmol/L CP_2_ and 84 mmol/L NaCl does the median growth time extend to 100 s. And while the median growth times tend to decrease with increasing protein, the amount by which they decrease is constant for all RNA-protein interaction strengths, as seen by the indistinguishable slopes of the best-fit lines in **Figure 6B** (slopes are 0.7 ± 0.3 at 84 mmol/L NaCl and 0.5 ± 0.1 at 167 mmol/L NaCl). Thus, while protein concentration appears to have a large effect on the start times, it has a much weaker effect on the growth times. Likewise, while RNA-protein interaction strength has a large effect on the start times, it does not significantly affect the growth times.

These results may explain why larger particles are favored at higher protein concentrations for all RNA-protein interactions tested, and at stronger RNA-protein interactions for all protein concentrations tested. These trends can be seen in the heat map shown in **Figure 6C**. In both cases—increasing protein concentration and increasing RNA-protein interaction strength—the nucleation times decrease relative to the growth times. Multiple nucleation events can therefore happen on the same RNA before any given nucleus has time to grow into a full capsid. Consequently, malformed structures consisting of multiple partially assembled capsids can occur. Indeed, the electron microscope images in **Figure 2C** show aggregates consisting of multiple capsid-like fragments that have the same curvature as the wild-type virus. The number and size of these aggregates decreases with decreasing RNA-protein interaction strength.

## Discussion

We discuss our results for the self-assembly kinetics of individual BMV particles in comparison with previous results using the same technique to study bacteriophage MS2 (18), focusing first on their similarities. By highlighting similarities in the assembly kinetics, we aim to identify common features of the assembly process that might be found in other RNA viruses.

The assembly traces for BMV are strikingly similar to those previously reported for MS2 (**Figure 7)** when RNA-protein interactions in BMV have been sufficiently weakened by sufficiently high salt concentrations (167 mmol/L or 250 mmol/L NaCl). For both viruses, we observe (i) broad distributions of start times that narrow with increasing protein concentration, consistent with a nucleation step; (ii) growth times that decrease with increasing protein concentration, but less rapidly than do the start times, consistent with growth involving a lower-order process; and (iii) increases in the fraction of overgrown particles with increasing protein concentration, with overgrown particles consisting of aggregates of partially formed capsids. We note that the concentration of coat protein used in the assembly of MS2 is roughly a factor of 10 higher than that used with BMV, related to BMV coat protein having a stronger overall binding affinity for RNA.

**Figure 7.**
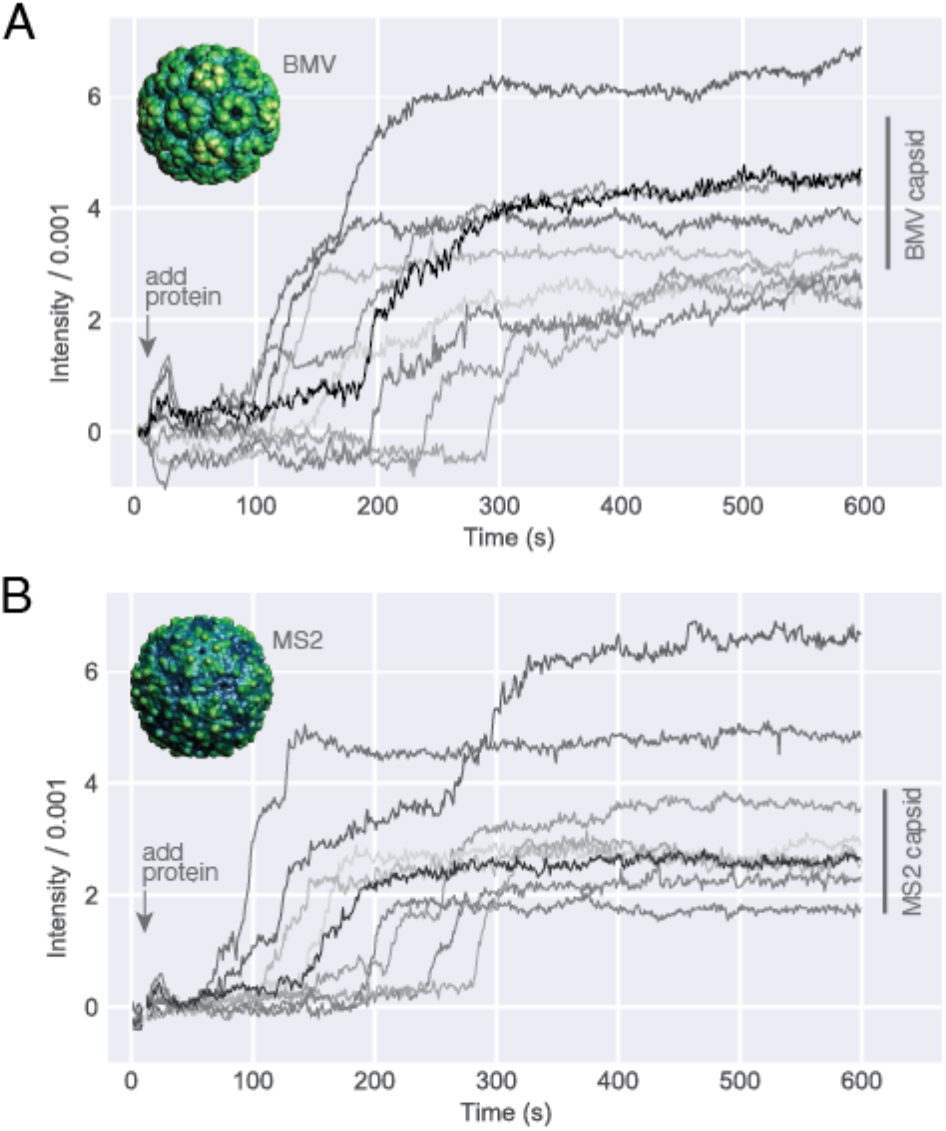
Comparison of the assembly traces for BMV and MS2 reveals similar nucleation-and-growth kinetics. (A) BMV assembly traces for 0.043 μmol/L protein and 167 mmol/L NaCl. (B) For comparison, we show traces recorded for bacteriophage MS2 assembly under conditions of 2 μmol/L protein and 84 mmol/L NaCl (data from (18)). Both sets of traces reveal a broad distribution of start times followed by rapid and monotonic increases in intensity. In both sets of traces, most traces plateau to a final value that is roughly consistent with a full capsid, with several traces plateauing to larger values. The bars to the right of the traces show the range of intensities that are consistent with a full capsid. These ranges are different for MS2 and BMV, reflecting the different molecular masses of their coat proteins.

Another interesting similarity between BMV and MS2 assembly is that the apparent critical nucleus size is small—a few proteins—in both cases. We estimate this nucleus size from the amplitude of the fluctuations prior to the start time. In nearly all traces under conditions of moderate to weak RNA-protein interactions (167 mmol/L or 250 mmol/L NaCl), these fluctuations are comparable to the noise level, which corresponds to the scattered intensity from a few CP_2_. We use the term “apparent critical nucleus size” because averaging reduces our ability to observe large sub-critical fluctuations. Nonetheless, this small apparent critical nucleus is comparable to that observed for MS2, and for both viruses we find that growth proceeds monotonically for nearly all traces (**Figure 7A and B**).

These results raise the question of how either BMV or MS2 is able to grow so rapidly, in a way that ensures the capsid has the correct curvature, from such an apparently small nucleus. Nucleation by itself cannot solve the “Levinthal paradox” of virus assembly because the nucleus is too small to dictate the formation of a *T*=3 structure during growth. And yet, following nucleation, both BMV and MS2 display essentially monotonic growth to the size of a full capsid, with little if any disassembly or backtracking along the way. Furthermore, the curvature of the assembly products observed by TEM is similar to that of wild-type capsids even when the products are malformed aggregates of partially assembled capsids. How, then, do the coat proteins add to a growing nucleus, forming both the pentamers and hexamers needed for the proper curvature, without frequent stalling events or needing to detach from the particle?

A recent hypothesis is that specific subsequences on the RNA mediate the growth process to produce the correct curvature and structure. These subsequences are sometimes called “packaging signals”—a term that has come to refer to any elements of the RNA secondary structure that have high local affinity for coat protein. A model for MS2 assembly has been proposed in which upwards of 60 strategically positioned packaging signals (11, 31) guide assembly of the coat proteins. While not providing direct evidence in support of this model, high-resolution cryo-electron microscopy studies of MS2 particles show a dozen or so specific interactions between the RNA genome and its surrounding protein capsid (32), suggesting that packaging signals play a role in ordering the genome. But BMV is different. The absence of such order in the structure of the packaged BMV genome (26), and the lack of evidence of strategically positioned packaging signals in BMV RNA, argue against there being a general paradigm that depends on a special distribution of specific RNA-protein interactions.

It is therefore interesting that we observe similar assembly kinetics for BMV and MS2. Our measurements suggests that whatever role packaging signals play in the self-assembly of these two viruses, their effect appears to be at most quantitative, with the qualitative features of the assembly process determined by general non-specific interactions between the assembling proteins and the RNA. Determining the quantitative effects of packaging signals will ultimately require kinetic experiments in which the RNA sequence and secondary structure is controlled (33).

Our experiments with BMV show that one should look to protein-protein interactions and intra-protein forces, rather than RNA-protein interactions, to understand how the capsid grows after nucleation. The median growth time does not vary with salt concentration (**Figure 5B**), indicating that RNA-protein interactions are not involved in the rate limiting step of the growth phase of assembly. Protein-protein interactions or intra-protein forces must therefore direct the rate-limiting step. We have also shown that the median growth time varies less strongly with protein concentration than does the start time (**Figure 6A-B**). A simple scenario consistent with these observations is that the RNA does not play a significant role in the assembly process after nucleation. Instead, during the growth phase, new proteins bind directly to other proteins in the assembling particle. These incoming proteins bind individually, not as clusters.

Thus far we have discussed the similarities between BMV and MS2 assembly, but there are differences that arise when RNA-protein interactions are strong. Under conditions of 84 mmol/L NaCl and 0.043 μmol CP_2_, experiments on BMV assembly show a small spread in start times and relatively large growth times (see upper left plot in **Figure 3**). With MS2, we found that when the spread in start times is smaller than the average growth time, many oversized structures form. Our interpretation was that multiple nucleation events occur on the same RNA strand before the first nucleus can grow and sequester the RNA. With BMV, we observe similar overgrown structures but only at higher protein concentration (**Figure 2C)**. We do not observe a significant number of overgrown structures at low protein concentration. One possible explanation is that assembly is diffusion-limited when RNA-protein interactions are strong, occurring with no nucleation barrier, and that the small observed variation between traces is due to measurement noise. Alternatively, it is possible that the assembly is nucleated, but the nucleation time is too small to resolve. In *Supporting Information*, we perform simulations to determine if our measurements are consistent with any of the above scenarios—a diffusion-limited pathway subject to measurement noise or a nucleated pathway with a small nucleation time. These simulations suggest that our measurements are consistent with either of these pathways. Thus, additional experiments that can resolve smaller spreads in start times are needed to determine the assembly pathway when RNA-protein interactions are strong.

There also remains the question of if and when an ordered capsid arises from the RNA-protein complex. Interferometric scattering measurements do not address this question because the intensity depends primarily on the number of RNA-bound proteins and weakly, if at all, on their structure. There are at least two possibilities for the pathway underlying those traces that do not show a clear separation between nucleation and growth. One is a barrier-less formation and growth of partially to completely ordered capsids, and the other is a barrier-less accretion of disordered proteins, followed by the onset of order among the bound proteins. These pathways cannot be distinguished in iSCAT because the measurement is blind to the onset of order. The same is true even when there is a clear separation between nucleation and growth processes. Here we observe a barrier to protein accretion, but it is possible that the proteins are not ordered. An analogous situation occurs in a bulk phase transition when gas-to-solid condensation takes place through a liquid-state intermediate. Thus, while our measurements can definitively resolve the presence of a barrier to protein accretion, future structural measurements are needed to distinguish whether the barrier involves ordered or disordered proteins.

## Conclusions and Future Directions

The iSCAT experiments, with their high temporal resolution and ability to resolve the kinetics of assembly of individual viral capsids, offer the most detailed view to date of virus self-assembly pathways. Although iSCAT cannot reveal at what stage the capsid becomes ordered, much information can be gleaned by combining the technique with electron microscopy and bulk assays, as we have shown.

These experiments have shown that when RNA-protein binding is weak, the assembly kinetics of BMV are similar to those of MS2, involving a nucleation phase in which small numbers of proteins bind to the RNA, followed by a monotonic growth phase in which a capsid-worth of proteins steadily accrues. When RNA-protein binding is strong, we observe no clear barrier to protein accretion and hence no separation between nucleation and growth. Because our method may obscure nucleation events that occur on fast timescales, future studies are needed to determine whether for strong RNA-protein interactions there is a qualitatively different assembly pathway involving a saturated—”en masse” (15, 34)—adsorption of proteins on the RNA.

When the assembly is nucleated, we have shown that the time required to form a nucleus depends strongly on the strength of RNA-protein interactions, whereas the time needed to accrue a capsid-worth of proteins does not, suggesting that the RNA plays a more central role in the nucleation phase than in the growth phase. These results are consistent with protein accretion being a heterogeneous process in which subcritical protein clusters form on—and are stabilized by—the RNA. This role of the RNA degree of freedom—involving both RNA-protein binding and the conformation of the RNA itself (35)—compromises the notion of the nucleus being a fixed arrangement of coat proteins, such as a hexamer (36) or a pentamer (37) of dimers, although the uncertainties in our measurements cannot rule out this possibility. Future structural experiments, or kinetic experiments that address a broader range of timescales, may clarify whether the RNA enables different nucleus structures in different conditions.

Our experiments also reveal important features of the growth process. The shapes of the iSCAT traces show that growth can take place rapidly and without significant errors starting from an apparently small critical nucleus of only a few proteins. Furthermore, the weak dependence of growth times on protein concentration suggests that growth occurs by a lower-order process, such as the addition of individual protein subunits from solution, rather than higher-order collective processes involving clusters of proteins.

The model of capsid assembly that emerges from our results is as follows: the strength of RNA-protein interactions primarily controls the kinetics of nucleation but not the local structure or curvature of the capsid, while protein-protein and intra-protein interactions control the growth phase and the emergence of the *T*=3 structure. This model is compatible with simulations and theory showing that the growth of a *T*=3 structure is driven by minimization of the elastic energy of the capsid (38, 39), which in general is related to the stretching and bending of coat-protein dimers as well as of the bonds between them. The elastic-energy hypothesis could explain why BMV and MS2 show similar assembly pathways.

To test this hypothesis, future experiments and analysis might focus on the growth process rather than nucleation. Recently, BMV, MS2, and Qβ capsids of different sizes and symmetries have been observed (40–45). Experiments that determine how such malformed or overgrown capsids form would shed light on the interactions that control growth. Another useful next step is to develop coarse-grained computer simulations (34) that include the diffusion of proteins to the surface that is inherent to the iSCAT experiments. With such models it would be possible to directly compare ensembles of simulated kinetic traces to the ensembles of traces measured with iSCAT. If agreement between these simulations and our data can be obtained, it would point the way toward a detailed mechanism of the nucleocapsid formation process.

The commonalities in the *in vitro* assembly pathways of BMV and MS2 are remarkable in light of the vast phylogenetic distance between these viruses, their different RNA structures, and the differences in the specificity and strength of RNA-protein interactions. We do not know if the same pathways are operative *in vivo*, and there is reason to suspect that, at least for MS2, there may be differences between the *in vivo* and *in vitro* pathways. Whereas wild-type BMV particles consist of only two components, the RNA genome and (180 copies of) the capsid protein, wild-type MS2 particles contain a single copy of a different gene product (the “maturation” protein) that replaces a coat protein dimer and breaks icosahedral symmetry. The *in vitro* assemblies of MS2 are reconstituted from viral RNA and capsid protein alone, without the maturation protein.and with *T*=3 structure.

Nonetheless, the existence of two different icosahedral, *T*=3 capsids that assemble *in vitro* in similar ways is intriguing from both a physical and evolutionary perspective. From a physical perspective, this result suggests that the assembly of *T*=3 viruses might be understood through a general physical theory. From an evolutionary perspective, it highlights the question of how icosahedral viruses with quasi-equivalent (*T* > 1) capsid subunits evolved. In MS2, for example, a single point mutation in the coat protein changes the structure of the capsid from *T*=3 to *T*=1 (45). Because capsids are self-assembled, such mutations can in principle also change the assembly pathway. Future studies might use interferometric scattering to explore how mutations affect the assembly of virus-like particles from different virus families, with the aim of discovering whether there are conserved interactions that promote robust assembly of the *T*=3 structure.

## Materials and Methods

### Buffers used

**Disassembly buffer**: 50 mmol/L Tris-HCl, pH 7.5; 500 mmol/L CaCl_2_, 1 mmol/L ethylenediamine tetraacetic acid (EDTA), 1 mmol/L dithiothreitol (DTT), 0.5 mmol/L phenylmethylsulfonyl fluoride (PMSF). **Protein storage buffer**: 20 mmol/L Tris-HCl, pH 7.2; 1 mol/L NaCl; 1 mmol/L EDTA; 1 mmol/L DTT; 1 mmol/L PMSF. **TAE buffer with NaCl**: 40 mmol/L Tris-HCl pH 8.3, 20 mmol/L acetic acid, 1 mmol/L EDTA; and 84 mmol/L, 167 mmol/L, or 250 mmol/L NaCl. **Assembly buffer**: 42 mmol/L 2-(N-morpholino)ethanesulfonic acid (MES), pH 6; 84 mmol/L, 167 mmol/L, or 250 mmol/L NaCl; 8.4 mmol/L MgCl_2_; and 3 mmol/L acetic acid. For DSF measurements, we also prepared assembly buffer at pH 7 by replacing MES with sodium phosphate.

### Synthesis of BMV RNA1

BMV RNA1 was made by in vitro transcription of the DNA plasmid pT7B1, linearized with BamHI (New England Biolabs, USA)*, with a T7 polymerase transcription system (Thermo Fisher, USA) and purified with an RNEasy Mini Kit (Qiagen, DEU), both following the manufacturers’ specifications.

### BMV coat protein purification

BMV was purified from infected barley leaves (*Hordeum vulgare*)(46), and coat protein was purified as described previously (47). Nucleocapsids were disassembled by dialyzing against disassembly buffer at 4 °C overnight. The RNA was pelleted and the coat protein isolated by ultracentrifugation at 90,000 rotations per minute for 100 min at 4 °C in a Beckman TLA110 rotor. Coat protein was extracted from the supernatant and immediately dialyzed against protein storage buffer. Protein concentration and purity were assessed by UV-Vis spectrophotometry; only protein solutions with 260/280 ratios less than 0.6 were used for assembly. Protein was frozen in liquid nitrogen and stored at −80 °C until ready to use, at which point it was defrosted on ice and stored at 4 °C for up to two weeks.

### Measuring the binding yield between fluorescently labeled 20U RNA and BMV CP using a nitrocellulose-binding assay

Fluorescently labeled 20U RNA at a concentration of 4 nmol/L was mixed with 0.043 μmol/L BMV CP_2_ in TAE buffer with either 84 mmol/L, 167 mmol/L, or 250 mmol/L NaCl, and left for 30 min at room temperature. The RNA was labeled at its 5’-end with an AlexaFluor647 dye (Integrated DNA Technologies, USA). 250 μL aliquots of each RNA-protein mixture were passed through a nitrocellulose membrane (0.45 μm pore size; Thermo Fisher, USA) that was presoaked in TAE buffer with 250 mmol/L NaCl using a 96-well dot-blot apparatus (Biorad, USA) under weak vacuum. Then the membrane was washed by passing through an additional 500 μL of the corresponding assembly buffer. The amount of membrane-bound RNA was quantified at each salt concentration by measuring the fluorescence emission intensity of the AlexaFluor647 dye using a fluorescence scanner. The amount of protein bound to the membrane was constant for all salt concentrations, as determined by staining with Ponceau S solution (MilliporeSigma, USA). Because protein is retained by the membrane, but free RNA is not, the intensity of membrane-bound RNA is taken to be proportional to the yield of RNA-protein binding.

### Measuring the apparent melting curve of BMV by differential scanning fluorimetry (DSF)

WT BMV was dialyzed against assembly buffer with pH 6 or pH 7 overnight at 4 °C. Aliquots of WT BMV at a final concentration of 0.2 mg/mL, 2.5x SYPRO orange fluorescent dye (Molecular Probes, USA), and a final salt concentration of 85 mmol/L, 167 mmol/L, or 250 mmol/L NaCl were prepared. DSF was performed in triplicate in a 96-well plate CFX Connect quantitative PCR machine (Bio-Rad, USA). All samples were heated from 25 °C to 95 °C, in 1 °C increments with a 1-min stabilization period at each temperature before the sample was measured. Excitation/emission wavelengths of 470/550 nm were used to detect the fluorescence emission of SYPRO orange binding to hydrophobic regions of coat protein exposed upon capsid disassembly. The apparent melting temperature is defined as the temperature at which the derivative of the fluorescence emission signal (−*dI*/*dT*) is maximal.

### Negative-stain electron microscopy

Negative-stain electron microscopy was used to image the protein structures that assemble around RNA in solution. The assembly reaction was carried out in assembly buffer by mixing 860 nmol/L BMV CP_2_ with 7.5 nmol/L BMV RNA1 and incubating at room temperature for 10 min. 6 μL of assembly reaction was deposited on glow-discharged carbon-coated copper (200-mesh) PELCO Pinpointer grids (Ted Pella, USA). After 1 min, the grids were blotted with Whatman filter paper, and then stained with 6 μL of 2 % uranyl acetate for 1 min followed by complete stain removal and storage in a desiccator overnight. Micrographs were acquired using a Tecnai G2 TF20 High-Resolution electron microscope (FEI, USA) with an accelerating voltage of 200 kV. Images were collected at 3 μm to 4 μm underfocus with a TIETZ F415MP 16-megapixel CCD camera (4000 by 4000 pixels, pixel size 15 μm).

### iSCAT experiments

iSCAT measurements were performed as described in detail in Reference (18). In brief, a spatially filtered, 450 nm diode laser was coupled into an oil-immersion objective to illuminate a small region of the coverslip with a collimated beam. Three-dimensional active stabilization was used to extend the measurement duration. We estimate the intensity range of full BMV capsids by scaling the measured intensity range for MS2 capsids (18) by the relative mass of BMV and MS2 capsids. Coverslips were functionalized with a layer of polyethylene-glycol (PEG) molecules, about 1 % of which were functionalized with a 20-base DNA strand (5’-GGTTGGTTGGTTGGTTGGTT-3’), to which BMV RNA1 strands were tethered using a 60-base DNA linker strand (5’-CCGTGGTCGACAAGGGATTGAACCTCGTTCCGTGGTCTACAACCAACCAACCAACCAACC-3’). Assembly kinetics experiments were performed at room temperature in assembly buffer with 84 mmol/L, 167 mmol/L, and 250 mmol/L NaCl and protein concentrations of 0.043 μmol/L, 0.135 μmol/L, and 0.427 μmol/L BMV CP_2_.

*Certain commercial materials and equipment are identified in order to adequately specify the experimental procedure. Such identification does not imply recommendation by the National Institute of Standards and Technology.

## Acknowledgments

We thank Ben Rogers and Abigail Chapman for many helpful discussions. Research reported in this publication was supported by the National Institute of General Medical Sciences of the National Institutes of Health (Award Number R00GM127751 to RFG) and the National Science Foundation Molecular and Cellular Biosciences Division (Awards MCB 1716925 and 2103700 to WMG). This research was partially supported by NSF through the Harvard University Materials Research Science and Engineering Center under NSF grants DMR-1420570 and DMR-2011754. AMG was partially supported by an NSF Graduate Research Fellowship under Grant DGE-1144152. We acknowledge the California Metabolic Research Foundation for its support of biochemical research at San Diego State University, and the Bauer Core Facility at Harvard University for shared experimental facilities used in this study.

