## Supporting Information for "Single-particle studies of the effects of RNA-protein interactions on the self-assembly of RNA virus particles"

**Supporting Information for: Single-particle studies of the effects of ionic strength and protein concentration on the self-assembly of RNA virus particles**

Rees F. Garmann, Aaron M. Goldfain, Cheylene R. Tanimoto, Christian E. Beren, Fernando F. Vasquez, Charles M. Knobler, William M. Gelbart, Vinothan N. Manoharan

### Appendix A: Simulating the median absolute deviation (MAD) of start times measured with a diffusion-limited assembly pathway.

Here we simulate how the measured MAD of start times will scale with protein concentration in a diffusion-limited assembly pathway. We assume each particle accretes protein with exponential growth kinetics, as predicted by simple Langmuir adsorption, and that the characteristic growth time depends on protein concentration, $C$, as $\tau_{\mathrm{g}} = \tau_{\mathrm{g},0}\left(\frac{C_{\mathrm{ref}}}{C}\right)^{\alpha_{\mathrm{g}}}$. Here, $\tau_{\mathrm{g},0}$ is a characteristic growth time independent of protein concentration, $C_{\mathrm{ref}}$ is a reference concentration which we take as 1 μmol/L, and the exponent $\alpha_{\mathrm{g}}$ describes how the growth time scales with protein concentration. In Langmuir diffusion-limited models $\alpha_{\mathrm{g}} = 1$, but here we allow it to vary to explore how different exponents might affect our results. In a diffusion-limited pathway, protein accretion for different RNA strands occurs simultaneously and the assembly kinetics of each particle in an ensemble is identical. However, noise in our measurement will make each measured trace unique. We assume Gaussian noise, $\sigma(t)$, which is independent for each trace and each time point. Putting everything together, our model of the assembly traces, normalized to a maximum intensity of 1 is,

$$I(t) = \sigma(t) + \begin{cases} 0, & t < t_s \\ 1 - \exp\left[-\frac{t-t_s}{\tau_{\mathrm{g},0}}\left(\frac{C}{C_{\mathrm{ref}}}\right)^{\alpha_{\mathrm{g}}}\right], & t \geq t_s \end{cases}.$$

For our experiments, we quantify the spread in start times by fitting each measured trace to a piecewise exponential and determining the MAD of the fitted start times. Figure 6A shows how the MAD of start times depends on protein concentration. We simulate the concentration dependence of the MAD of start times by fitting an ensemble of simulated traces to the same piecewise exponential function and calculate the MAD of fitted start times at different concentrations. We perform simulations with $\alpha_{\mathrm{g}}$ values of 0.7 and 1 because these are the best fit value from our experiments and the expected value from Langmuir kinetics, respectively. We choose $\tau_{\mathrm{g},0}$ values of 12 s and 5.5 s for $\alpha_{\mathrm{g}}$ values of 0.7 and 1, such that the growth time for each concentration is similar to the experimentally measured median growth times. We choose a noise standard deviation of 0.05 relative to the maximum intensity to visually match variations in the measured traces. Figures S1A – S1C show all measured traces at 84 mmol/L NaCl along with an ensemble of five simulated traces. Figure S1D shows the MAD values extracted from fits to the simulated curves along with power-law fits for simulations with different protein concentrations. The fit power-law exponent from the simulations, $\alpha_{\mathrm{sim}}$, is 0.46 and 0.68 for $\alpha_{\mathrm{g}}$ values of 0.7 and 1, respectively. We note that the $\alpha_{\mathrm{sim}}$ does depend on the simulated noise magnitude (data not shown).

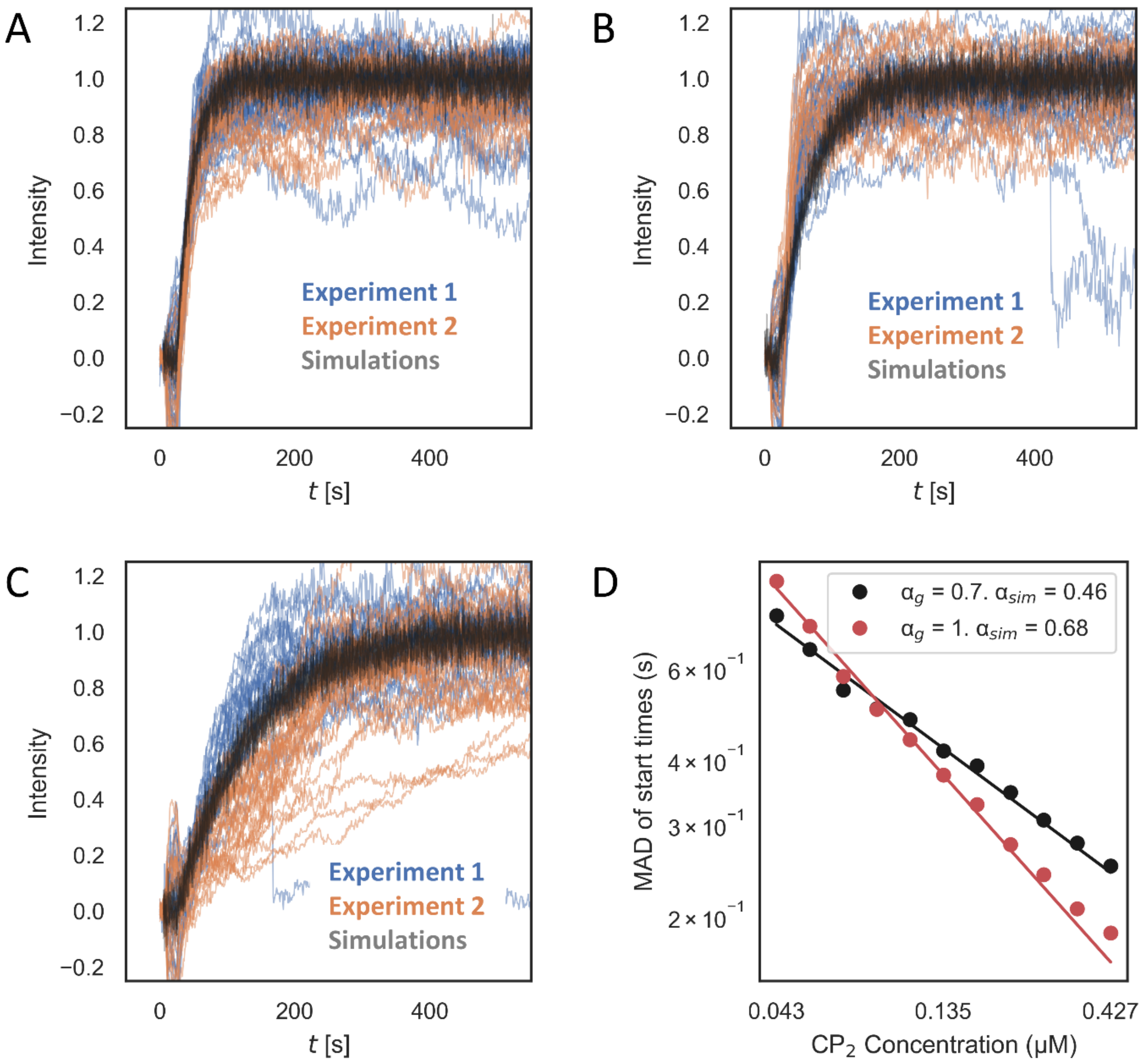


**Figure S1. Simulating the effects of measurement noise at 84 mmol/L NaCl with $\alpha_g$ = 0.7.** (A) Five simulated diffusion-limited growth traces and all measured growth traces from both experiments at 84 mmol/L NaCl and 0.426 µmol/L $CP_2$. (B) Five simulated diffusion limited growth traces and all measured growth traces from both experiments at 84 mmol/L NaCl and 0.135 µmol/L $CP_2$. (C) Five simulated diffusion limited growth traces and all measured growth traces from both experiments at 84 mmol/L NaCl and 0.043 µmol/L $CP_2$. In panels (A)-(C), the simulated traces are offset horizontally by about 50 s so they do not obscure the measured traces. (D) A log-log plot of the MAD of start times extracted from ensembles of simulated traces at various protein concentrations. 10,000 traces were simulated at each protein concentration. The lines are best fits to a power law function and $\alpha_{sim}$ is the negative of the exponent from the fit.

### Appendix B: The effects of non-instantaneous protein introduction.

Here we estimate how the non-instantaneous introduction of protein affects the measured start-time concentration-scaling exponent.

In our iSCAT experiments, we introduce protein solution into the sample chamber by pumping it in over the course of 20 seconds. The flow is at the low Reynold's number of 0.5, so we expect laminar flow within the chamber. Thus, with no slip boundary conditions, we expect a parabolic flow profile, as illustrated in Fig. S2A, top. After pumping in a protein solution with concentration $C_0$, we expect the protein solution to be centered in the chamber such that the protein must diffuse a distance to reach the chamber surface where the RNA is tethered. In our experiments, the field of view is approximately 9.3 mm from the chamber entrance, and given the volume of protein solution we introduce, the tip of the parabola would be approximately 45 mm beyond the chamber entrance. Thus, above the small, 10 µm by 10 µm iSCAT field of view, we expect the protein solution forms a nearly flat planar layer once pumping stops.

After pumping stops, the protein then diffuses to fill the chamber. To model how the protein diffuses to the surface-bound RNA, we solve the 1D diffusion equation. We assume the initial condition illustrated in Fig. S2A, bottom, where the protein and buffer solutions form infinite layers. The chamber has a height of $2L$ = 0.75 mm and the protein must diffuse a distance $L - a$ to reach the chamber surface. We believe the assumption of planar layers is justified because the protein solution is pumped well beyond the iSCAT field of view, and the 0.75-mm channel height is small compared to the distance the protein solution is pumped. We impose no-flux boundary conditions at the chamber walls and solve the 1D diffusion equation to find the protein concentration at the surface as a function of time. The concentration of protein at the surface is

$$C(t) = C_0 \frac{a}{L} + \frac{2C_0}{\pi} \sum_{n=1}^{\infty} \frac{(-1)^n}{n} \sin\left(\frac{n\pi a}{L}\right) e^{-n^2\pi^2 t/t_D},$$

where $t_D = L^2/D$ is a characteristic diffusion time and $D$ is the diffusion coefficient of a protein dimer. For BMV protein dimers, we assume $D$ = 91 µm$^2$/s based on measurements[1] of the hydrodynamic radius of MS2 $CP_2$ (2.5 nm).

Fig. S2B plots the protein concentration at the surface relative to $C_0$ for different distances that the protein must diffuse, assuming that it starts diffusing at $t$ = 20 s. Assuming parabolic flow and given the volume of protein we flow into the chamber, we calculate that the distance the protein must diffuse to reach the surface is 40 µm ($a/L$ = 0.89). We experimentally verify this distance using bright-field microscopy to image a solution of 1 µm-diameter polystyrene particles that we flow into the imaging chamber. With bright-field microscopy, we find that the distance protein must diffuse ranges from 20 µm to 50 µm ($a/L$ = 0.95 to 0.87) which is consistent with that calculated based on parabolic flow. Based on the diffusion model, the time required for the protein concentration at the surface to reach half of the bulk protein concentration ranges from 6 s to 39 s.

To model how protein diffusion affects the measured nucleation kinetics, we couple the protein diffusion model to a classical nucleation model. In the classical nucleation model, $\frac{df}{dt} = KC_0^{\alpha_\mathrm{n}}(1-f)$, where $f$ is the faction of RNA strands on which a nucleus has formed, $K$ is a rate constant determining the nucleation rate, and the exponent $\alpha_\mathrm{n}$ describes how the nucleation time scales with protein concentration. By the law of mass action, $\alpha_\mathrm{n}$ is the number of proteins in the critical nucleus, so we expect $\alpha_\mathrm{n} > 1$. The solution to this differential equation is an exponential distribution of nucleation times. We extend this classical nucleation model by allowing the protein concentration to vary with time, as determined by our diffusion model. The differential equation becomes $\frac{df}{dt} = KC_0^{\alpha_\mathrm{n}} c(t)^{\alpha_\mathrm{n}}(1-f)$, where $c(t) = C(t)/C_0$. This can be solved, yielding

$$f(t) = \begin{cases} 0, & t < t_0 \\ 1 - \exp\left[-\left(\frac{C_0}{C_{\text{ref}}}\right)^{\alpha_\text{n}} \frac{t^*}{\tau_{\text{n},0}}\right], & t \geq t_0 \end{cases}$$

where $\tau_{\text{n},0} = 1/KC_{\text{ref}}^{\alpha}$ is the characteristic nucleation time, $t^* = \int_{t_0}^{t} c(t' - t_0)^{\alpha_\text{n}}\, dt'$, and $t_0$ is the time at which protein starts to diffuse. Fig. S2C plots $f(t)$ for varying values of $a/L$ with $\alpha_\text{n} = 2$, $\tau_{\text{n},0}$ = 50 s, $t_0$ = 20 s, and $C_0$ = 1 μmol/L . Note that if $a/L = 1$, that is, the protein solution instantaneously fills the entire chamber, $t^* = t - t_0$, and classical exponential nucleation kinetics are recovered.

Our goal is to use the model of nucleation kinetics to determine how protein diffusion affects the experimentally measured nucleation concentration scaling exponent. To do this, we perform a least-squares fit on measured cumulative distributions of start times. We simultaneously fit the cumulative distributions from all experiments at a given salt concentration to determine the best fit to $\alpha_\text{n}$ at each salt concentration. At both 84 mmol/L and 167 mmol/L salt, we perform 6 experiments each (two experiments at three protein concentrations). At each salt concentration, there are 14 fitting parameters: $\alpha_\text{n}$, $\tau_{\text{n},0}$, six values of the ratio $a/L$ (one for each experiment), and six values of $t_0$ (one for each experiment). Figs. S2D and S2E show the best fit to our measurements at 84 mmol/L NaCl and 167 mmol/L NaCl, respectively.

The best fit value of $\alpha_\text{n}$ depends strongly on bounds limiting the fit value of the ratio $a/L$. With lower bound on $a/L$, at 84 mmol/L NaCl, the fit converges to $\alpha_\text{n}$ = 3.3 ± 1.4, as indicated in Fig. S2F (the uncertainty is from the covariance of the fit). However, two experiments have fit values of $a/L$ significantly below 0.87, the smallest expected value of $a/L$ (Fig. S2F). Similarly, at 167 mmol/L NaCl, the fit converges to $\alpha_\text{n}$ = 2.8 ± 0.4, but again, two experiments have fit values of $a/L$ below 0.87. If a lower bound is placed on $a/L$, the fit value of $\alpha_\text{n}$ decreases. Thus, we use our fits with no lower limit on $a/L$ to set an upper bound on $\alpha_\text{n}$, and we perform additional fits where we limit $a/L$ to be no less than 0.87 to set a lower bound on $\alpha_\text{n}$. The lower bound on $\alpha_\text{n}$, as determined from these fits is $\alpha_\text{n}$ = 1.2 ± 0.4 at 84 mmol/L NaCl, and $\alpha_\text{n}$ = 1.5 ± 0.2 at 167 mmol/L NaCl. We combine the upper and lower bounds, to arrive at the values of $\alpha_\text{n}$ mentioned in the Discussion section of the main text, $\alpha_\text{n}$ = 2.8 ± 2.0 at 84 mmol/L NaCl, and $\alpha_\text{n}$ = 2.3 ± 1.0 at 167 mmol/L NaCl.

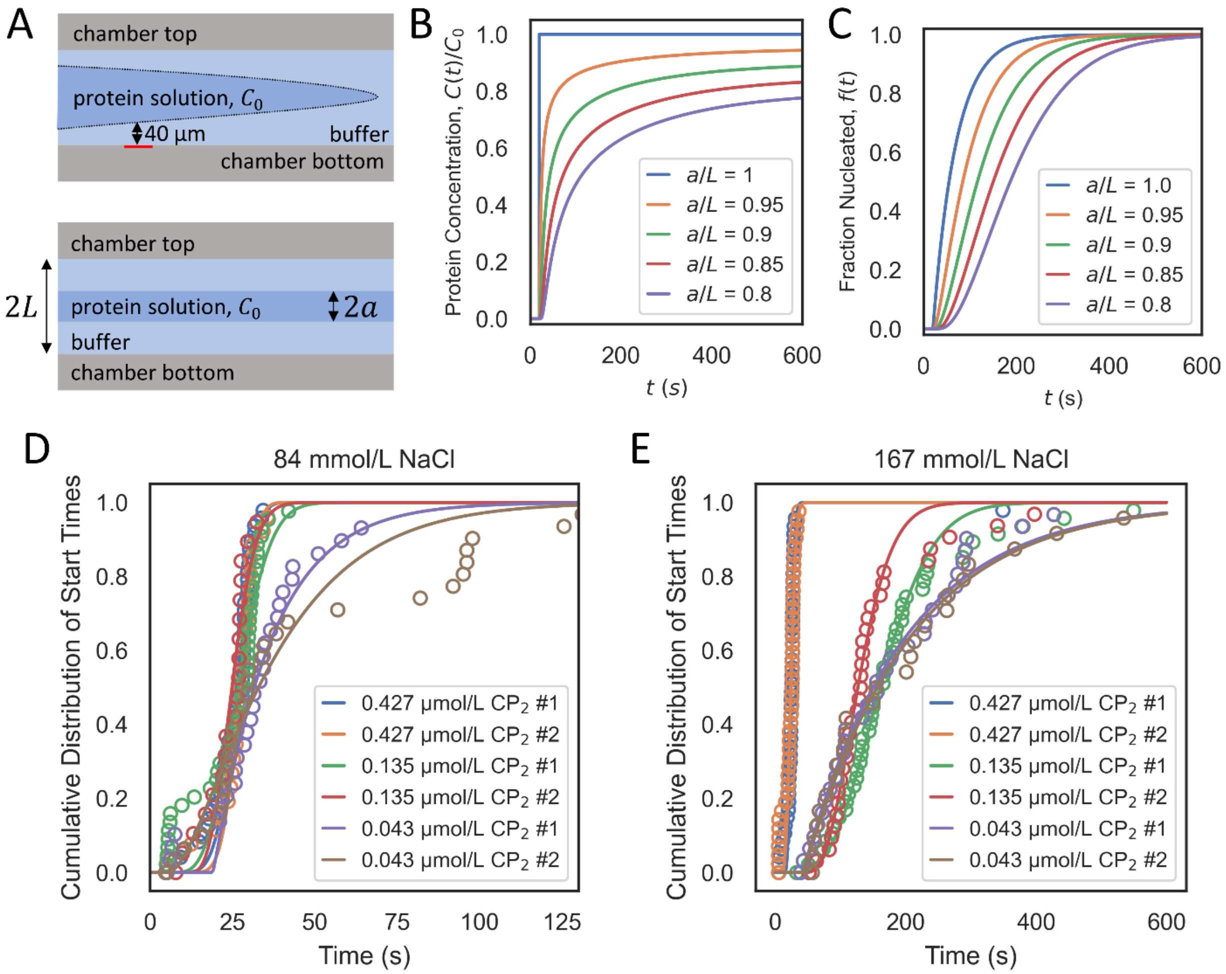


F

| | $\alpha_n$ | $a/L$ 0.427 µmol/L #1 | $a/L$ 0.427 µmol/L #2 | $a/L$ 0.135 µmol/L #1 | $a/L$ 0.135 µmol/L #2 | $a/L$ 0.043 µmol/L #1 | $a/L$ 0.043 µmol/L #2 |
|---|---|---|---|---|---|---|---|
| **84 mmol/L NaCl Best Fit** | 3.3 | 0.73 | 0.75 | 0.84 | 0.88 | 1.0 | 0.96 |
| **84 mmol/L NaCl Uncertainty** | 1.4 | 0.06 | 0.06 | 0.04 | 0.03 | 0.02 | 0.03 |
| **167 mmol/L NaCl Best Fit** | 2.8 | 0.88 | 0.84 | 0.63 | 0.75 | 1.0 | 1.0 |
| **167 mmol/L NaCl Uncertainty** | 0.4 | 0.03 | 0.03 | 0.04 | 0.04 | 0.02 | 0.02 |

**Figure S2. Modeling how the non-instantaneous introduction of protein affects the measured nucleation concentration exponent.** (A) Top: A cartoon of parabolic flow in our sample chambers. The red bar represents the location of the iSCAT field of view. Bottom: Initial condition used when solving the diffusion equation. (B) Modelled protein concentration as a function of time at the surface-bound RNA strands. The concentration of protein at the surface gradually increases as proteins diffuse in the sample chamber. (C) Modelled nucleation kinetics where the protein concentration gradually increases with time as determined by the diffusion model. (D) Cumulative distributions of measured start times for all 6 experiments at 84 mmol/L NaCl (circles) and best fit to the modelled nucleation kinetics including protein diffusion (lines). There are two experiments at each protein concentration. (E) Same as (D) but at 167 mmol/L NaCl. (F) Best fit parameters for the fits shown in parts (D) and (E). The fit nucleation scaling exponent, $\alpha_n$, and the ratio $a/L$ representing how far protein diffused to the surface are given along with their uncertainties. The uncertainties are determined from the covariance of the fit.

**Figure S3. Plots of all assembly traces and fits.** In the following pages, all of the assembly traces analyzed in the main text are plotted in various colors, which designate the experimental conditions used, and fitted with a piecewise function, which is shown in red. As described in the main text, the piecewise function has the form $I(t) = \begin{cases} I_0, \ t < t_s \\ I_0 + I_f\left(1 - \exp\left[-(t - t_s)/\tau_g\right]\right), \ t \geq t_s \end{cases}$, where *I(t)* is the intensity as a function of time, $I_0$ is the initial intensity (which can be offset from zero because of microscope drift), $I_f$ is the final intensity, $t_s$ is the start time – the time at which the trace begins to rise from its initial value, and $\tau_g$ is the growth time—the time for the trace to reach (1-1/*e*) = 0.63 of its final value once it has started increasing. The gray bar to the right of each trace shows the size of a full BMV capsid.

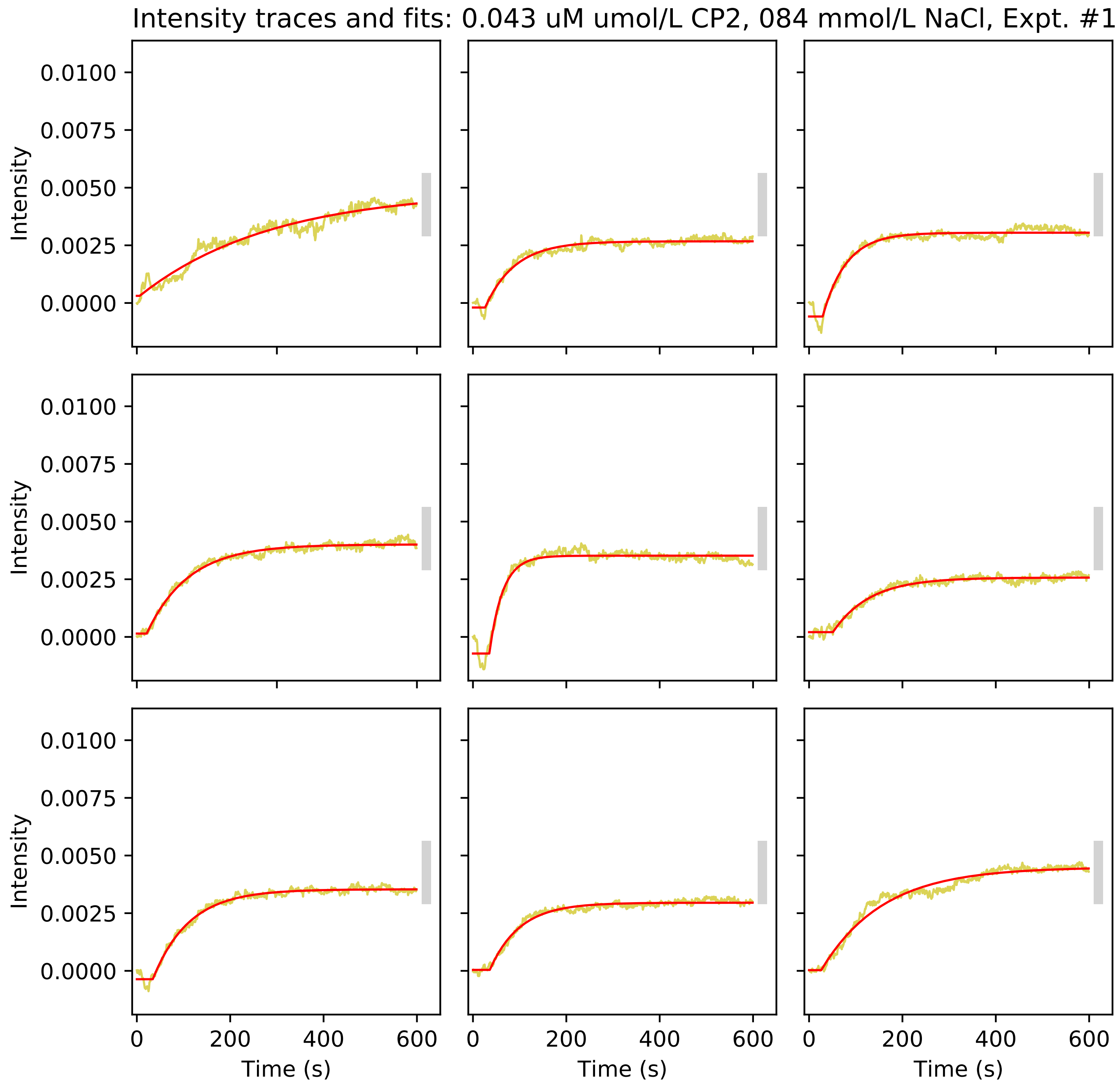
Intensity traces and fits: 0.043 uM umol/L CP2, 084 mmol/L NaCl, Expt. #1
Intensity
0.0100
0.0075
0.0050
0.0025
0.0000
0
200
400
600
Time (s)

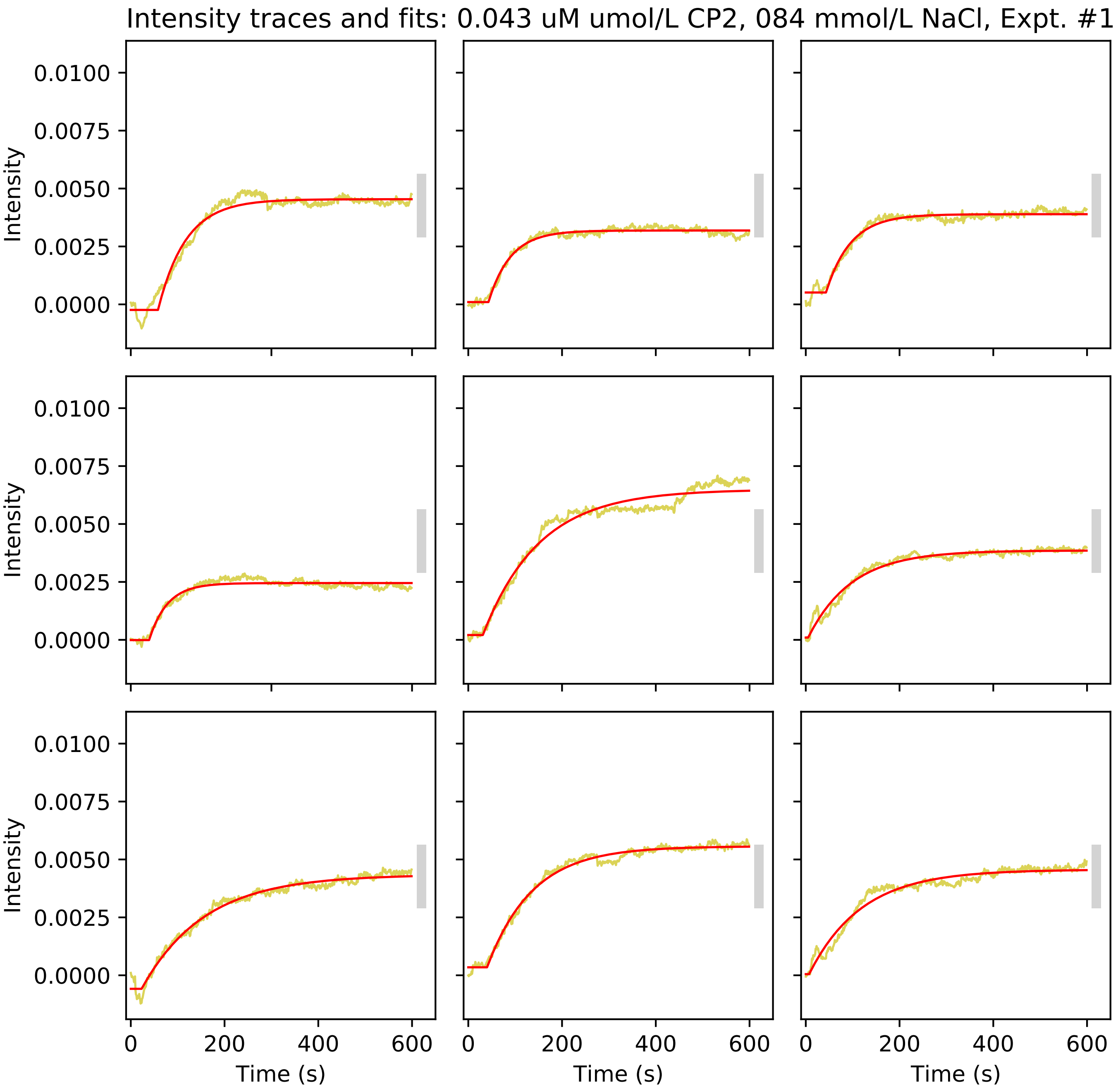
Intensity traces and fits: 0.043 uM umol/L CP2, 084 mmol/L NaCl, Expt. #1
Intensity
0.0100
0.0075
0.0050
0.0025
0.0000
0
200
400
600
Time (s)

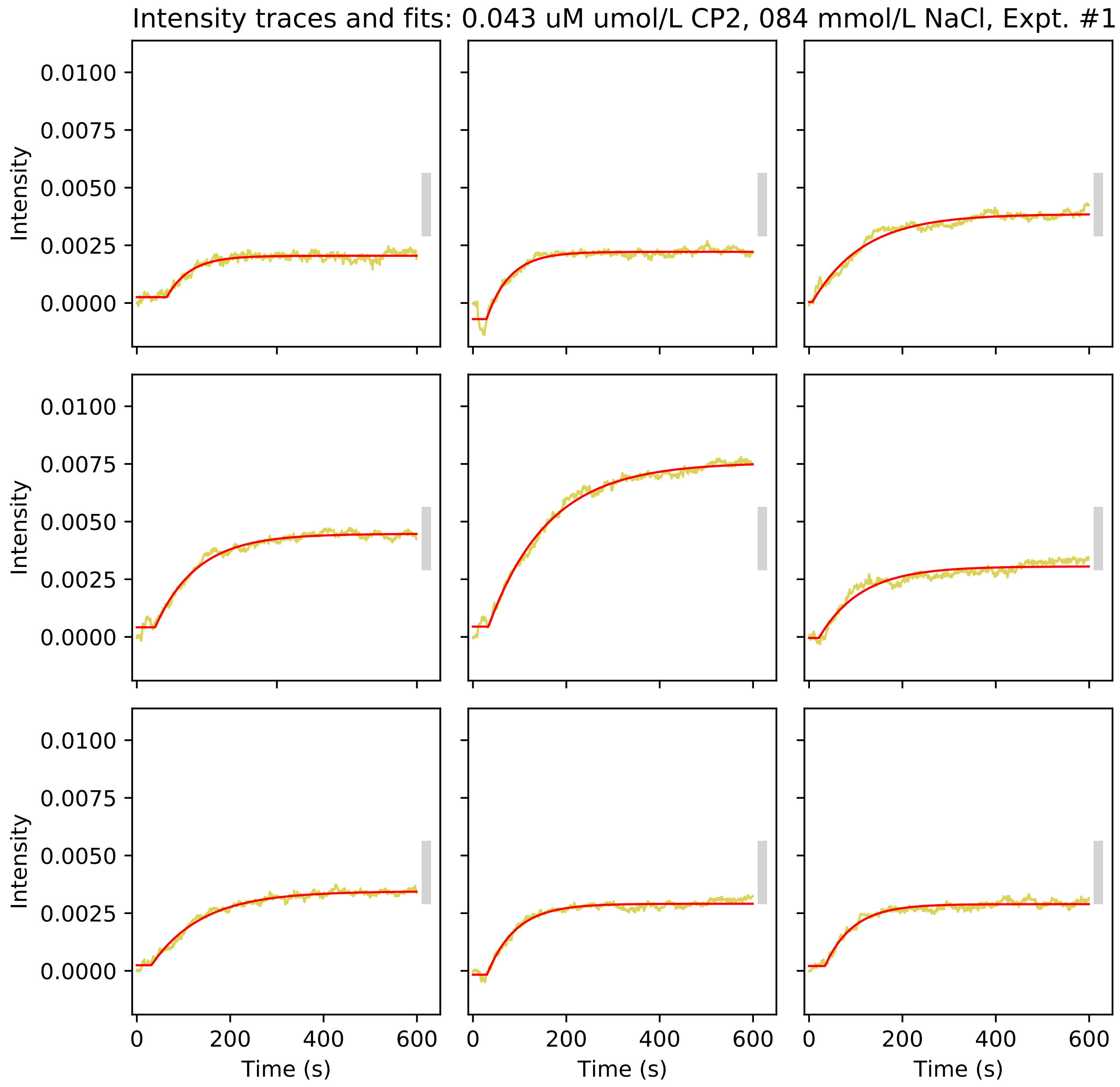
Intensity traces and fits: 0.043 uM umol/L CP2, 084 mmol/L NaCl, Expt. #1
Intensity
0.0100
0.0075
0.0050
0.0025
0.0000
0
200
400
600
Time (s)

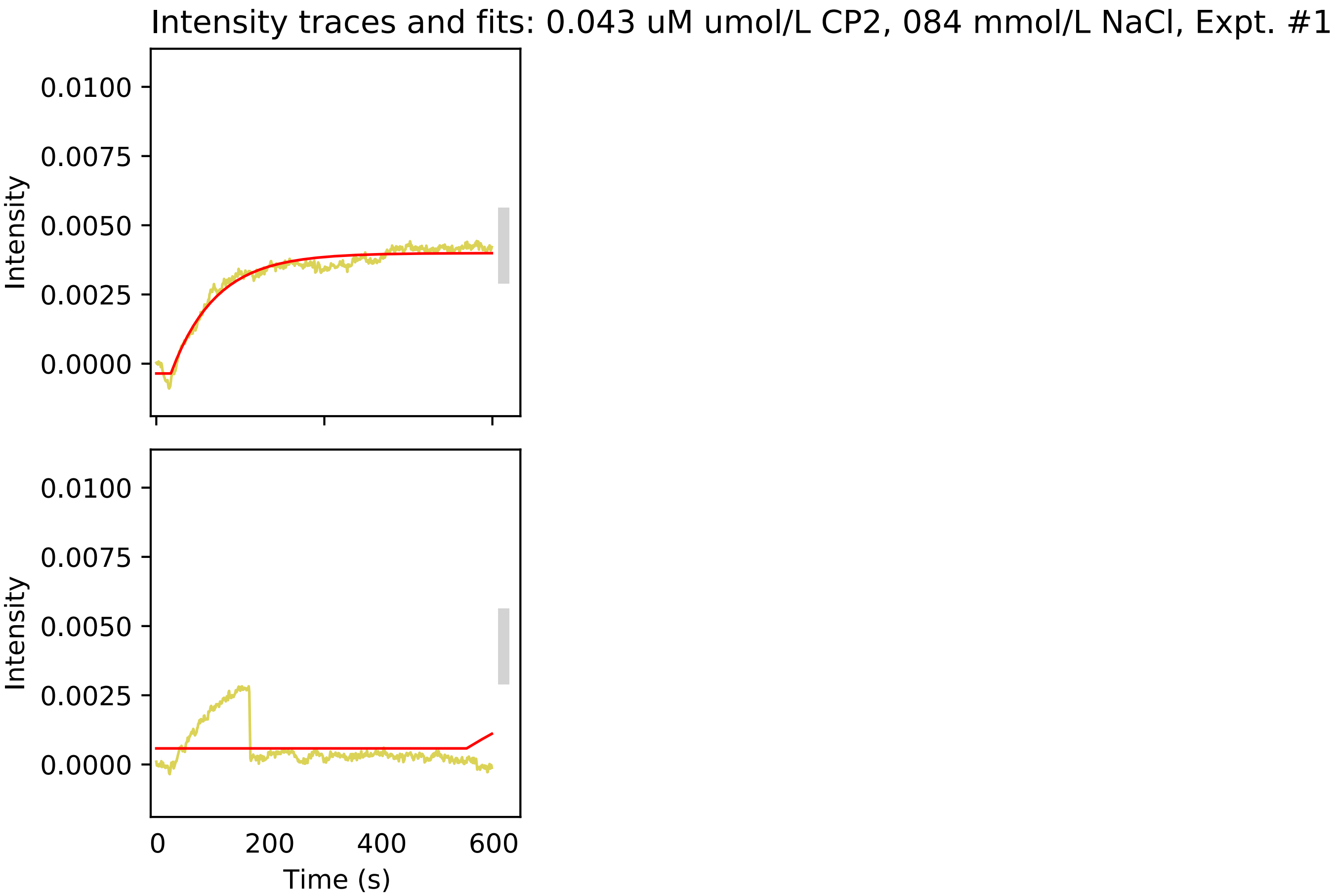
Intensity traces and fits: 0.043 uM umol/L CP2, 084 mmol/L NaCl, Expt. #1
Intensity
0.0100
0.0075
0.0050
0.0025
0.0000
Intensity
0.0100
0.0075
0.0050
0.0025
0.0000
0
200
400
600
Time (s)

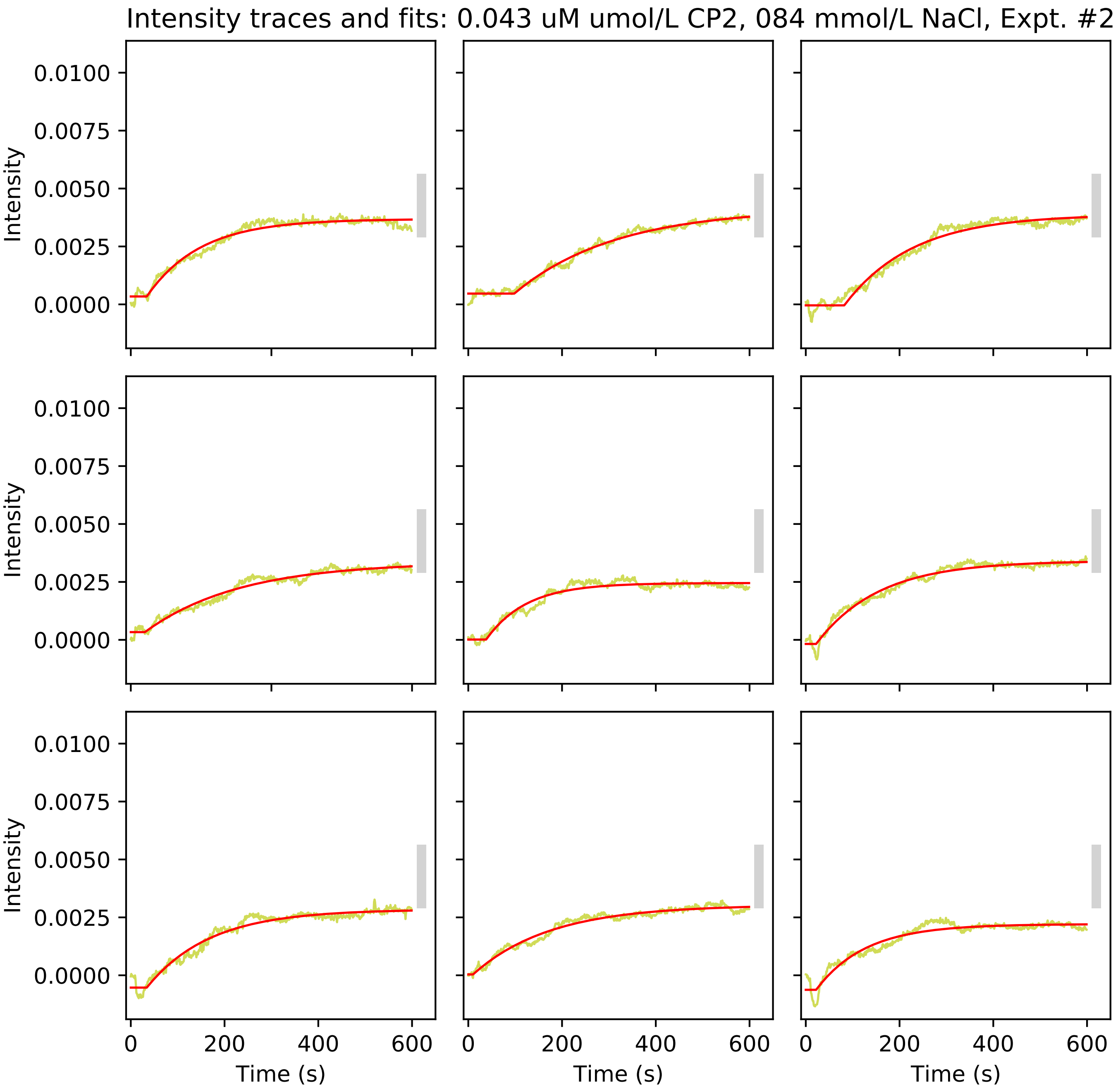

Intensity traces and fits: 0.043 uM umol/L CP2, 084 mmol/L NaCl, Expt. #2
Intensity
0.0100
0.0075
0.0050
0.0025
0.0000
Time (s)
0
200
400
600

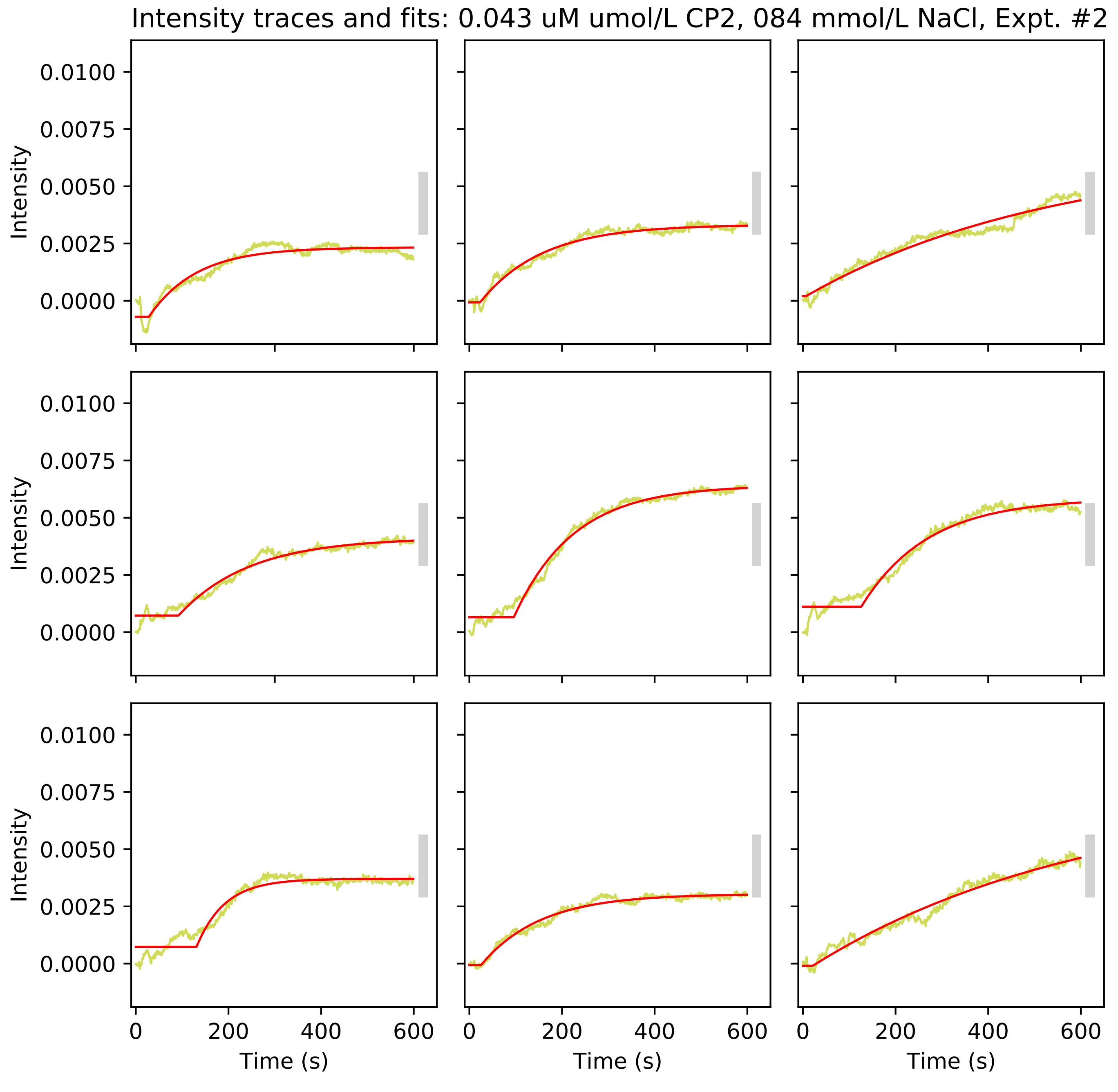

Intensity traces and fits: 0.043 uM umol/L CP2, 084 mmol/L NaCl, Expt. #2
Intensity
0.0100
0.0075
0.0050
0.0025
0.0000
0
200
400
600
Time (s)

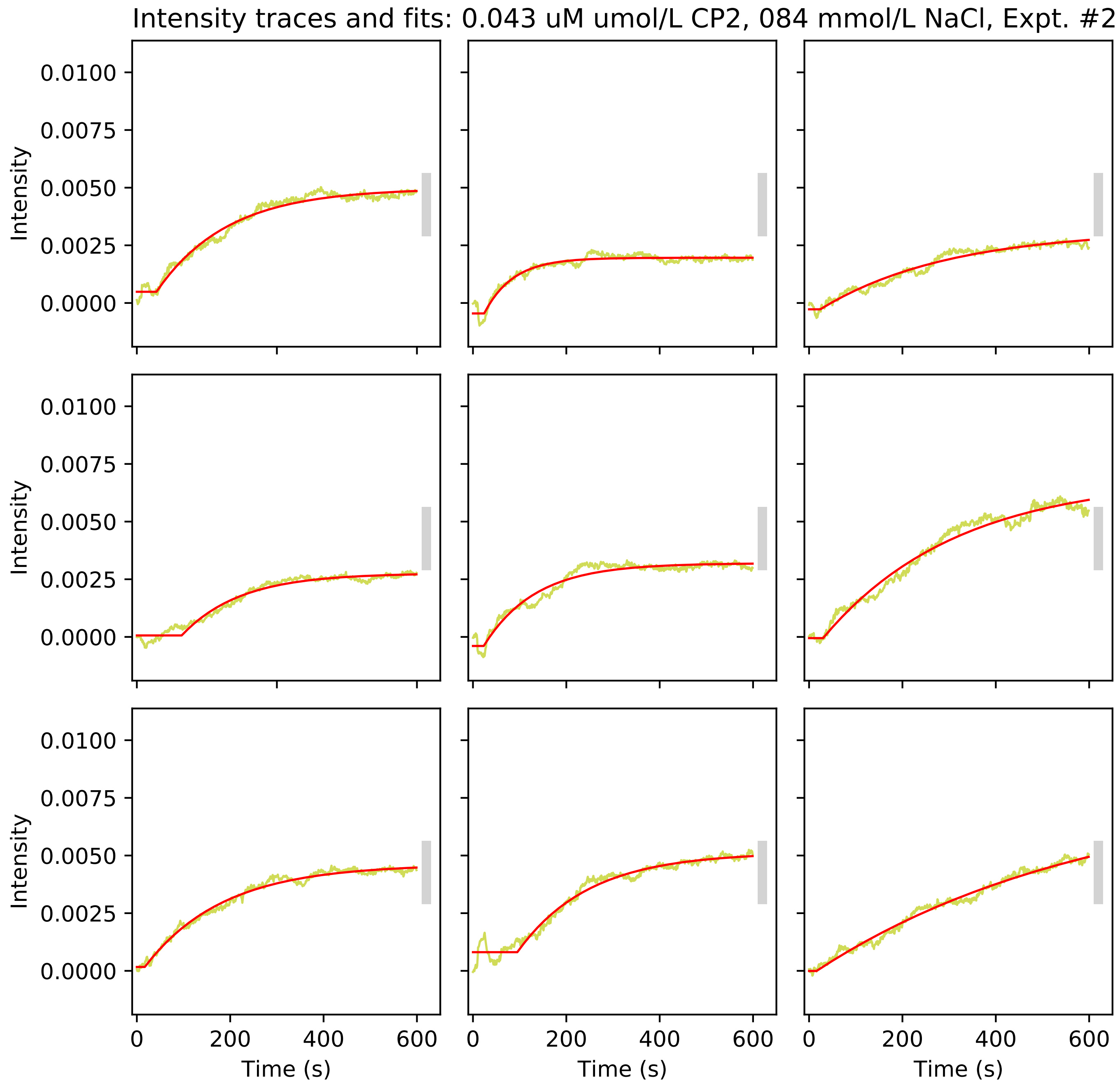
Intensity traces and fits: 0.043 uM umol/L CP2, 084 mmol/L NaCl, Expt. #2
Intensity
0.0100
0.0075
0.0050
0.0025
0.0000
0
200
400
600
Time (s)

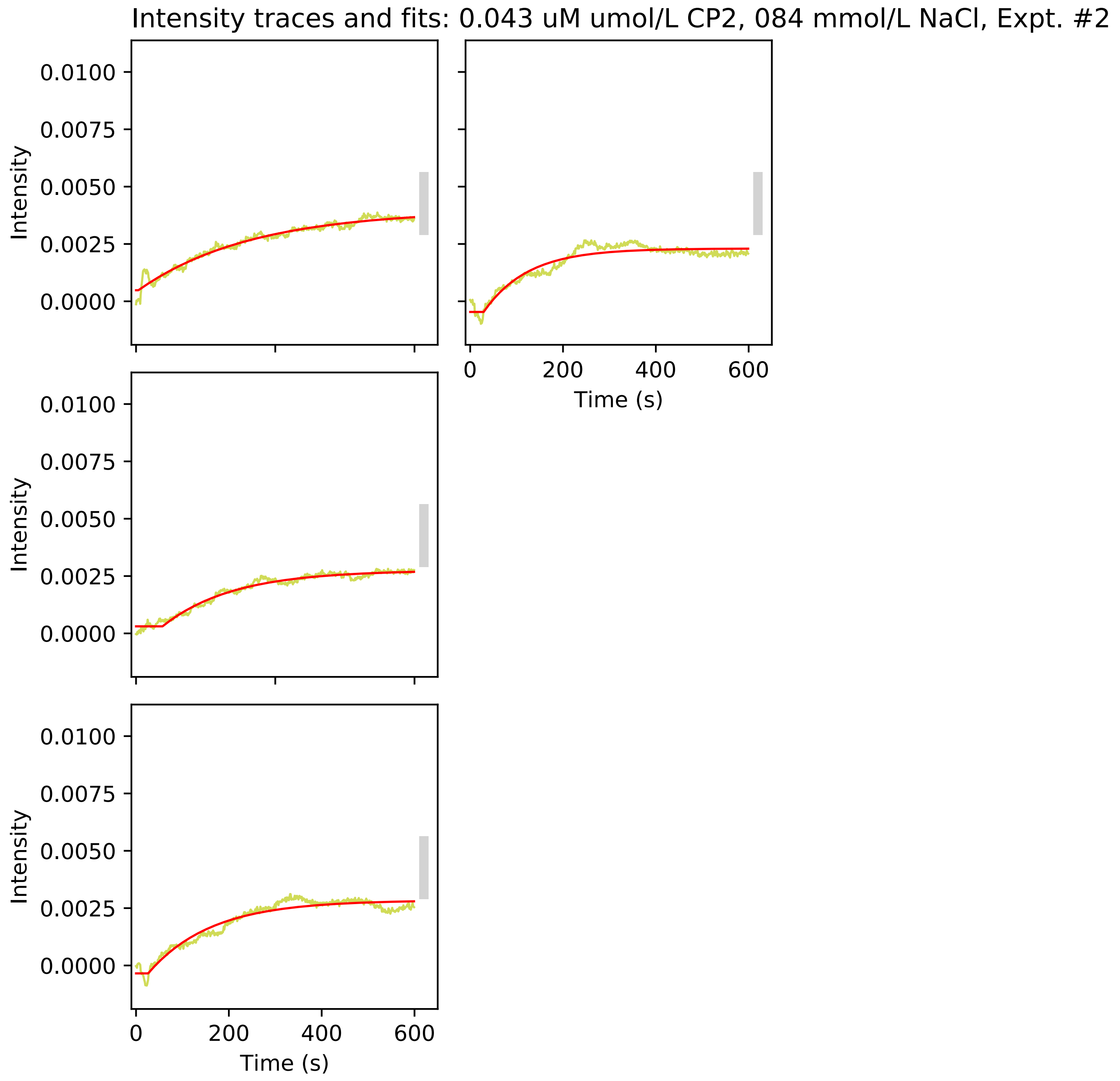
Intensity traces and fits: 0.043 uM umol/L CP2, 084 mmol/L NaCl, Expt. #2
Intensity
0.0100
0.0075
0.0050
0.0025
0.0000
Time (s)
0
200
400
600

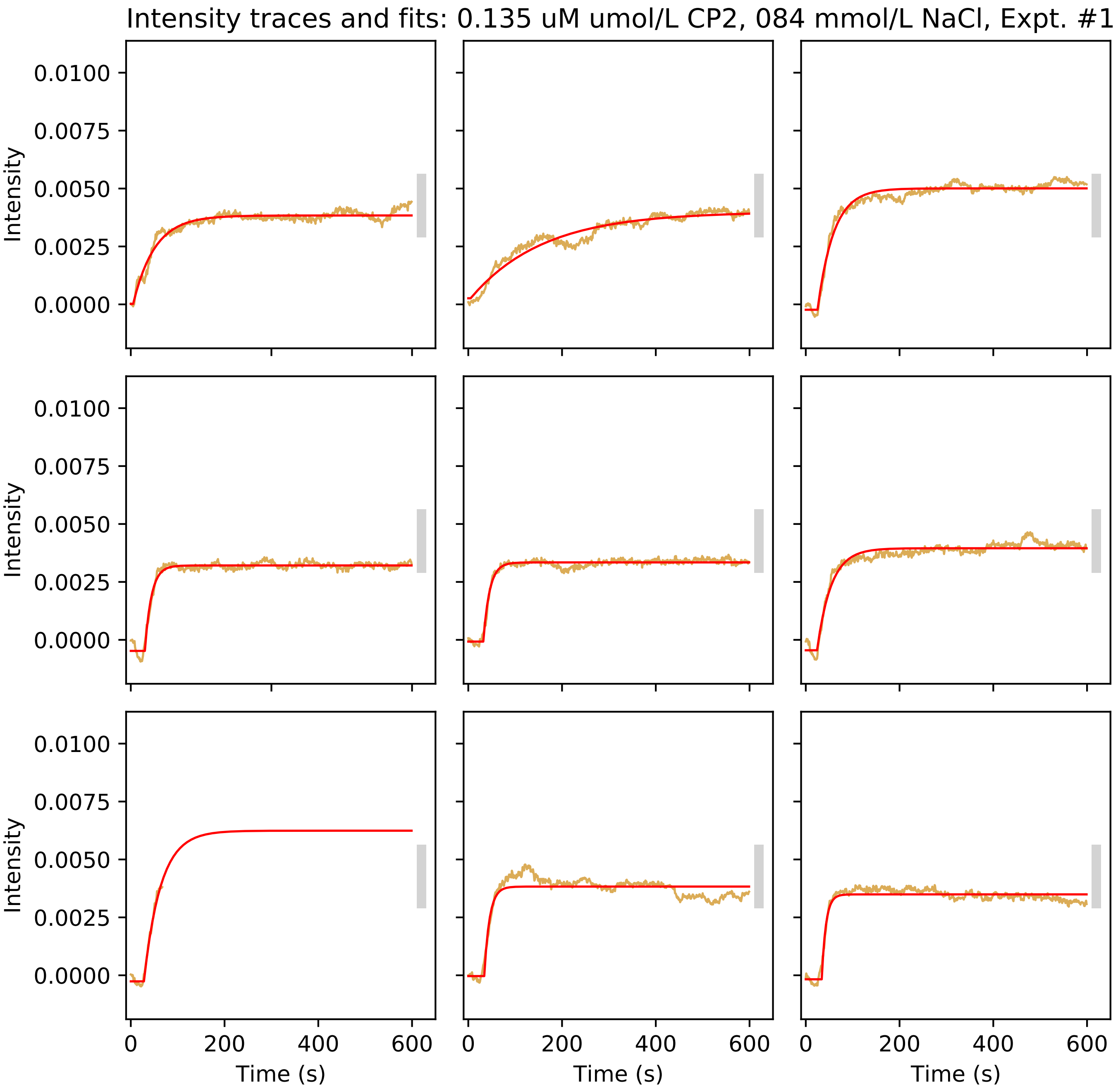
Intensity traces and fits: 0.135 uM umol/L CP2, 084 mmol/L NaCl, Expt. #1
Intensity
0.0100
0.0075
0.0050
0.0025
0.0000
0
200
400
600
Time (s)

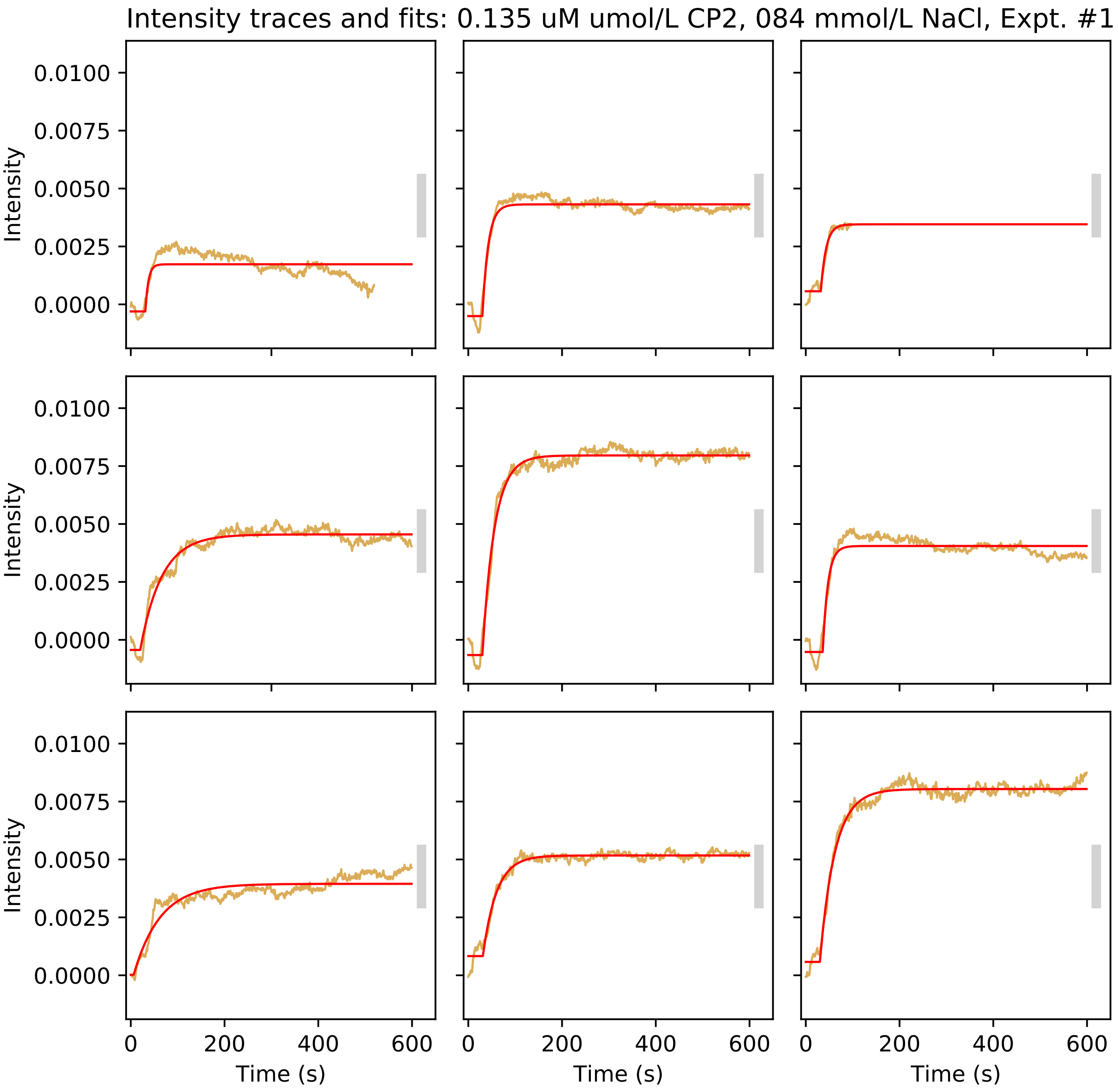
Intensity traces and fits: 0.135 uM umol/L CP2, 084 mmol/L NaCl, Expt. #1
Intensity
0.0100
0.0075
0.0050
0.0025
0.0000
Time (s)
0
200
400
600

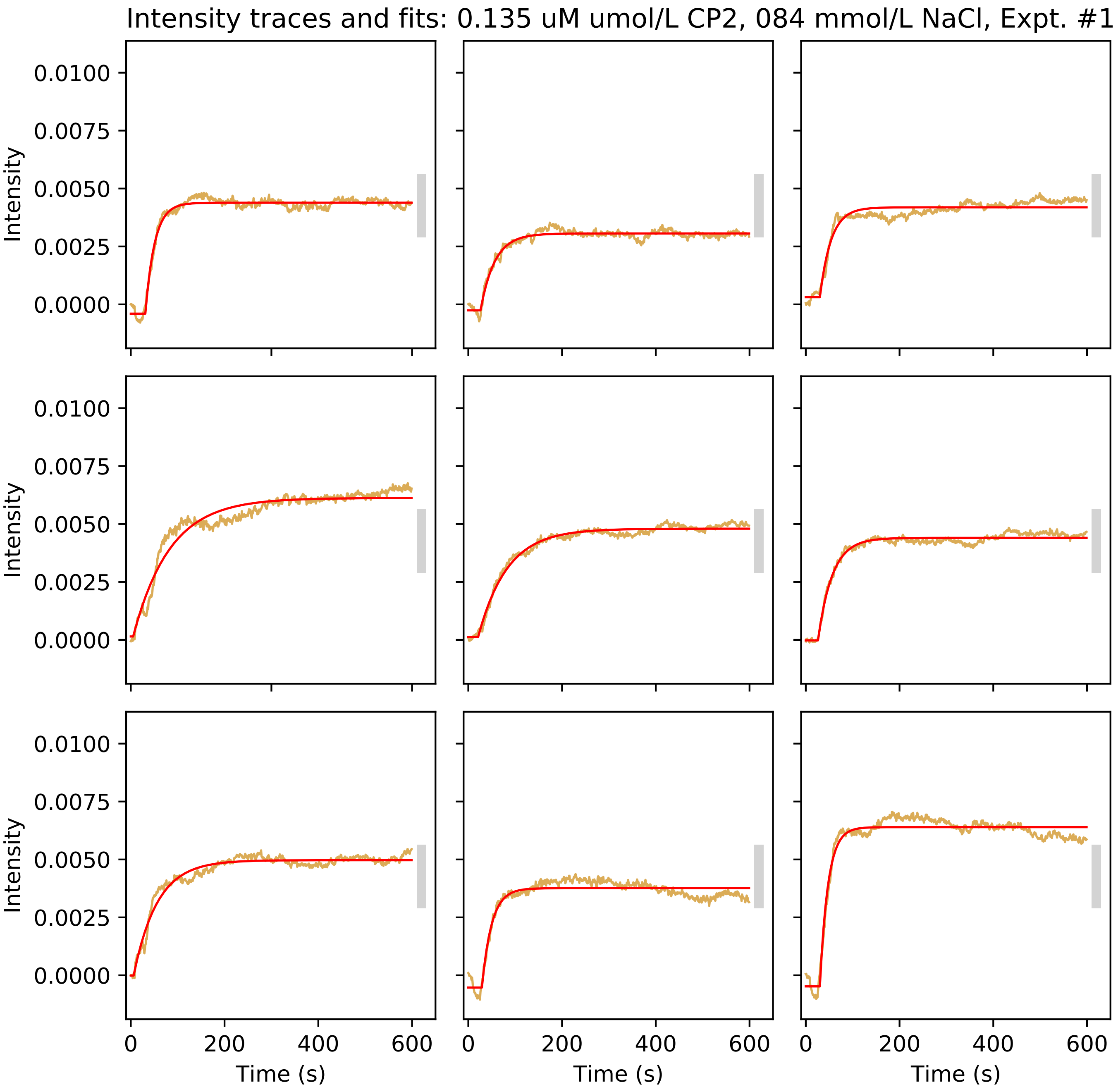
Intensity traces and fits: 0.135 uM umol/L CP2, 084 mmol/L NaCl, Expt. #1
Intensity
0.0100
0.0075
0.0050
0.0025
0.0000
0
200
400
600
Time (s)

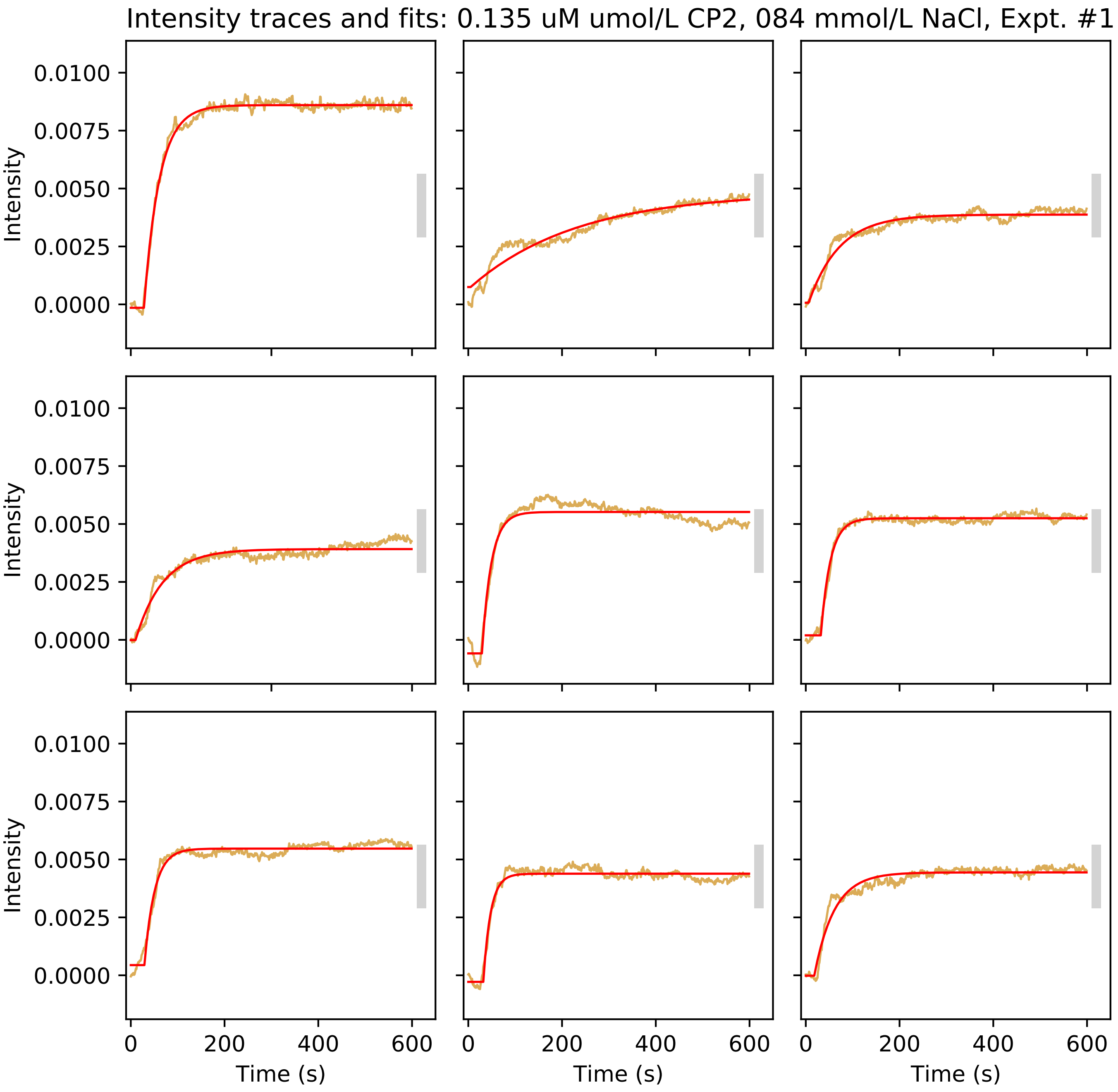
Intensity traces and fits: 0.135 uM umol/L CP2, 084 mmol/L NaCl, Expt. #1
Intensity
Time (s)
0.0100
0.0075
0.0050
0.0025
0.0000
0
200
400
600

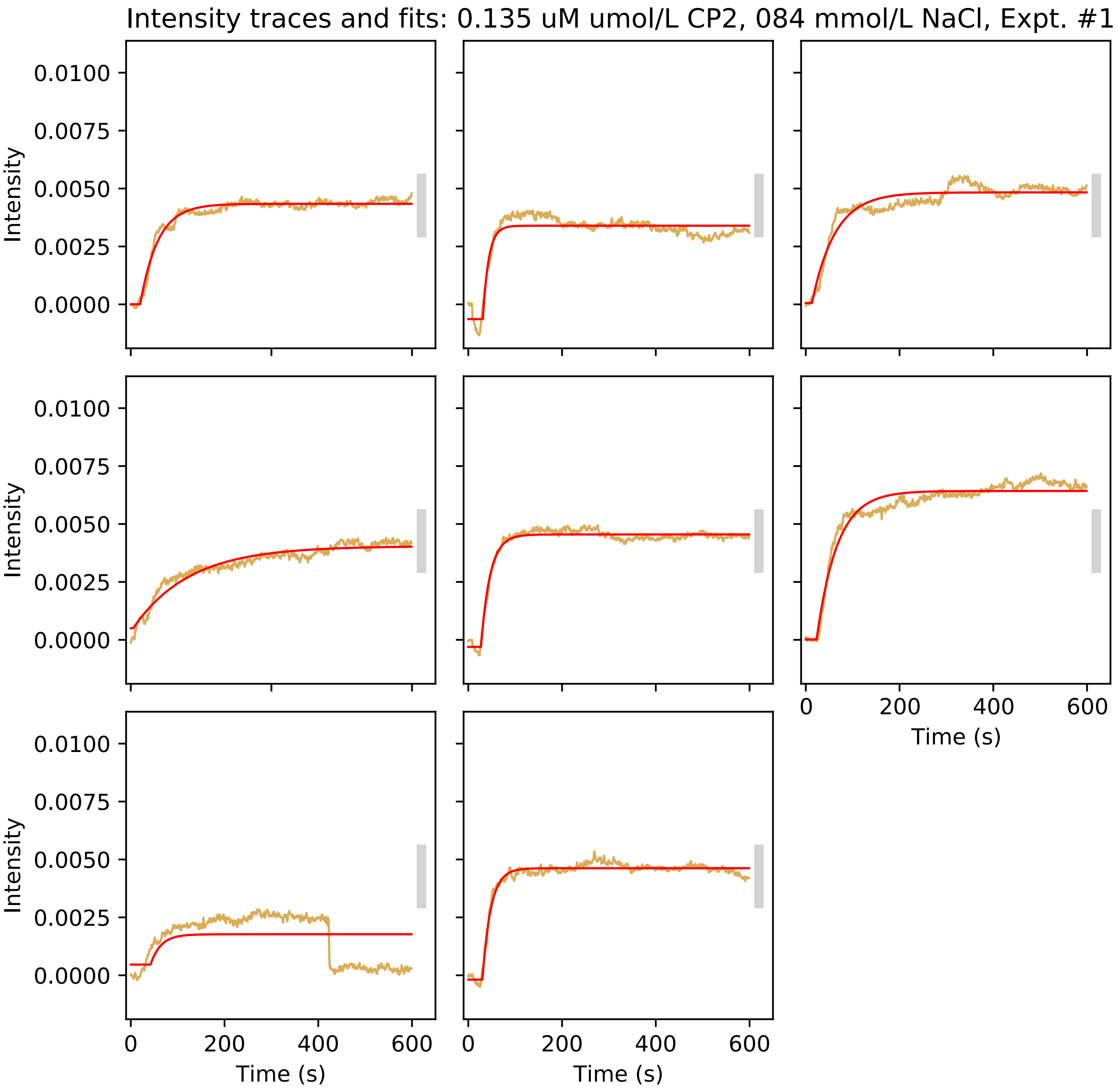

Intensity traces and fits: 0.135 uM umol/L CP2, 084 mmol/L NaCl, Expt. #1
Intensity
0.0100
0.0075
0.0050
0.0025
0.0000
0
200
400
600
Time (s)

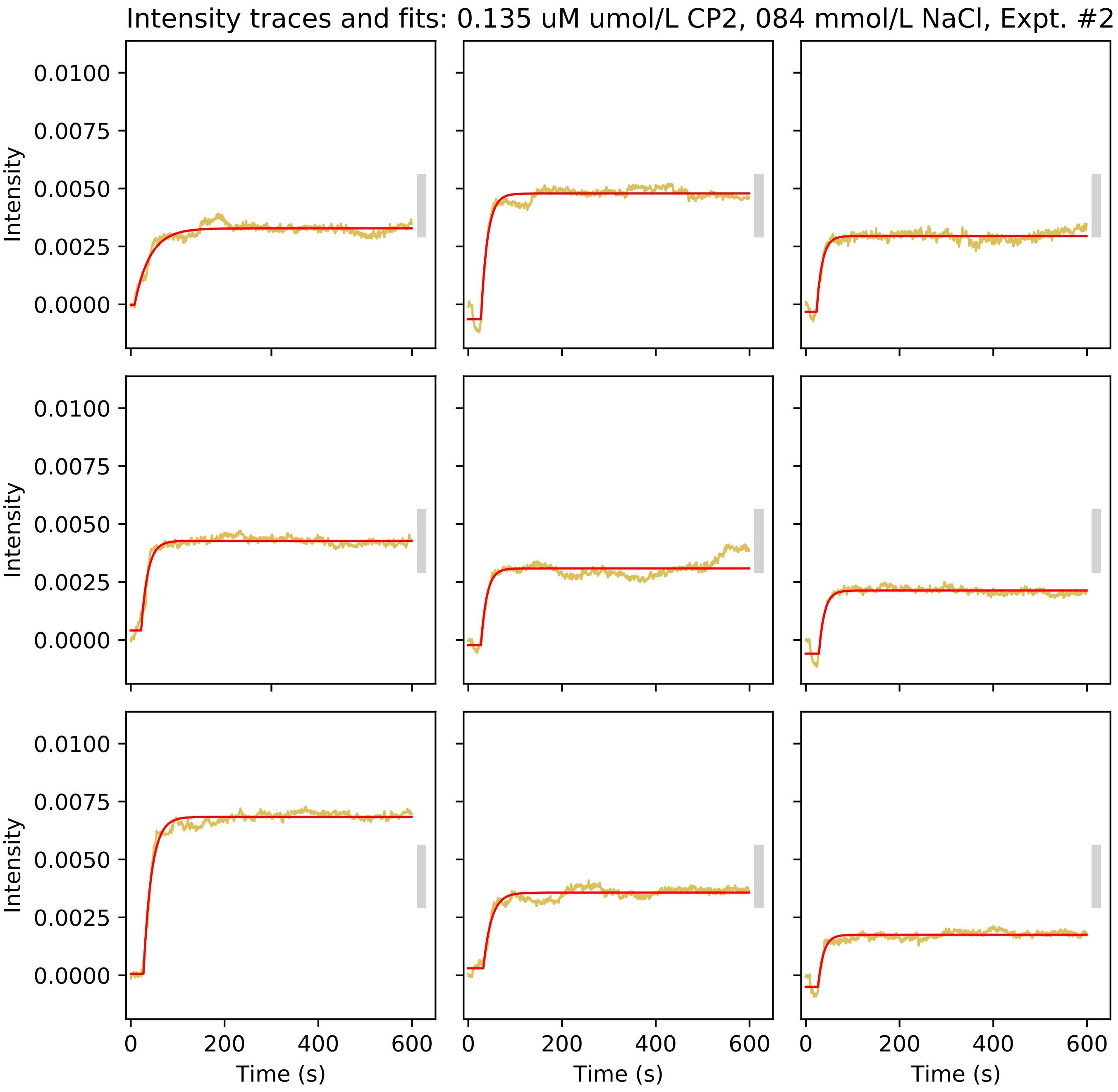
Intensity traces and fits: 0.135 uM umol/L CP2, 084 mmol/L NaCl, Expt. #2
Intensity
0.0100
0.0075
0.0050
0.0025
0.0000
Time (s)
0
200
400
600

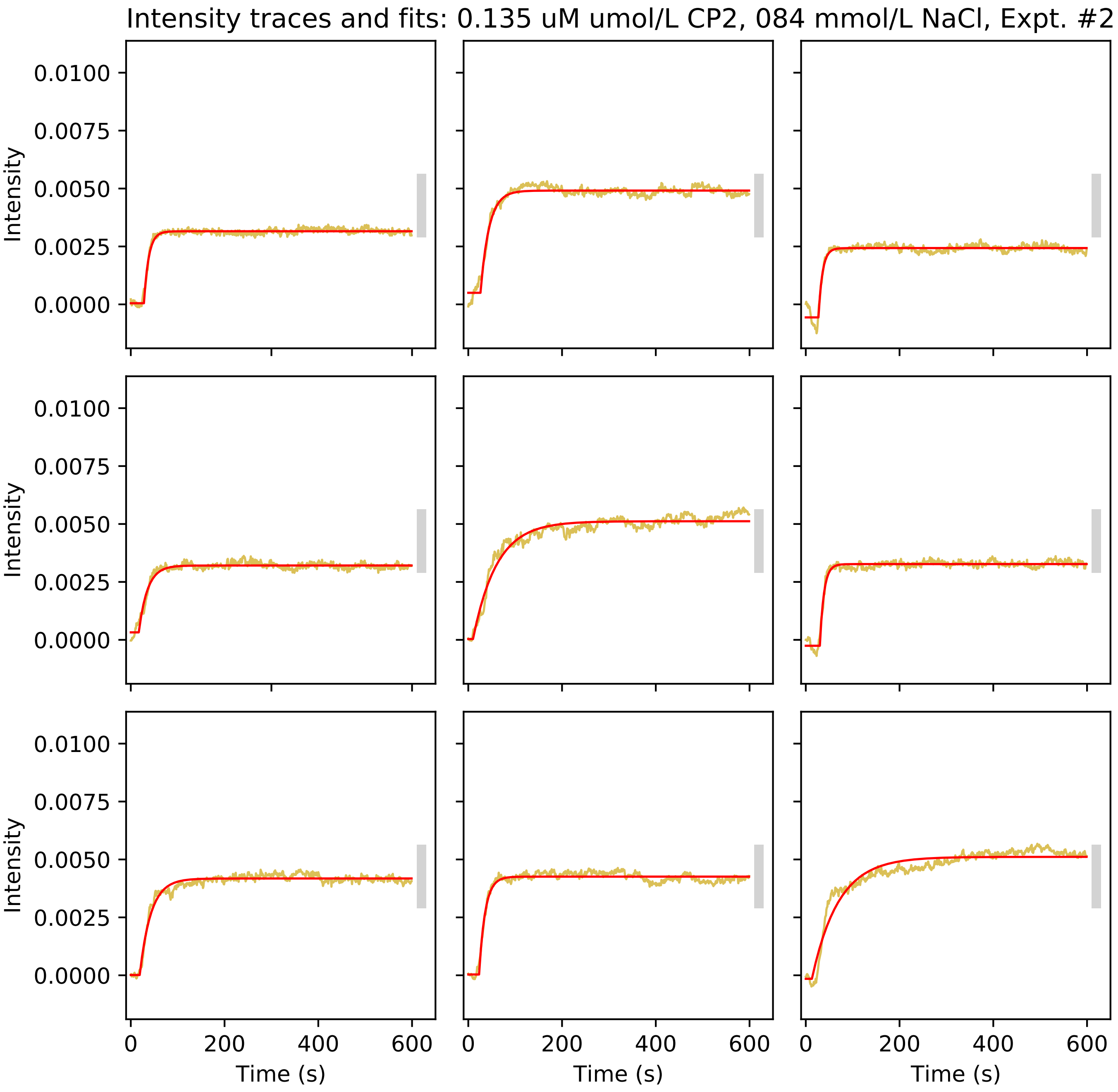
Intensity traces and fits: 0.135 uM umol/L CP2, 084 mmol/L NaCl, Expt. #2
Intensity
0.0100
0.0075
0.0050
0.0025
0.0000
0
200
400
600
Time (s)

Intensity traces and fits: 0.135 uM umol/L CP2, 084 mmol/L NaCl, Expt. #2

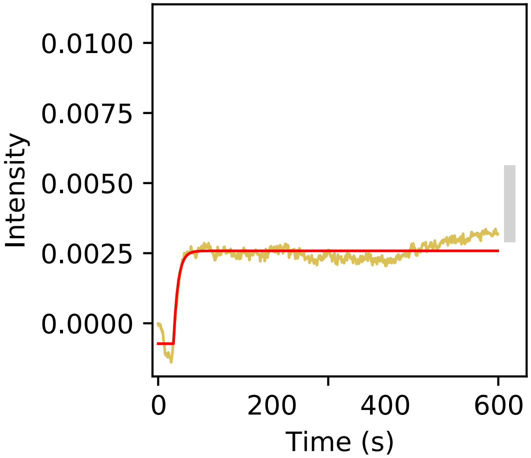

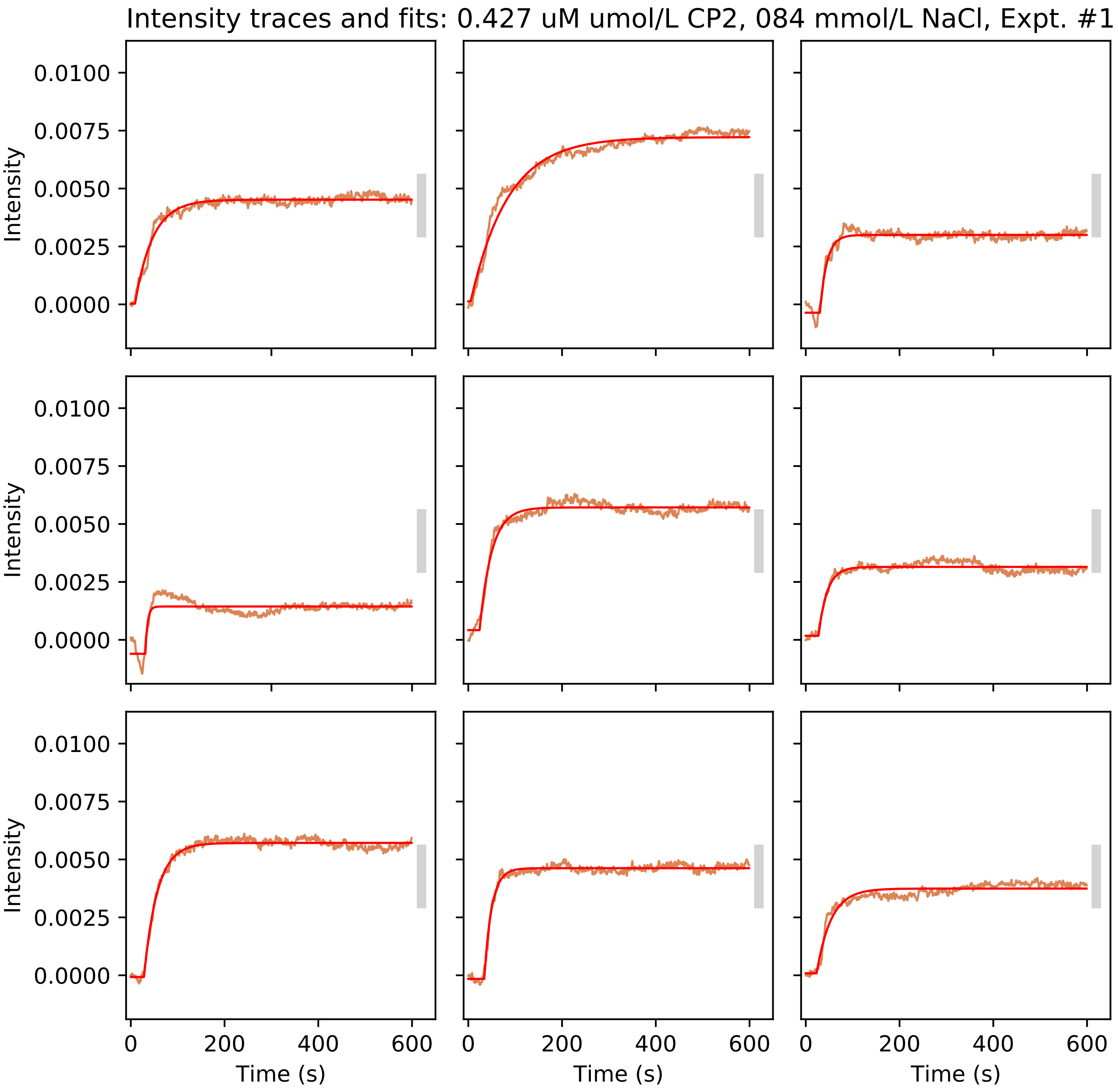
Intensity traces and fits: 0.427 uM umol/L CP2, 084 mmol/L NaCl, Expt. #1
Intensity
0.0100
0.0075
0.0050
0.0025
0.0000
Time (s)
0
200
400
600

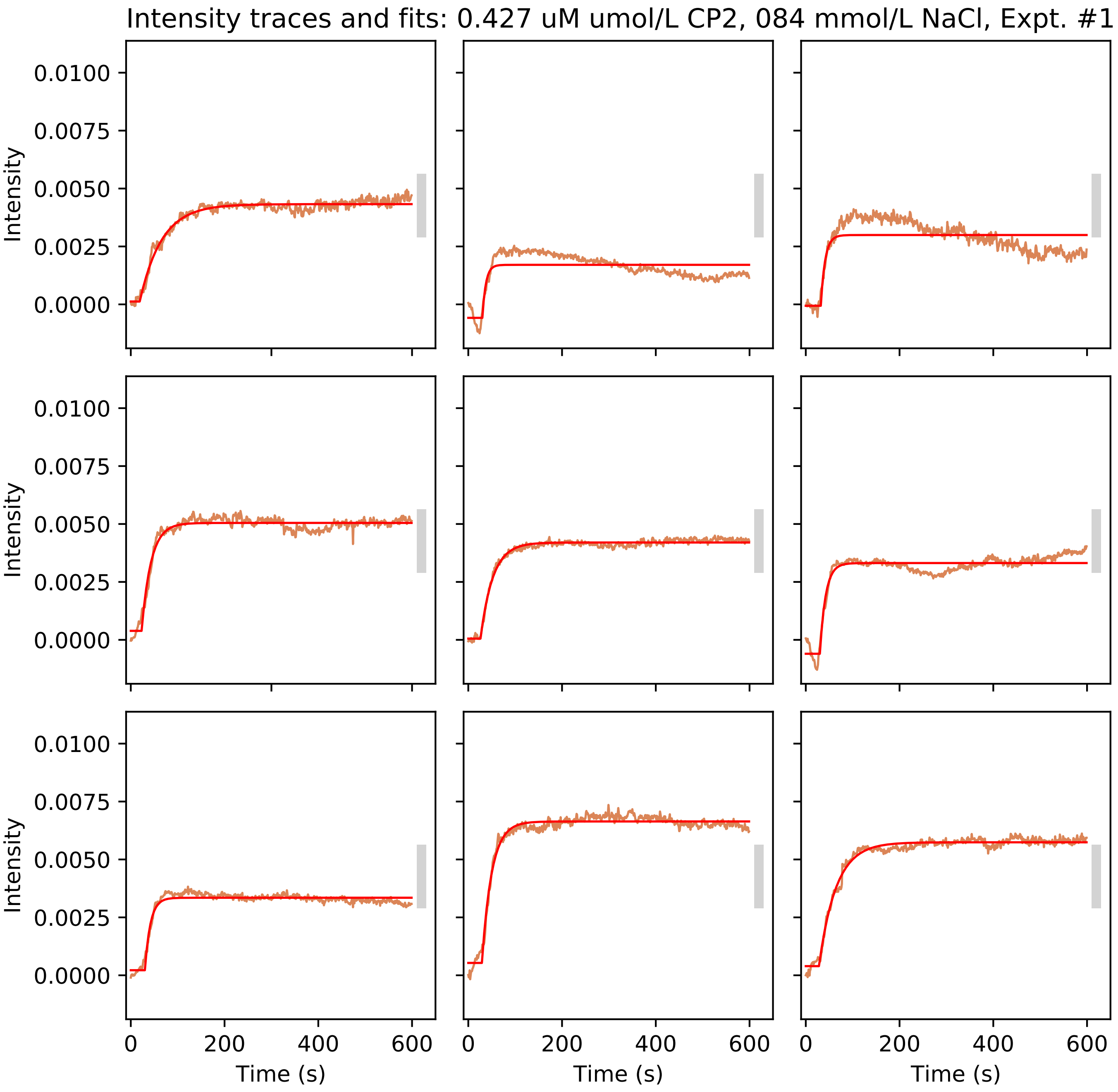
Intensity traces and fits: 0.427 uM umol/L CP2, 084 mmol/L NaCl, Expt. #1
Intensity
0.0100
0.0075
0.0050
0.0025
0.0000
0
200
400
600
Time (s)

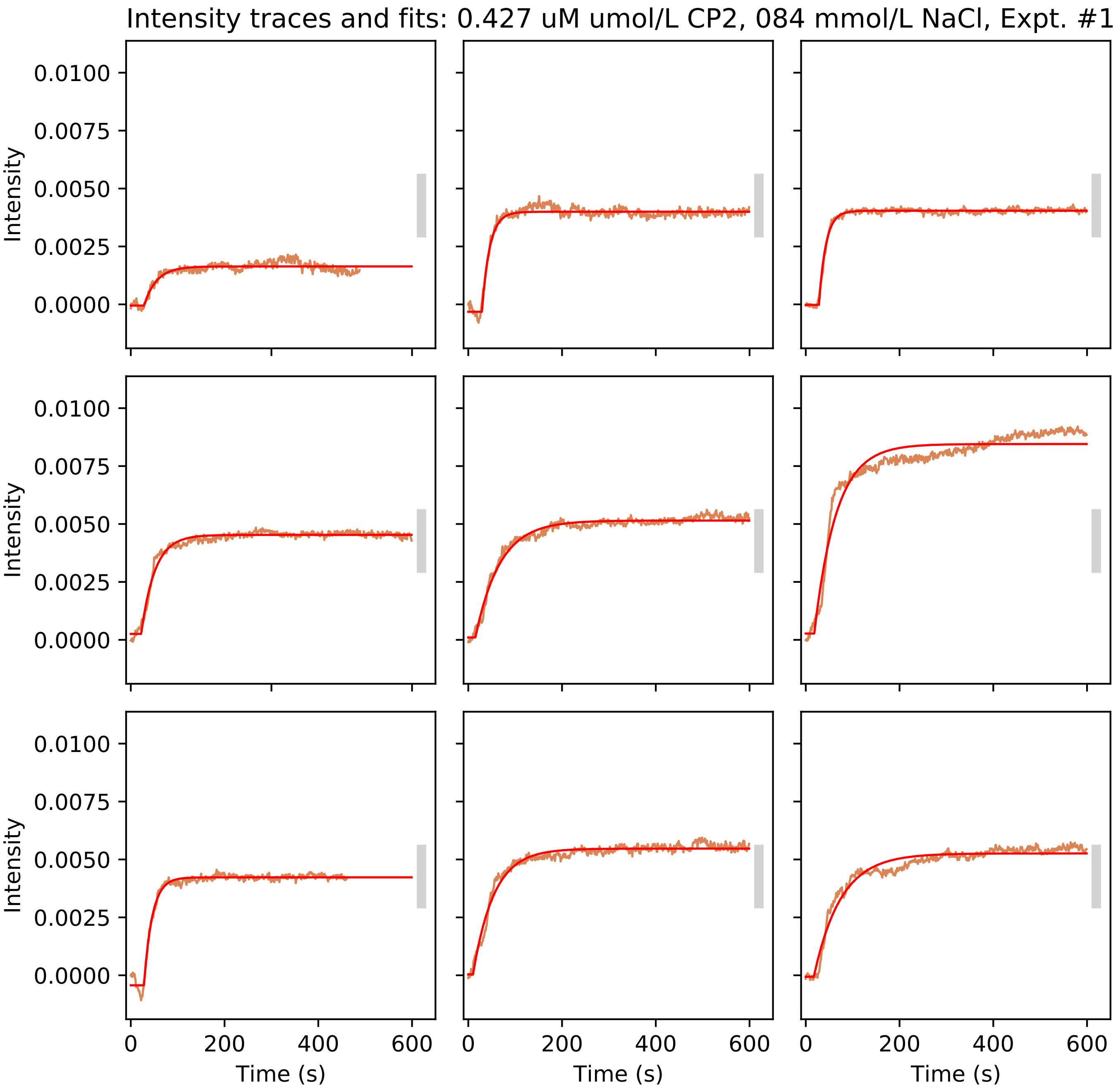
Intensity traces and fits: 0.427 uM umol/L CP2, 084 mmol/L NaCl, Expt. #1
Intensity
0.0100
0.0075
0.0050
0.0025
0.0000
Time (s)
0
200
400
600

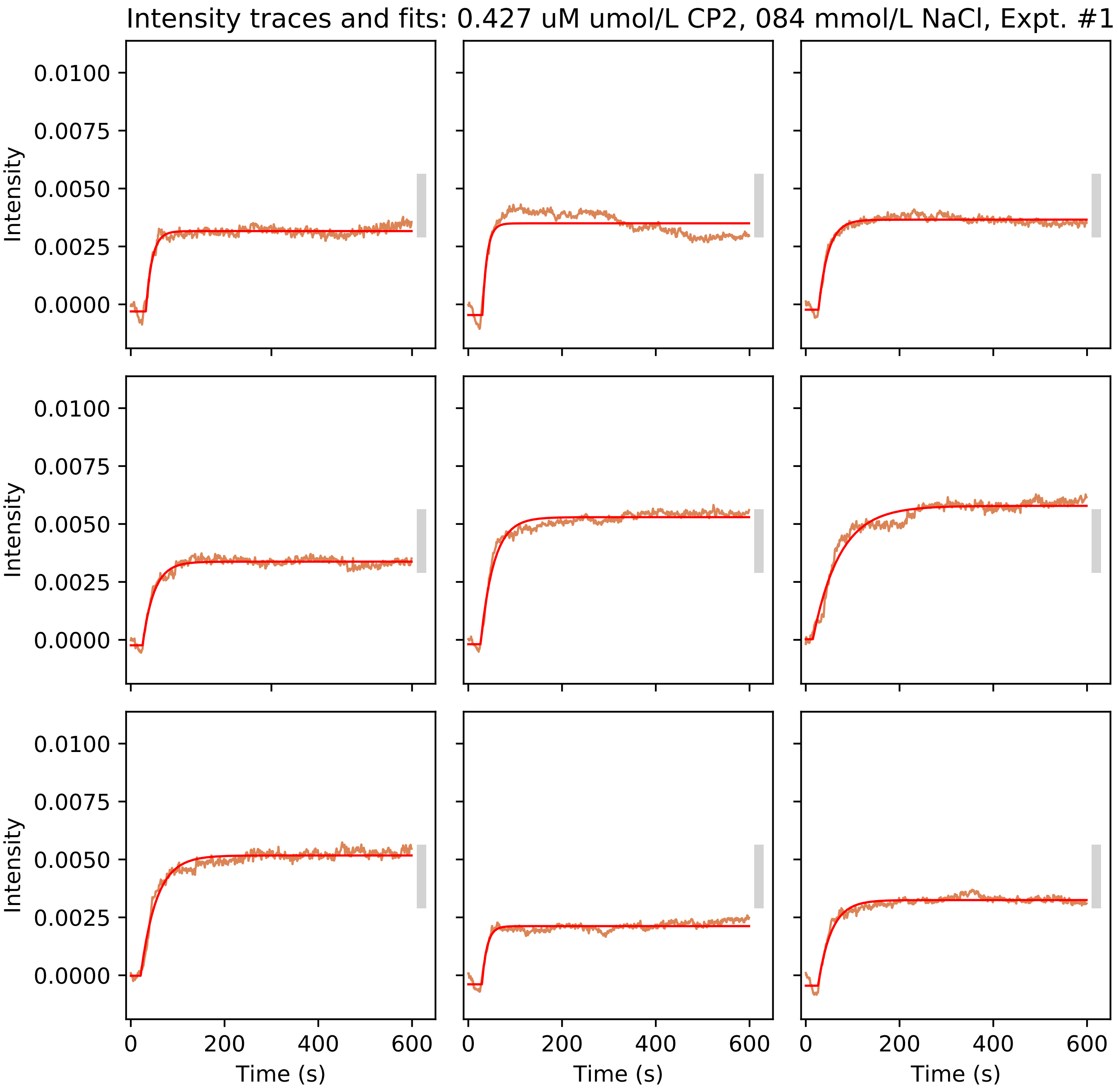
Intensity traces and fits: 0.427 uM umol/L CP2, 084 mmol/L NaCl, Expt. #1
Intensity
0.0100
0.0075
0.0050
0.0025
0.0000
Time (s)
0
200
400
600

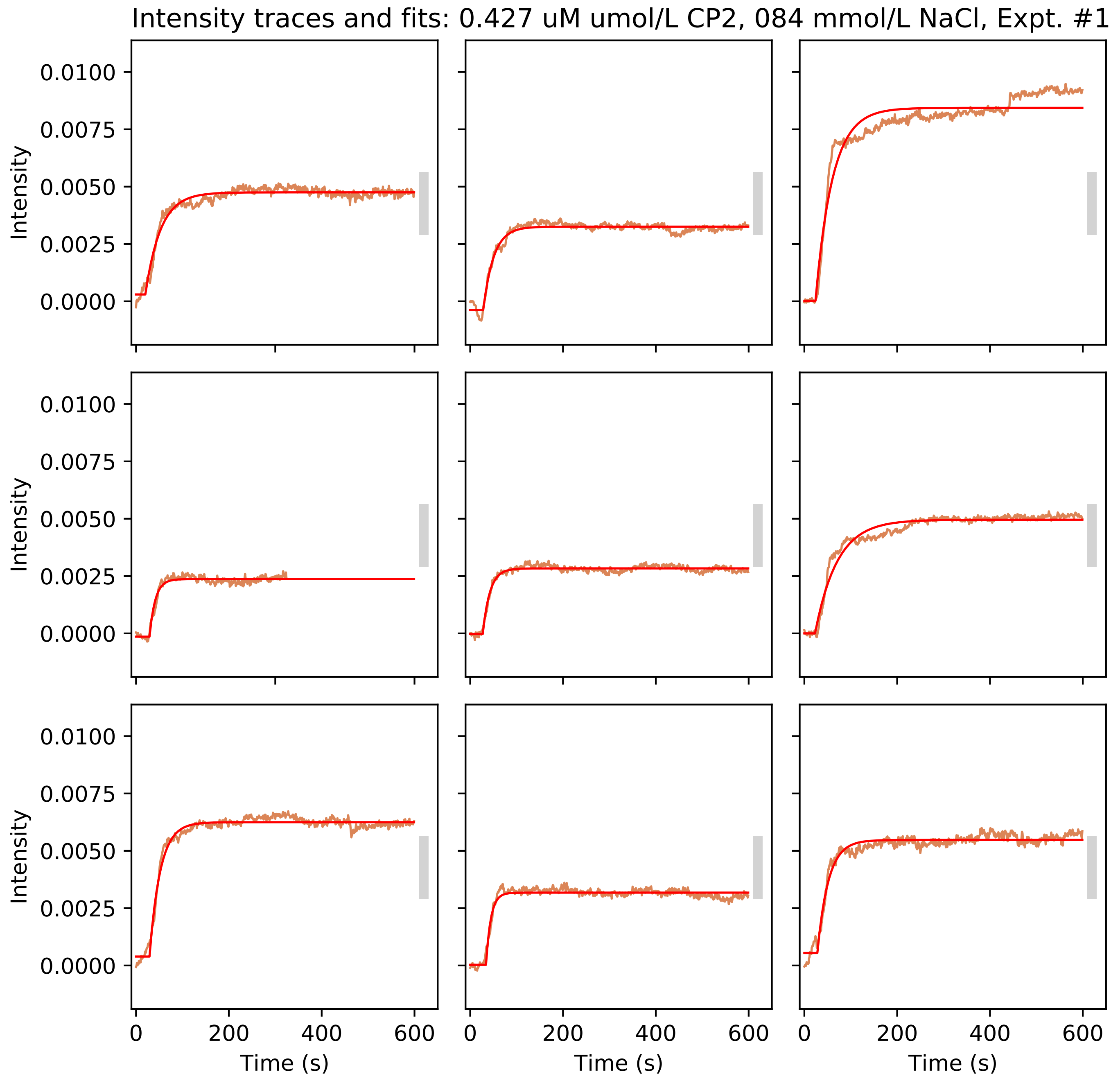

Intensity traces and fits: 0.427 uM umol/L CP2, 084 mmol/L NaCl, Expt. #1
Intensity
0.0100
0.0075
0.0050
0.0025
0.0000
0
200
400
600
Time (s)

Intensity traces and fits: 0.427 uM umol/L CP2, 084 mmol/L NaCl, Expt. #1

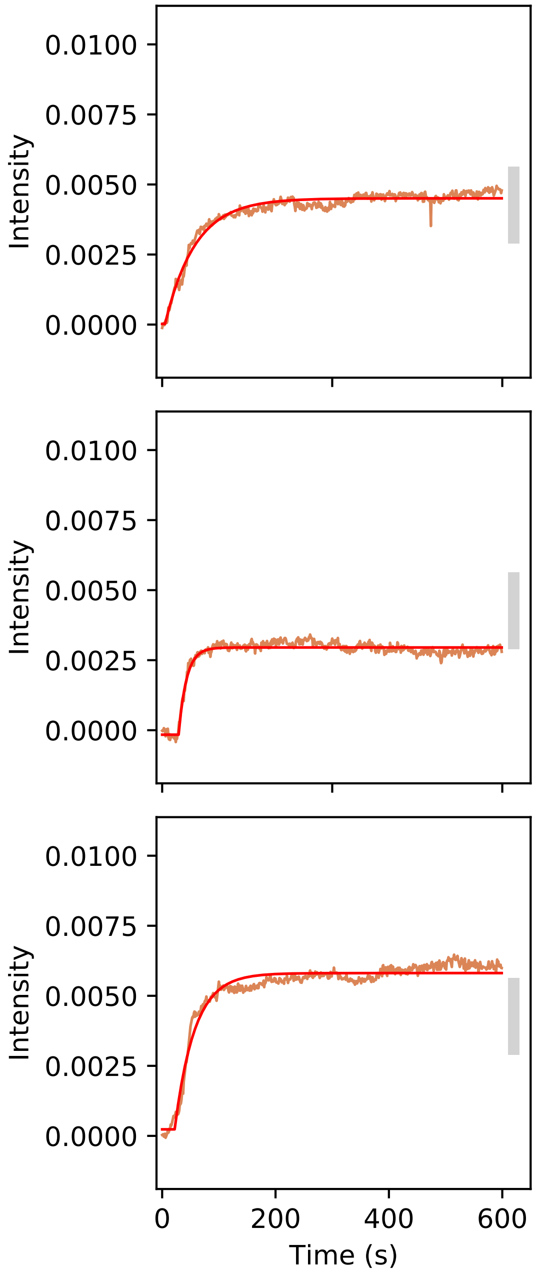

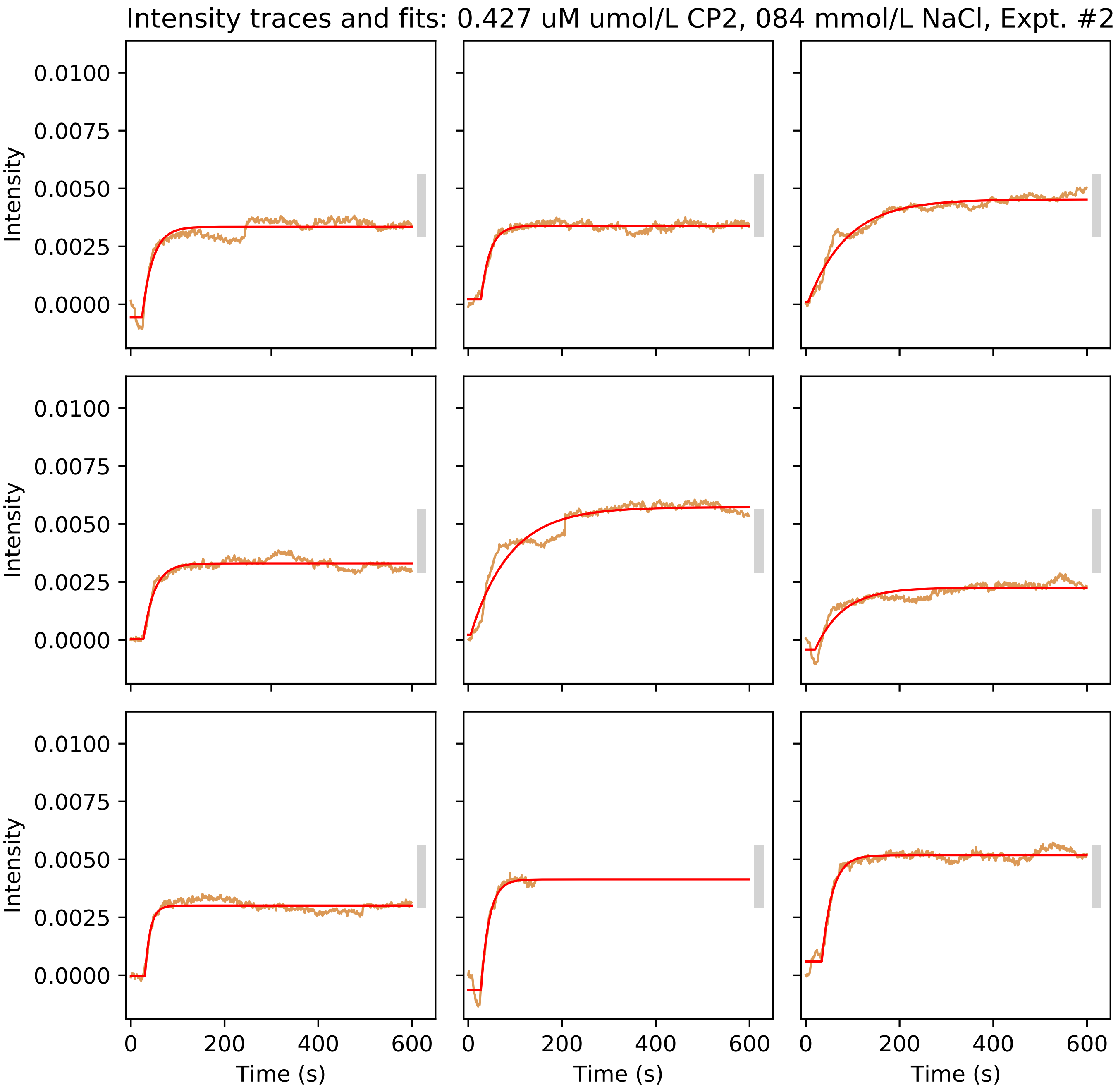

Intensity traces and fits: 0.427 uM umol/L CP2, 084 mmol/L NaCl, Expt. #2
Intensity
0.0100
0.0075
0.0050
0.0025
0.0000
Time (s)
0
200
400
600

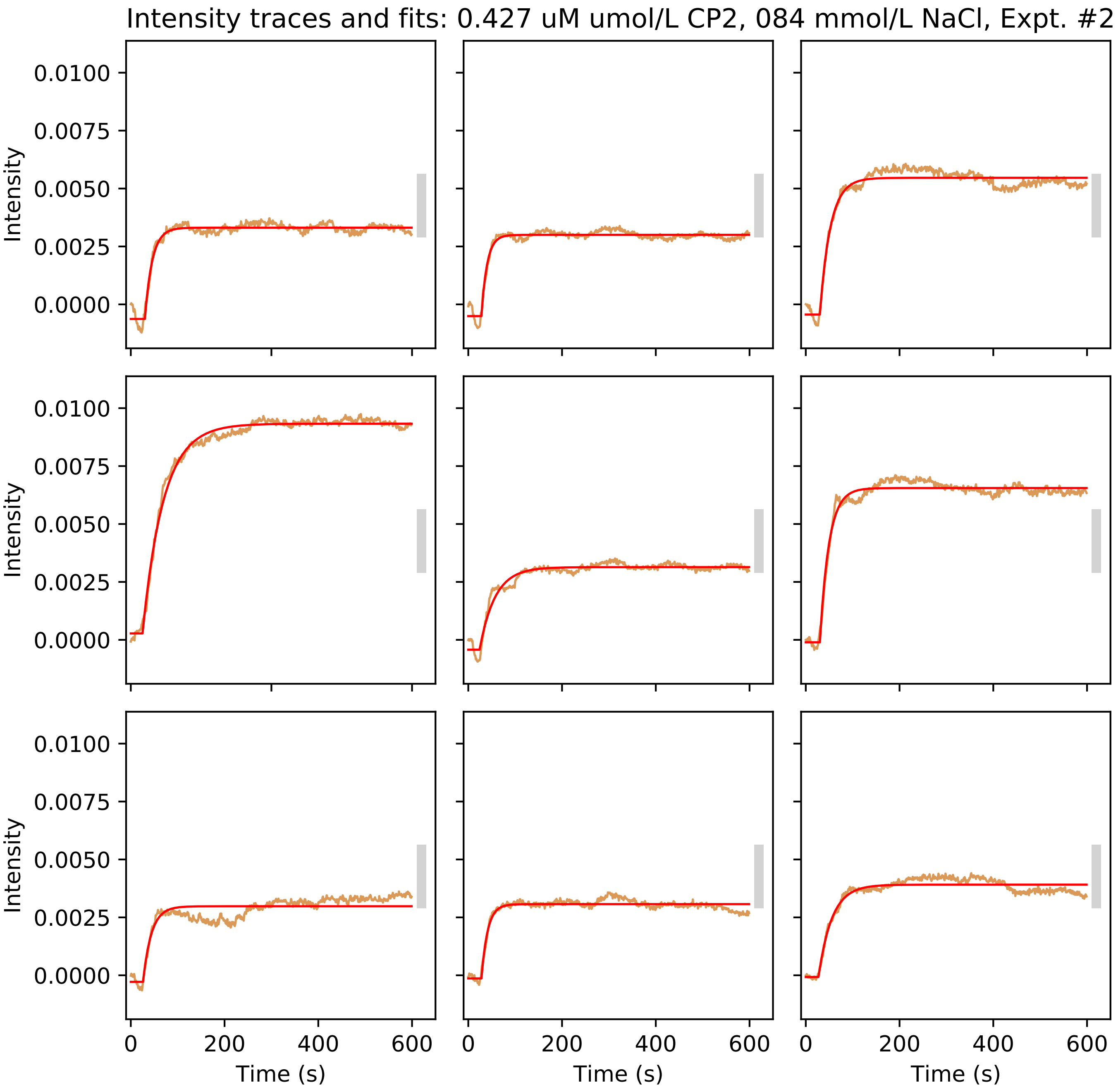
Intensity traces and fits: 0.427 uM umol/L CP2, 084 mmol/L NaCl, Expt. #2
Intensity
0.0100
0.0075
0.0050
0.0025
0.0000
0
200
400
600
Time (s)

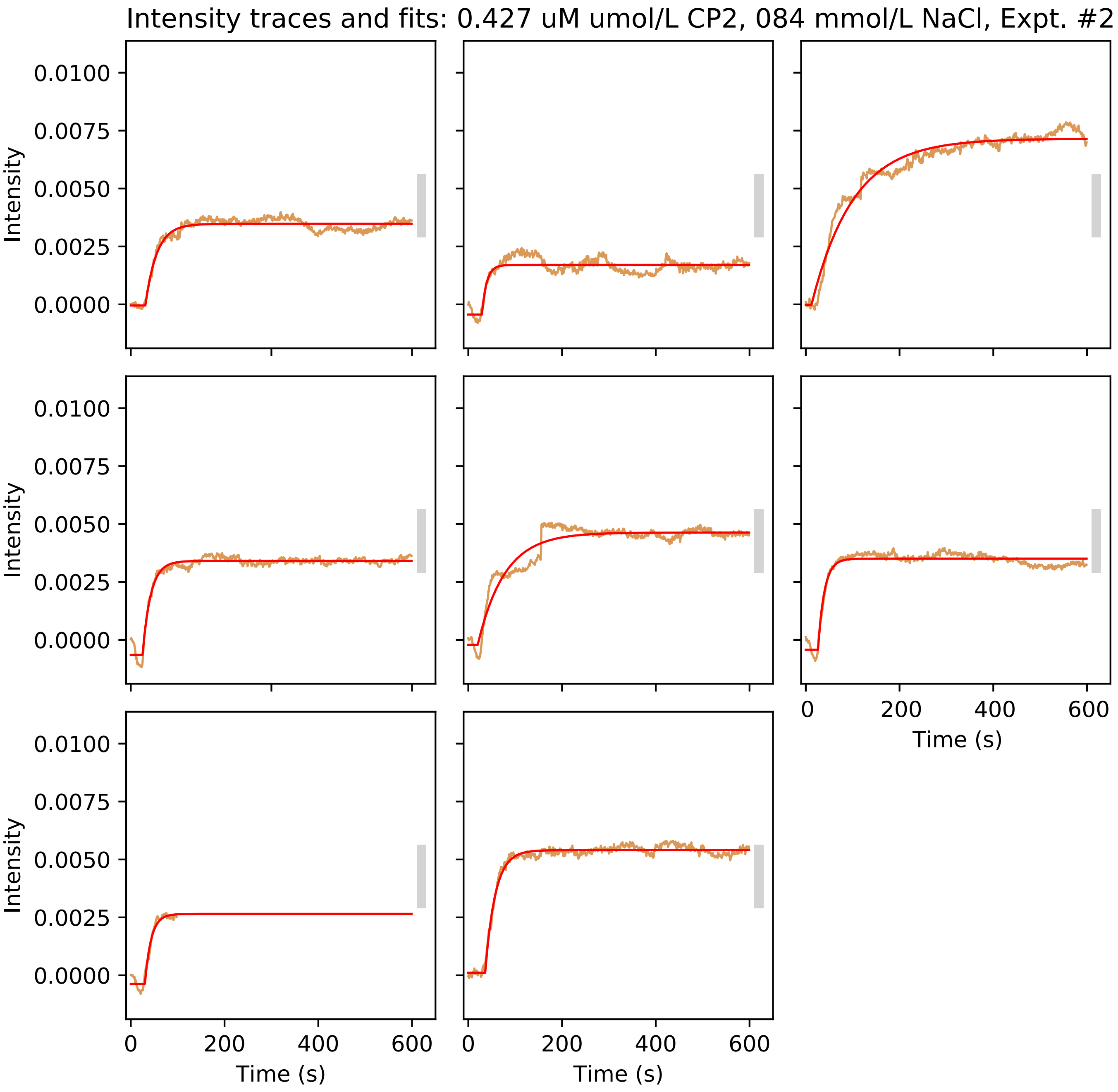
Intensity traces and fits: 0.427 uM umol/L CP2, 084 mmol/L NaCl, Expt. #2
Intensity
0.0100
0.0075
0.0050
0.0025
0.0000
0
200
400
600
Time (s)

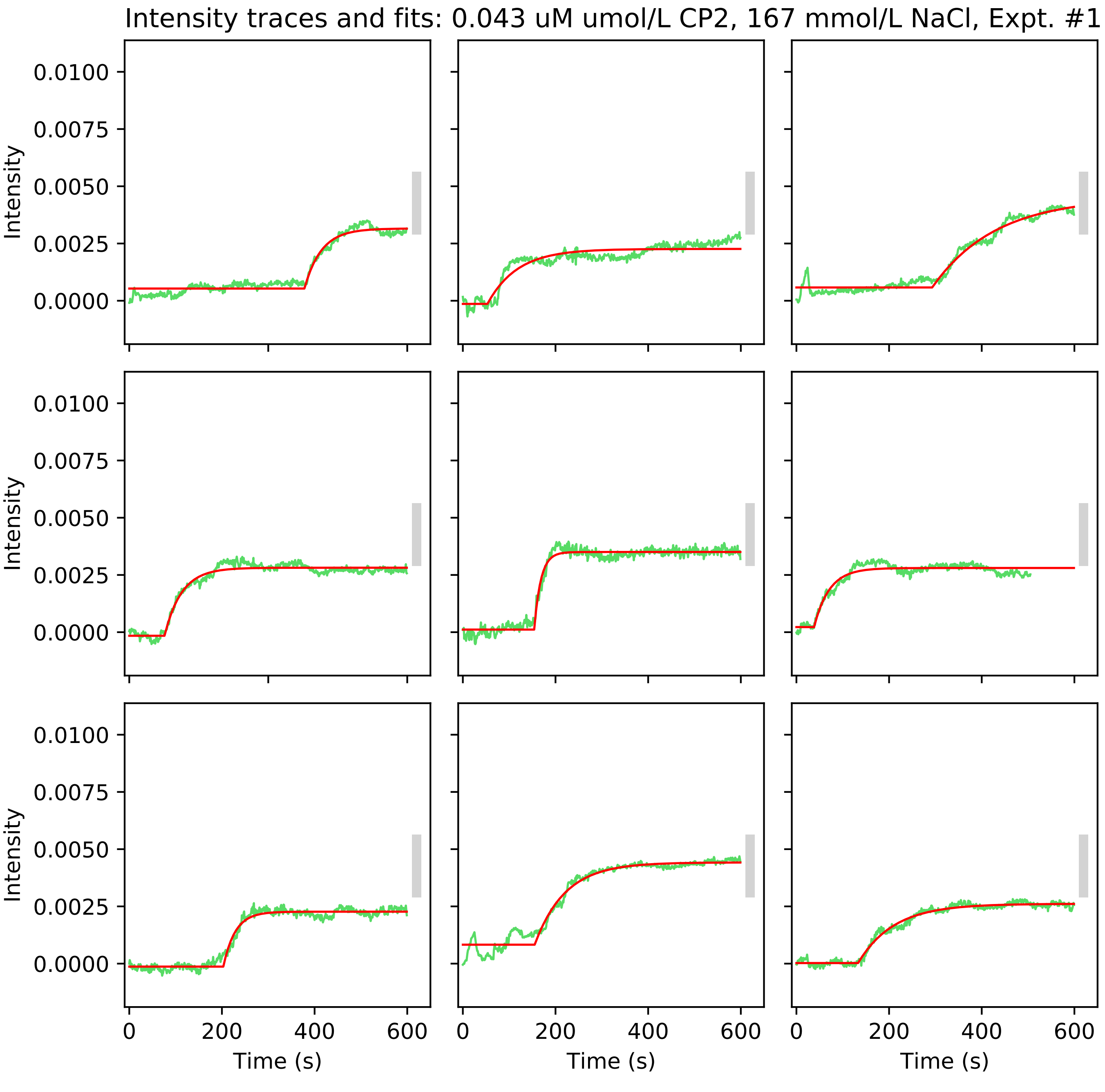

Intensity traces and fits: 0.043 uM umol/L CP2, 167 mmol/L NaCl, Expt. #1
Intensity
0.0100
0.0075
0.0050
0.0025
0.0000
Time (s)
0
200
400
600

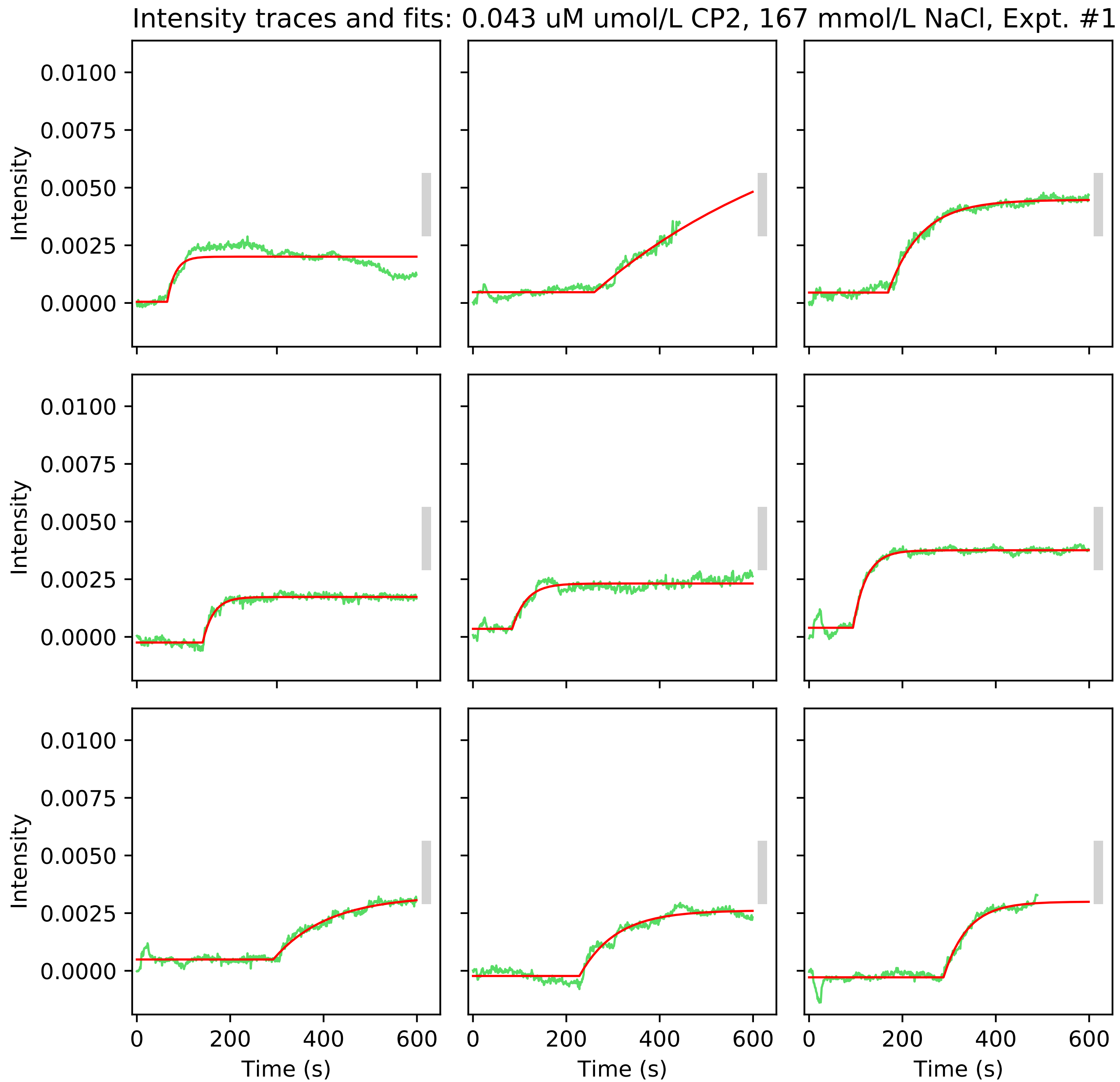

Intensity traces and fits: 0.043 uM umol/L CP2, 167 mmol/L NaCl, Expt. #1
Intensity
0.0100
0.0075
0.0050
0.0025
0.0000
0
200
400
600
Time (s)

Intensity traces and fits: 0.043 uM umol/L CP2, 167 mmol/L NaCl, Expt. #1

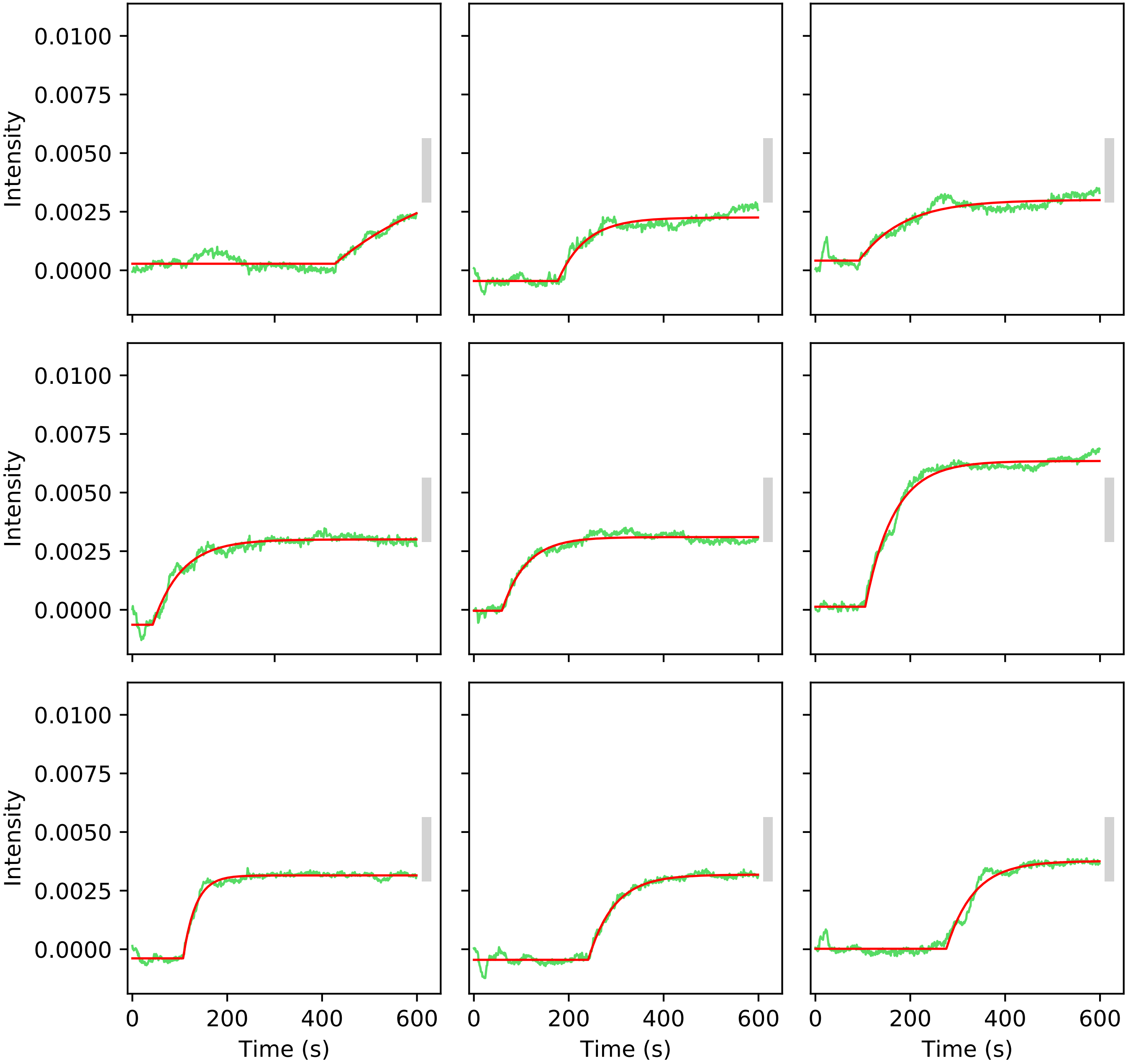

Intensity traces and fits: 0.043 uM umol/L CP2, 167 mmol/L NaCl, Expt. #1

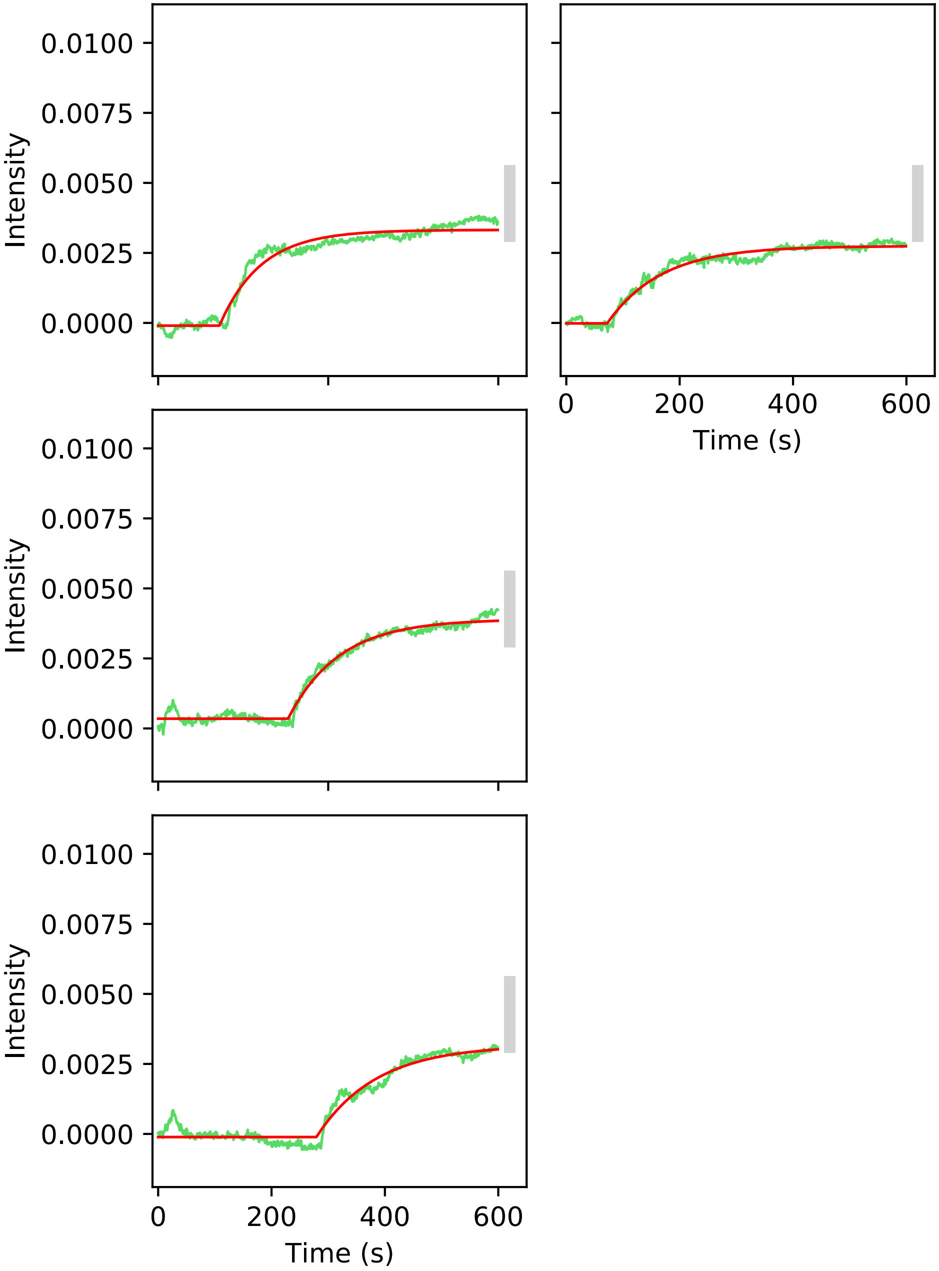

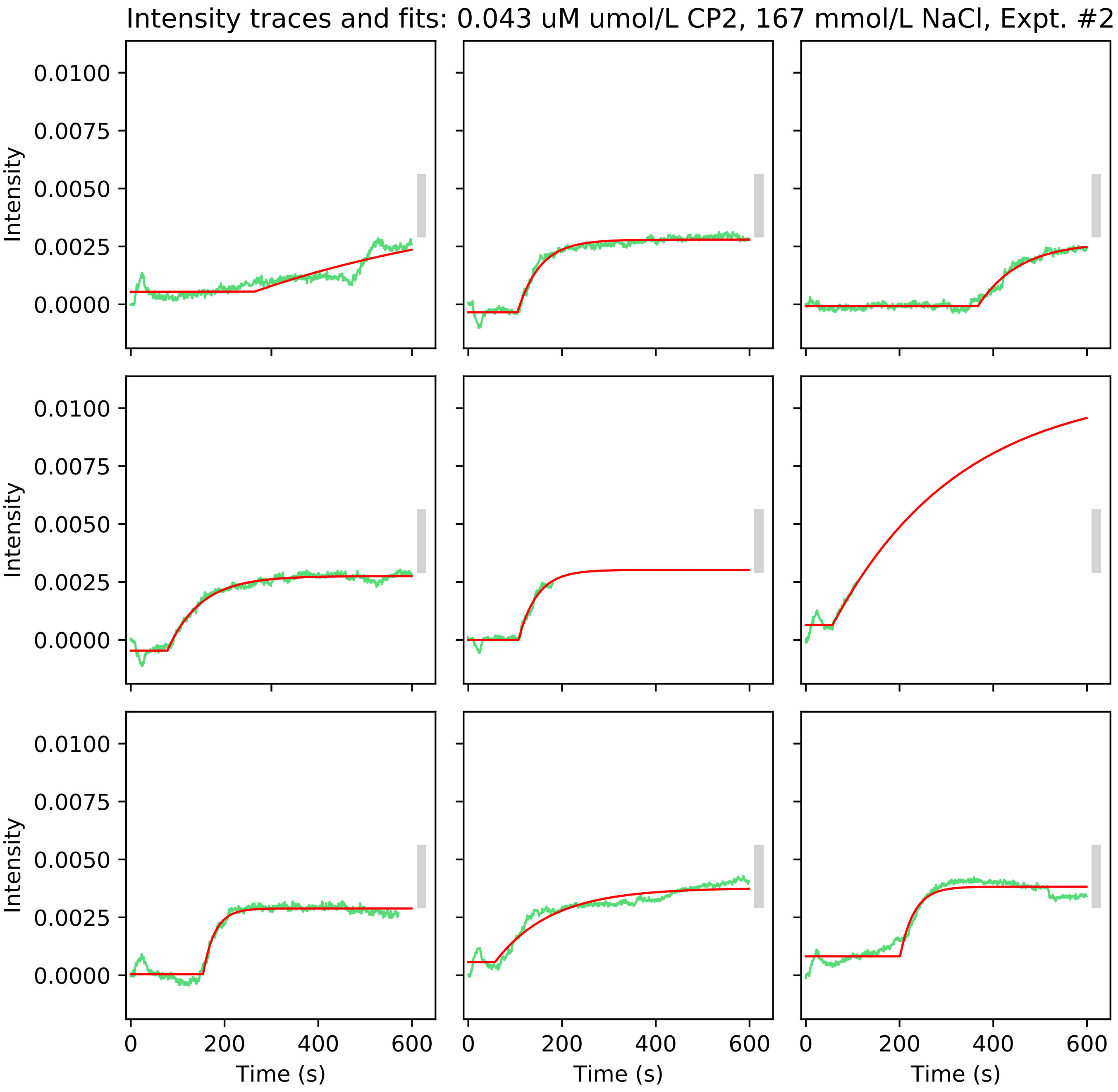
Intensity traces and fits: 0.043 uM umol/L CP2, 167 mmol/L NaCl, Expt. #2
Intensity
0.0100
0.0075
0.0050
0.0025
0.0000
0
200
400
600
Time (s)

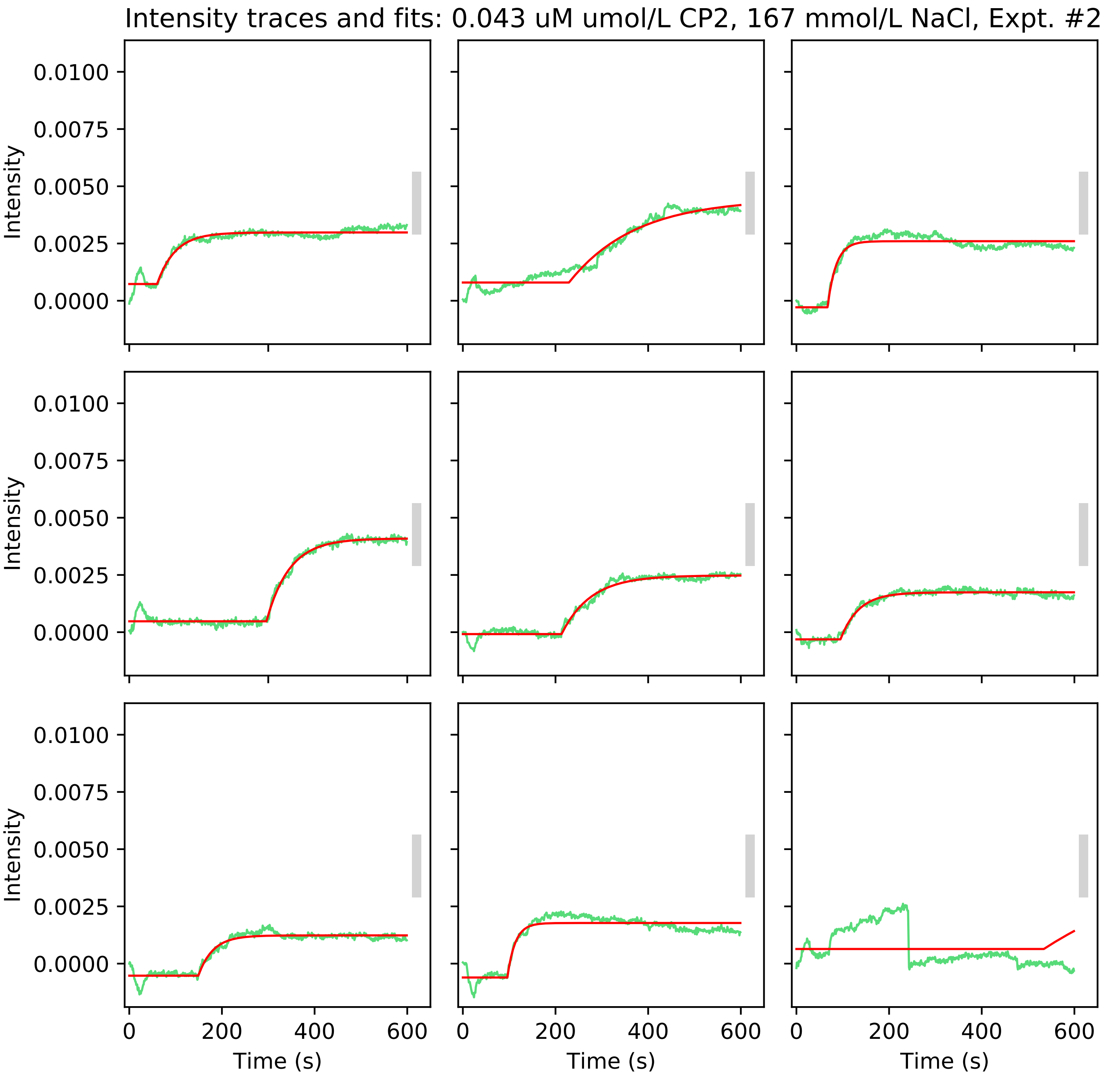

Intensity traces and fits: 0.043 uM umol/L CP2, 167 mmol/L NaCl, Expt. #2
Intensity
0.0100
0.0075
0.0050
0.0025
0.0000
0
200
400
600
Time (s)

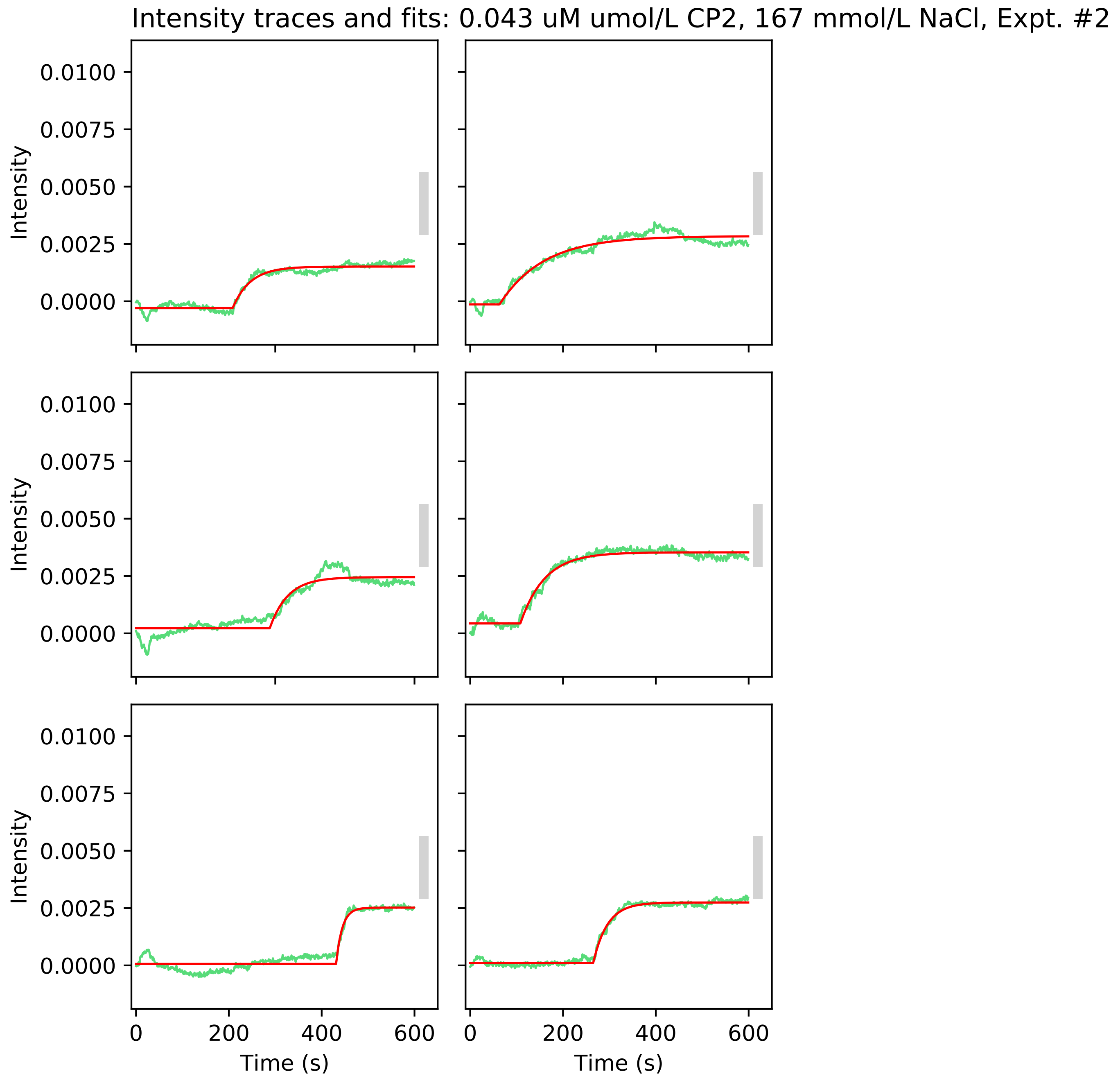
Intensity traces and fits: 0.043 uM umol/L CP2, 167 mmol/L NaCl, Expt. #2
Intensity
0.0100
0.0075
0.0050
0.0025
0.0000
Time (s)
0
200
400
600

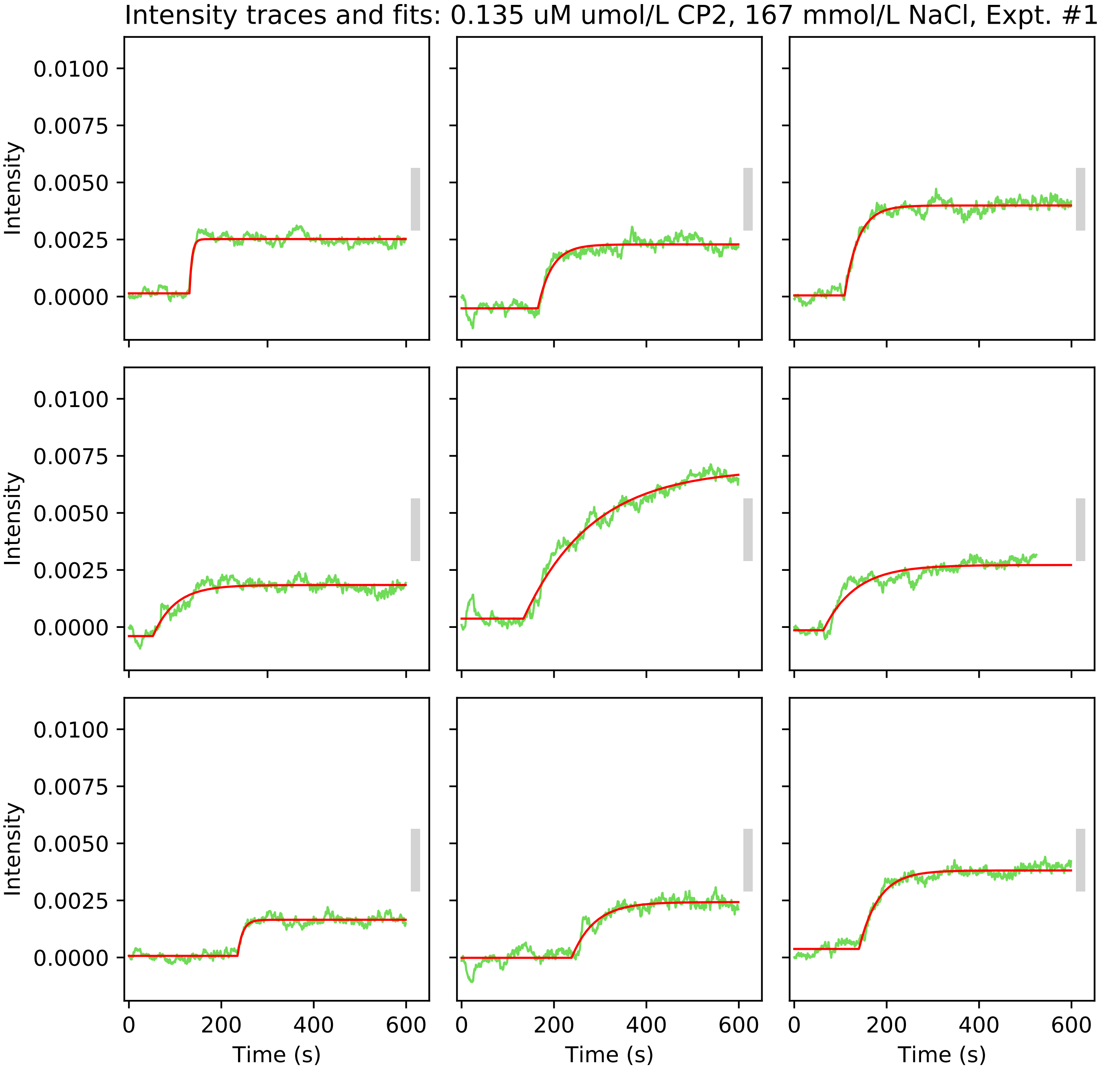
Intensity traces and fits: 0.135 uM umol/L CP2, 167 mmol/L NaCl, Expt. #1
Intensity
0.0100
0.0075
0.0050
0.0025
0.0000
0
200
400
600
Time (s)

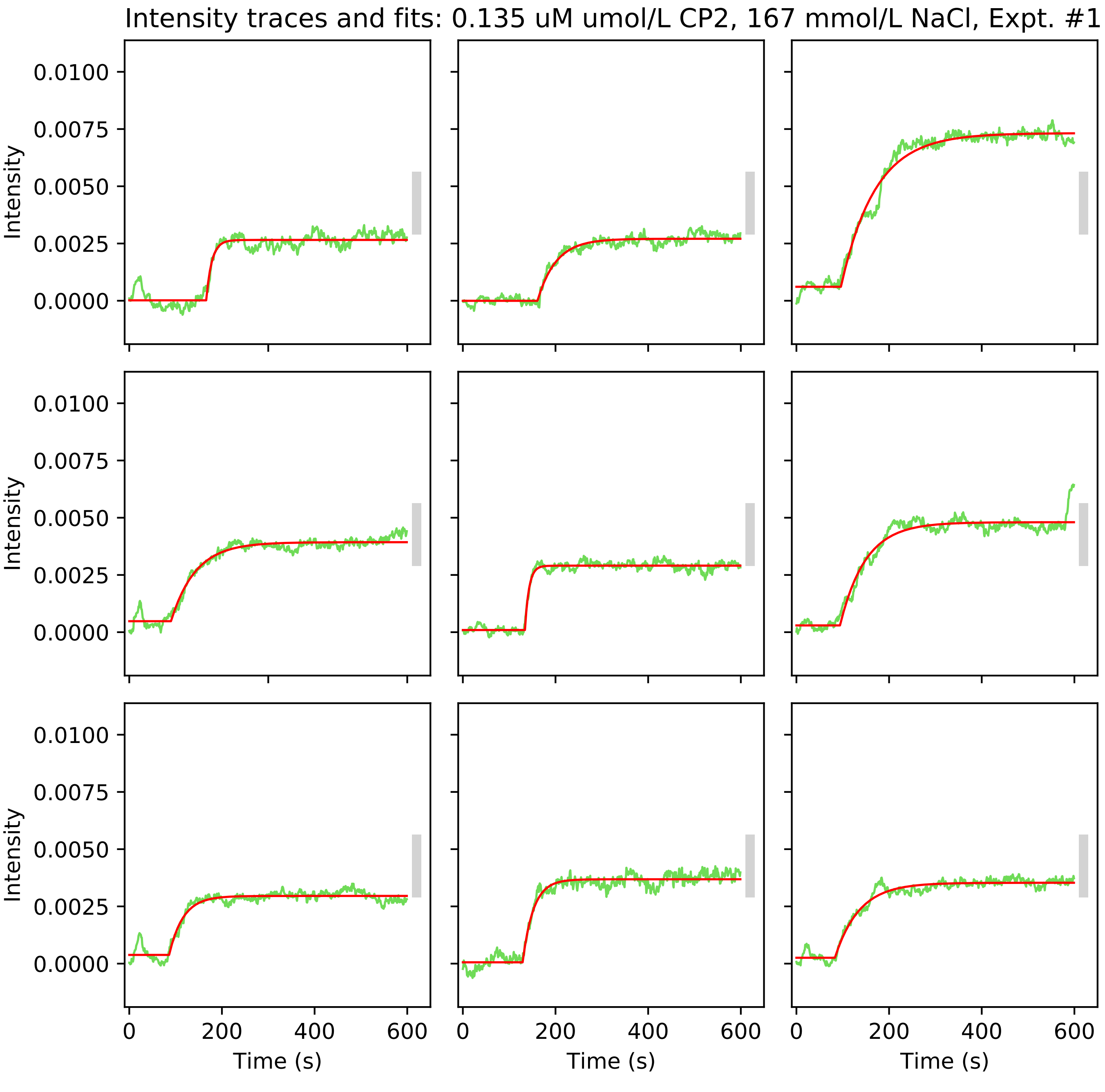
Intensity traces and fits: 0.135 uM umol/L CP2, 167 mmol/L NaCl, Expt. #1
Intensity
0.0100
0.0075
0.0050
0.0025
0.0000
Time (s)
0
200
400
600

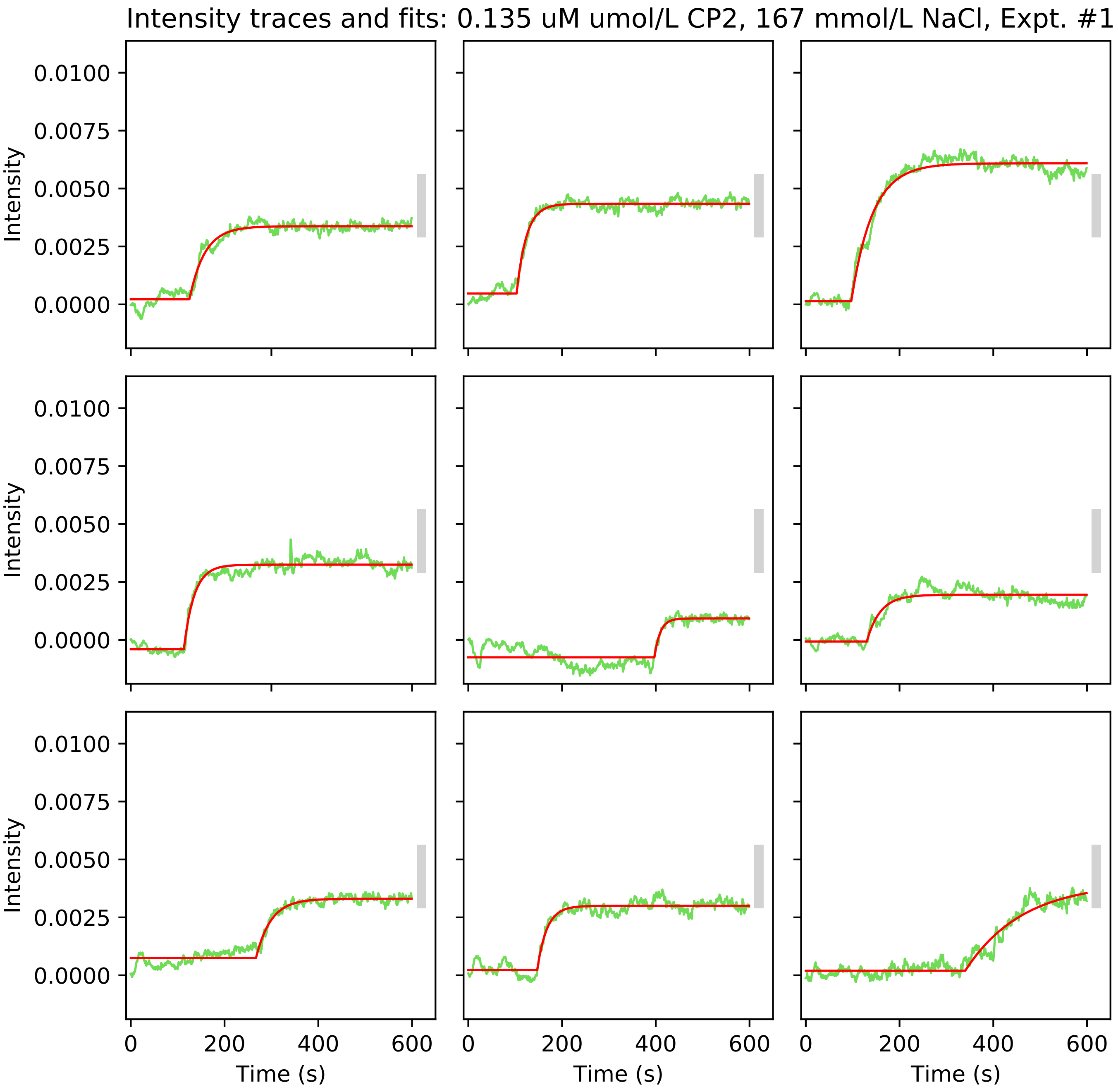

Intensity traces and fits: 0.135 uM umol/L CP2, 167 mmol/L NaCl, Expt. #1
Intensity
0.0100
0.0075
0.0050
0.0025
0.0000
0
200
400
600
Time (s)

Intensity traces and fits: 0.135 uM umol/L CP2, 167 mmol/L NaCl, Expt. #1

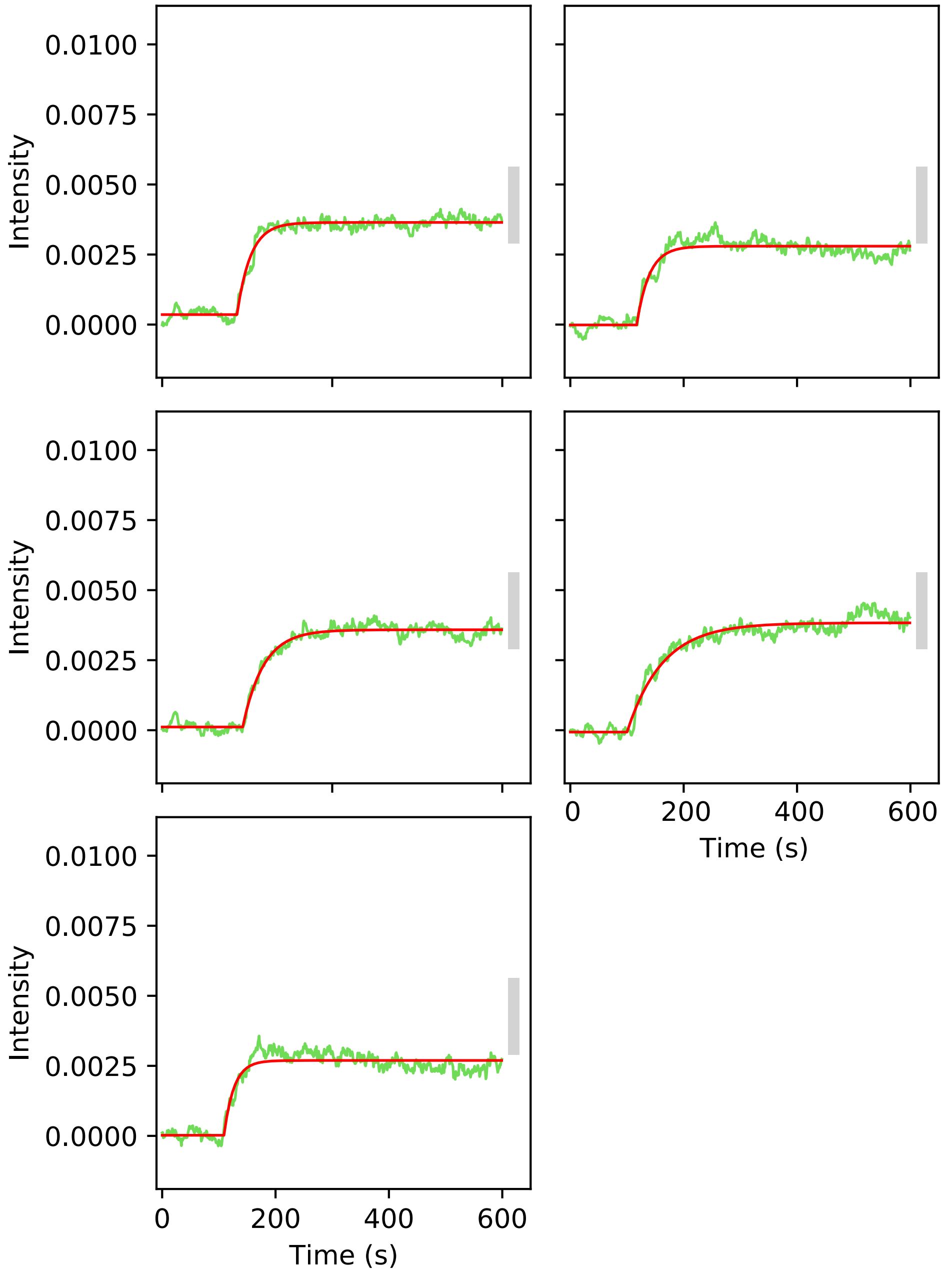

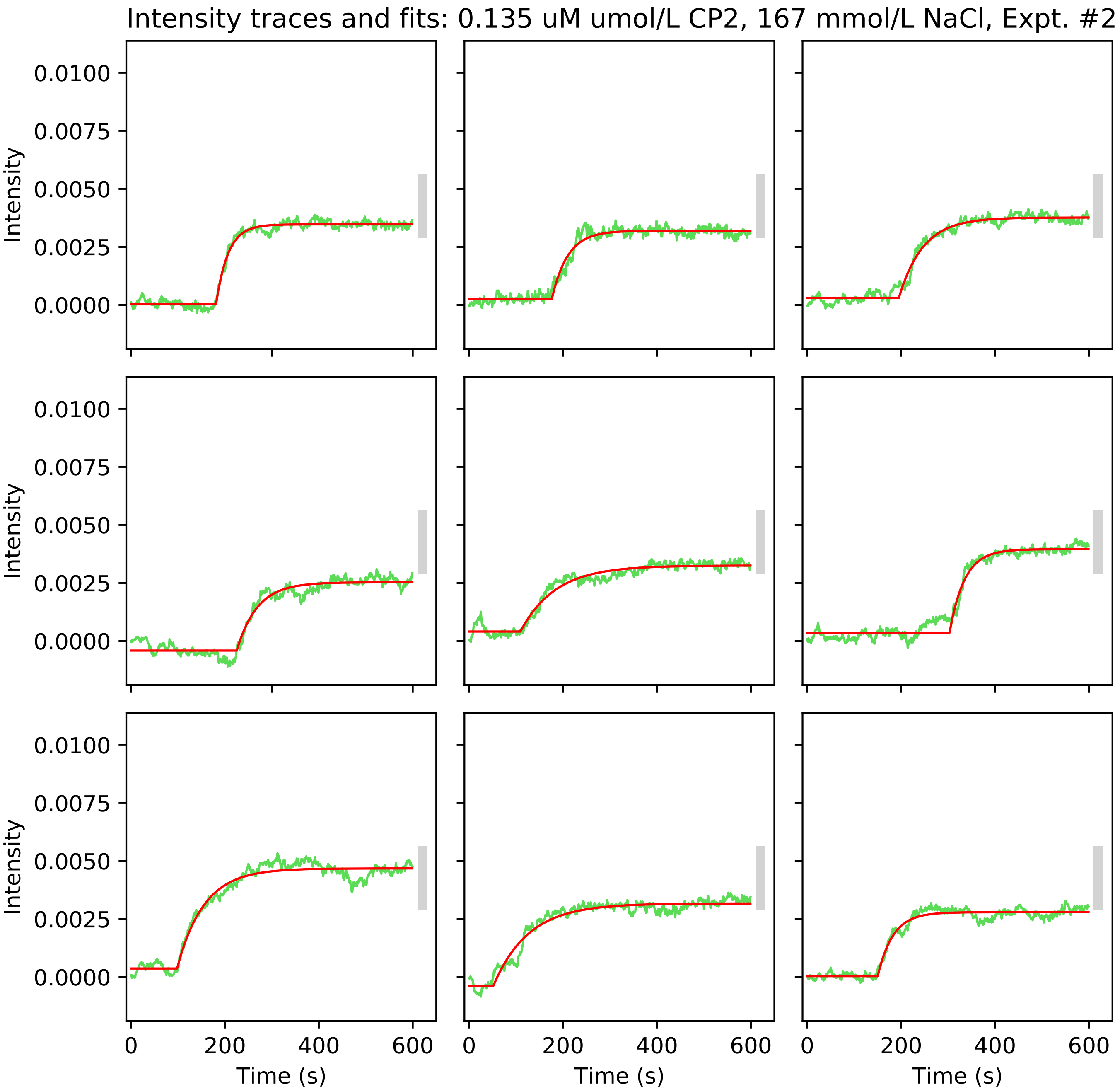
Intensity traces and fits: 0.135 uM umol/L CP2, 167 mmol/L NaCl, Expt. #2
Intensity
0.0100
0.0075
0.0050
0.0025
0.0000
Time (s)
0
200
400
600

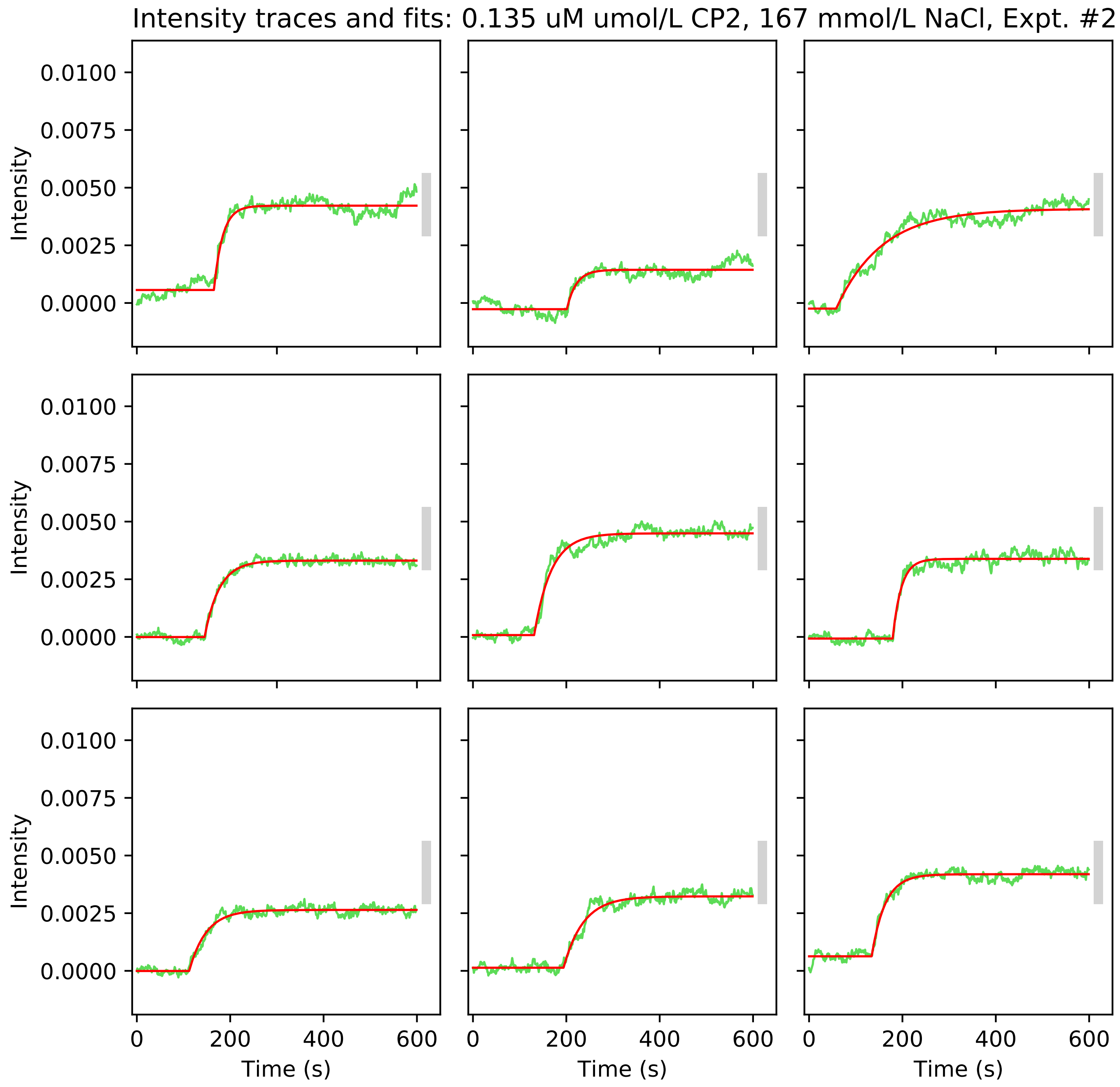
Intensity traces and fits: 0.135 uM umol/L CP2, 167 mmol/L NaCl, Expt. #2
Intensity
0.0100
0.0075
0.0050
0.0025
0.0000
0
200
400
600
Time (s)

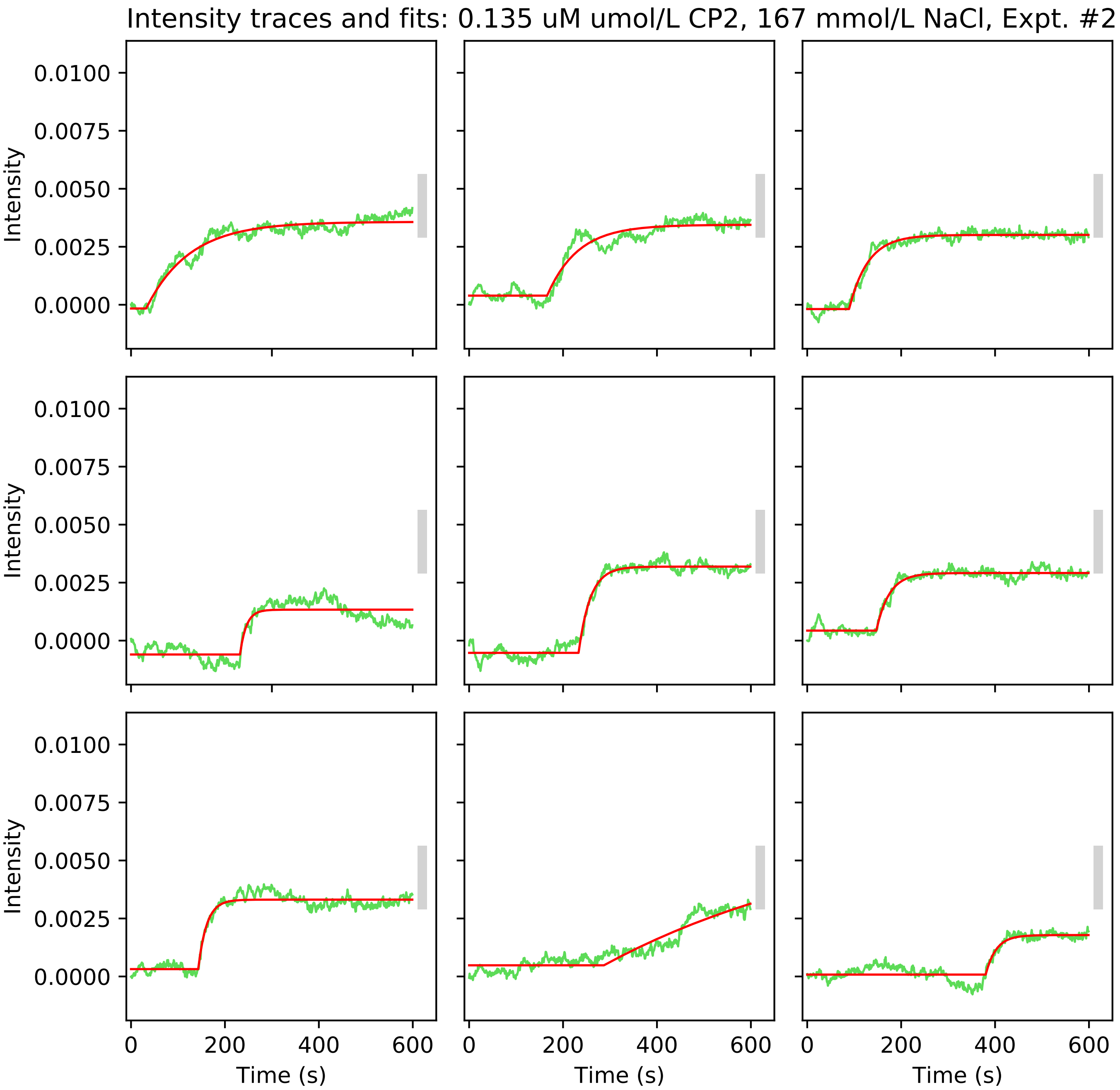
Intensity traces and fits: 0.135 uM umol/L CP2, 167 mmol/L NaCl, Expt. #2
Intensity
0.0100
0.0075
0.0050
0.0025
0.0000
Time (s)
0
200
400
600

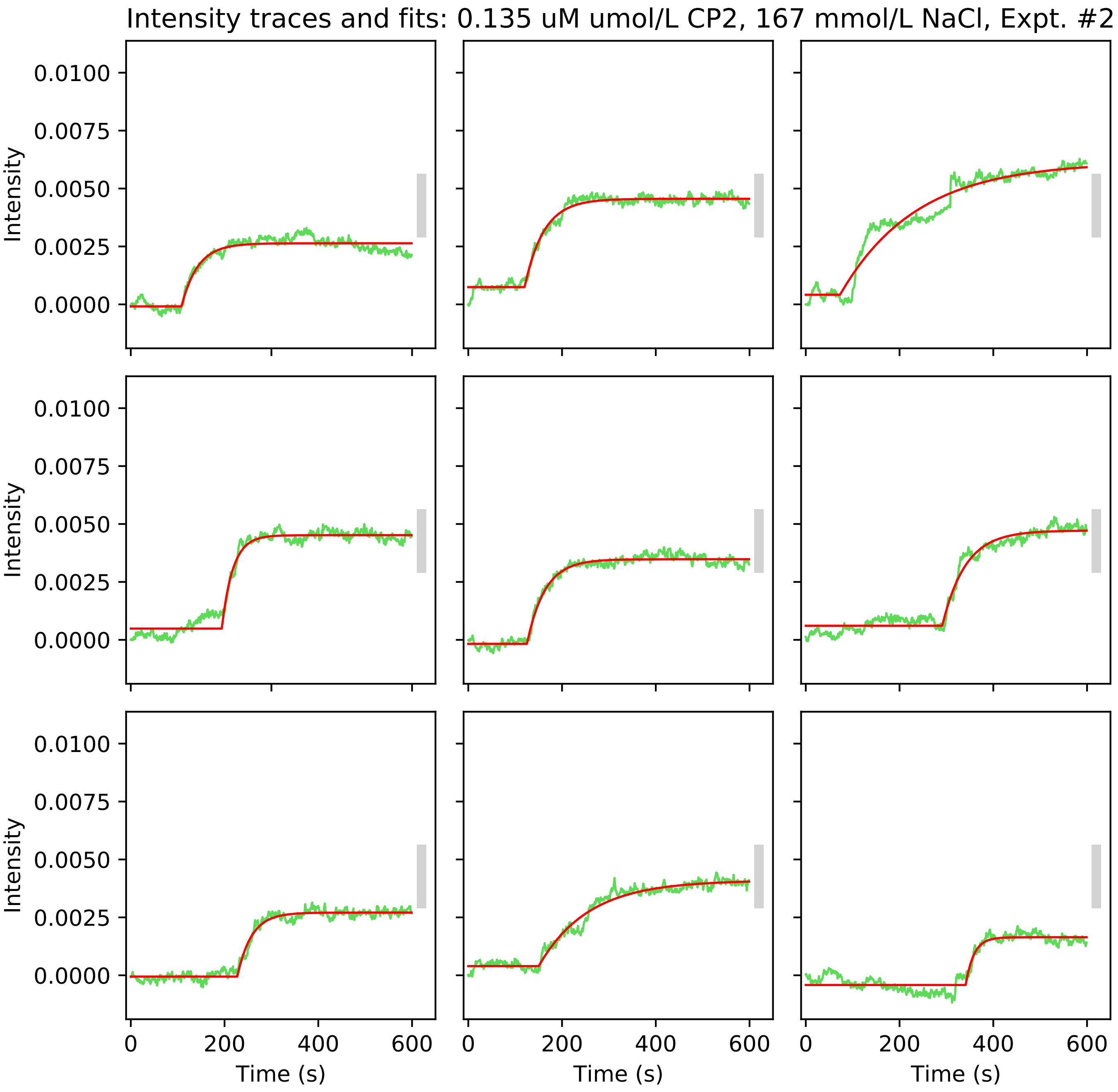

Intensity traces and fits: 0.135 uM umol/L CP2, 167 mmol/L NaCl, Expt. #2
Intensity
0.0100
0.0075
0.0050
0.0025
0.0000
0
200
400
600
Time (s)

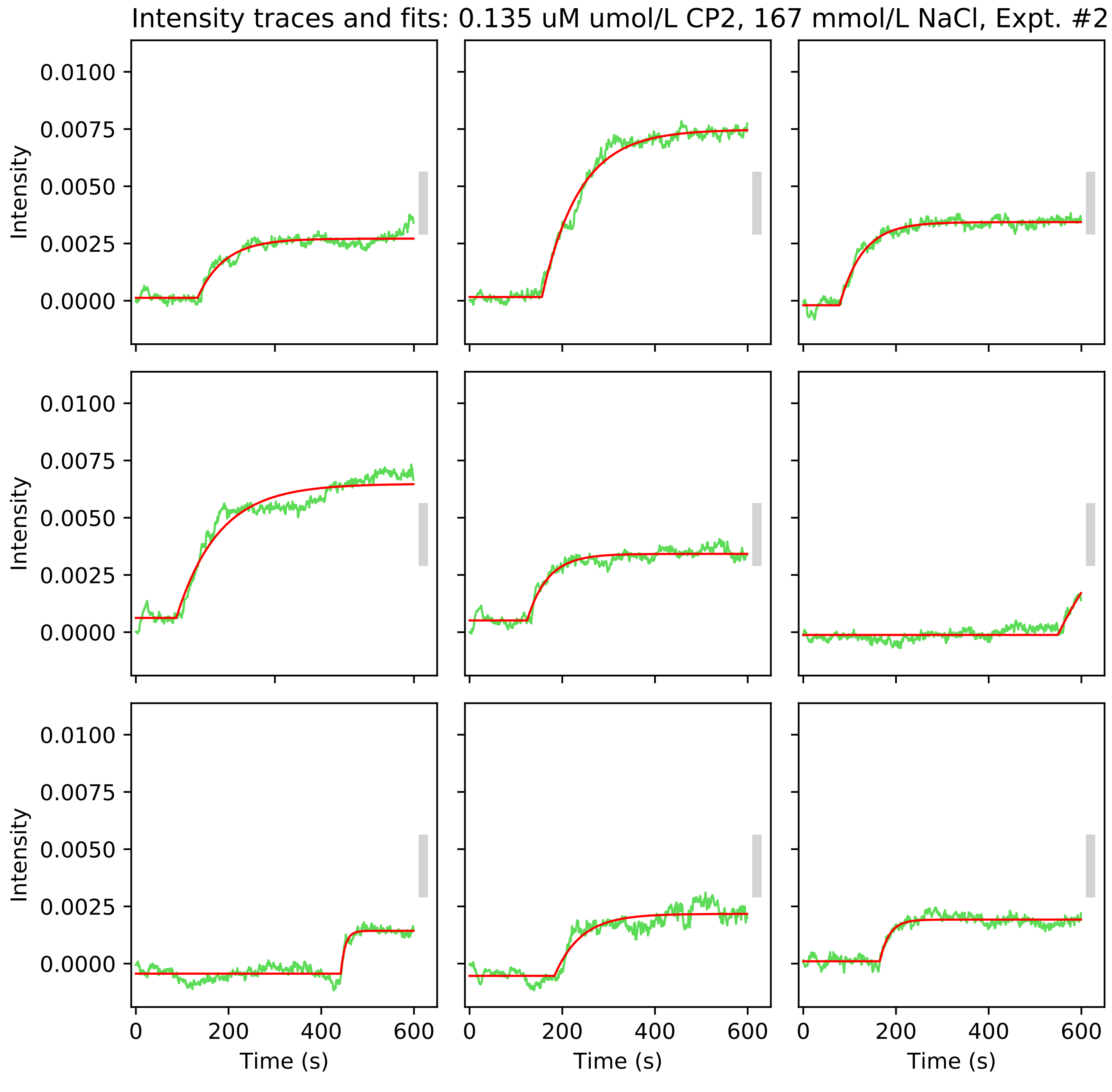

Intensity traces and fits: 0.135 uM umol/L CP2, 167 mmol/L NaCl, Expt. #2
Intensity
0.0100
0.0075
0.0050
0.0025
0.0000
0
200
400
600
Time (s)

Intensity traces and fits: 0.135 uM umol/L CP2, 167 mmol/L NaCl, Expt. #2

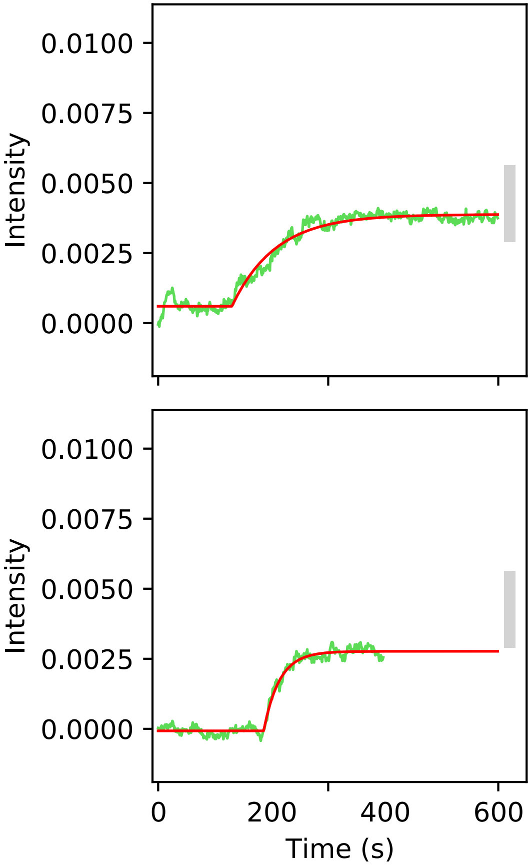

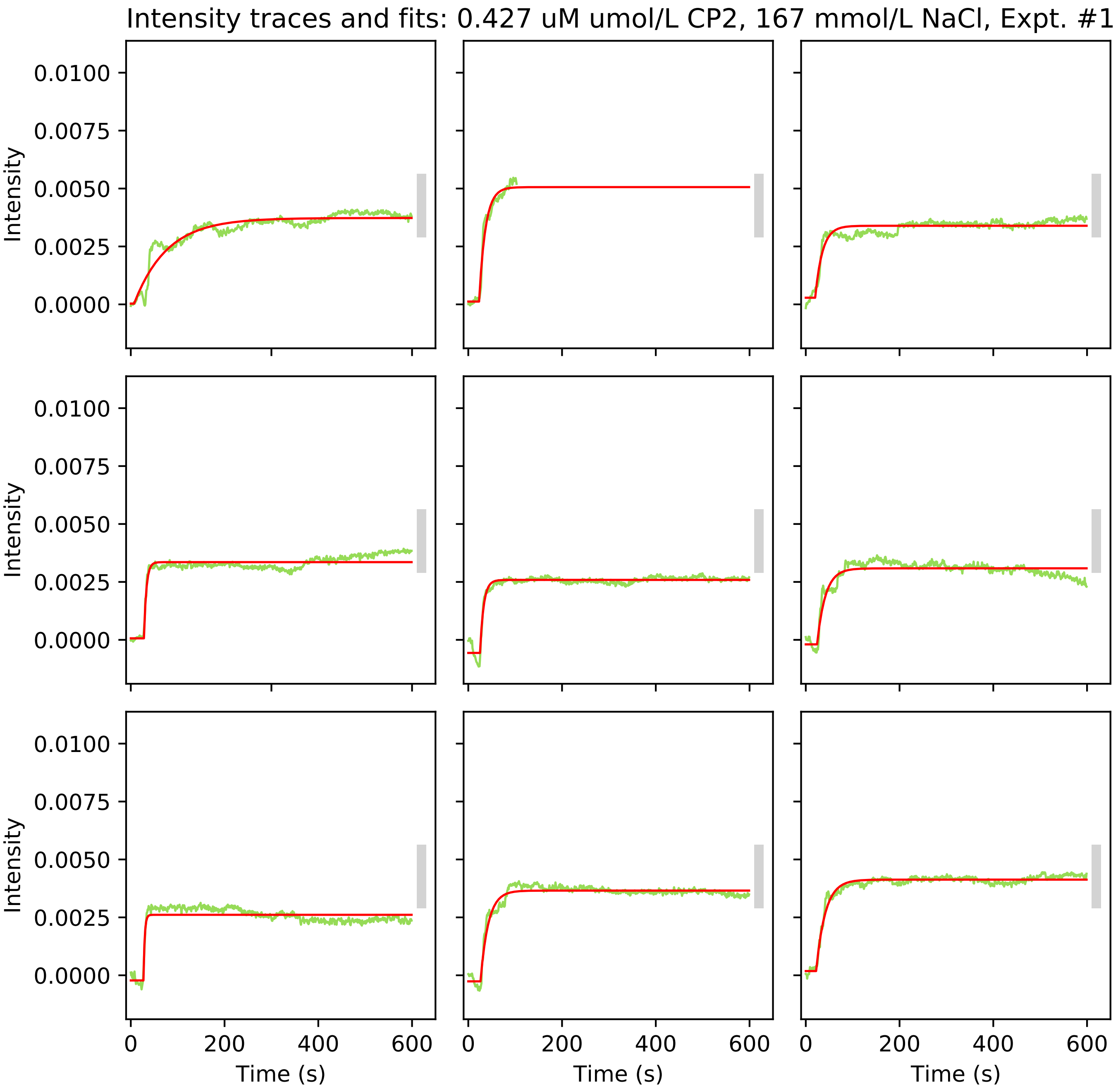
Intensity traces and fits: 0.427 uM umol/L CP2, 167 mmol/L NaCl, Expt. #1
Intensity
0.0100
0.0075
0.0050
0.0025
0.0000
Time (s)
0
200
400
600

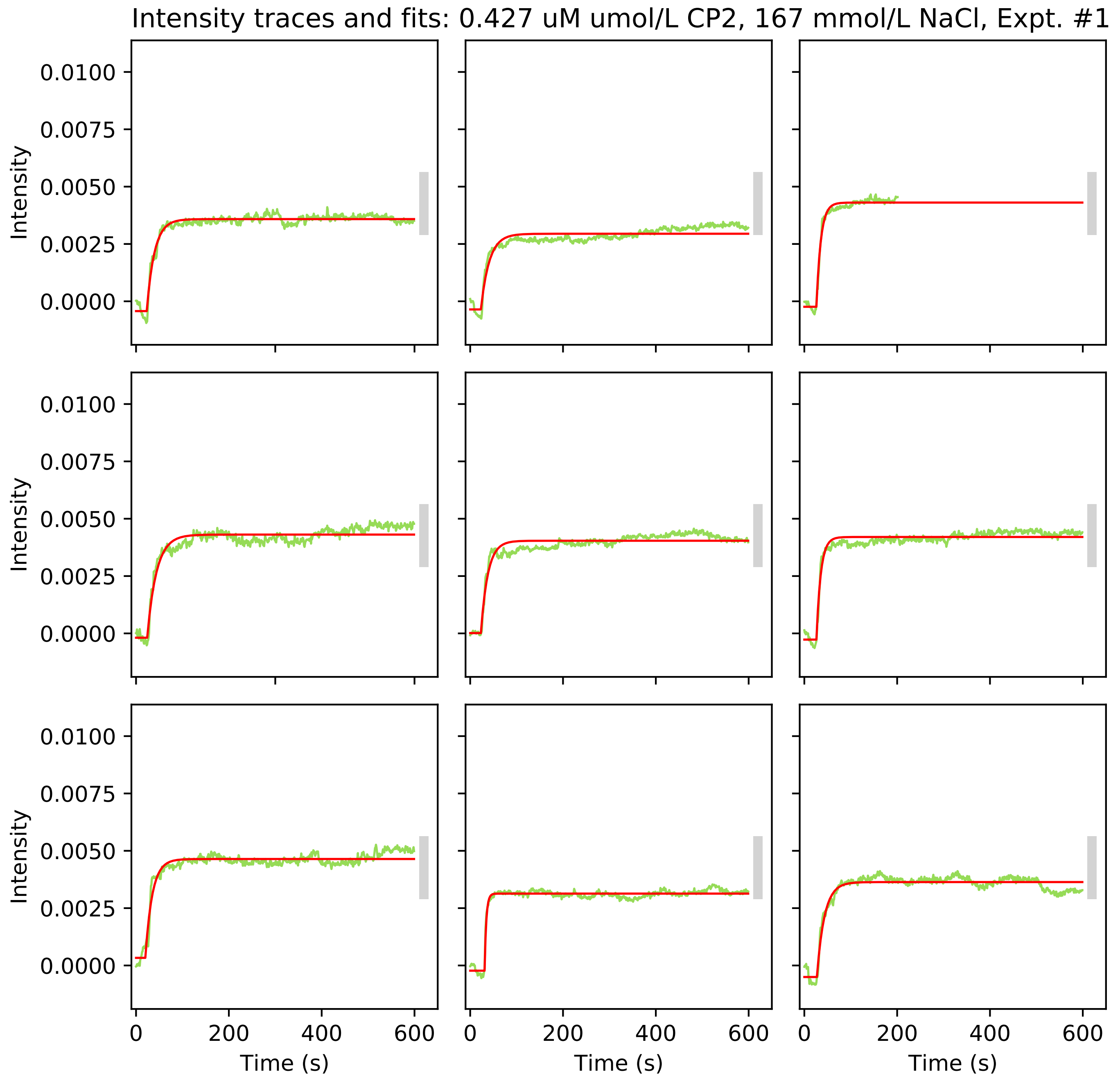

Intensity traces and fits: 0.427 uM umol/L CP2, 167 mmol/L NaCl, Expt. #1
Intensity
0.0100
0.0075
0.0050
0.0025
0.0000
0
200
400
600
Time (s)

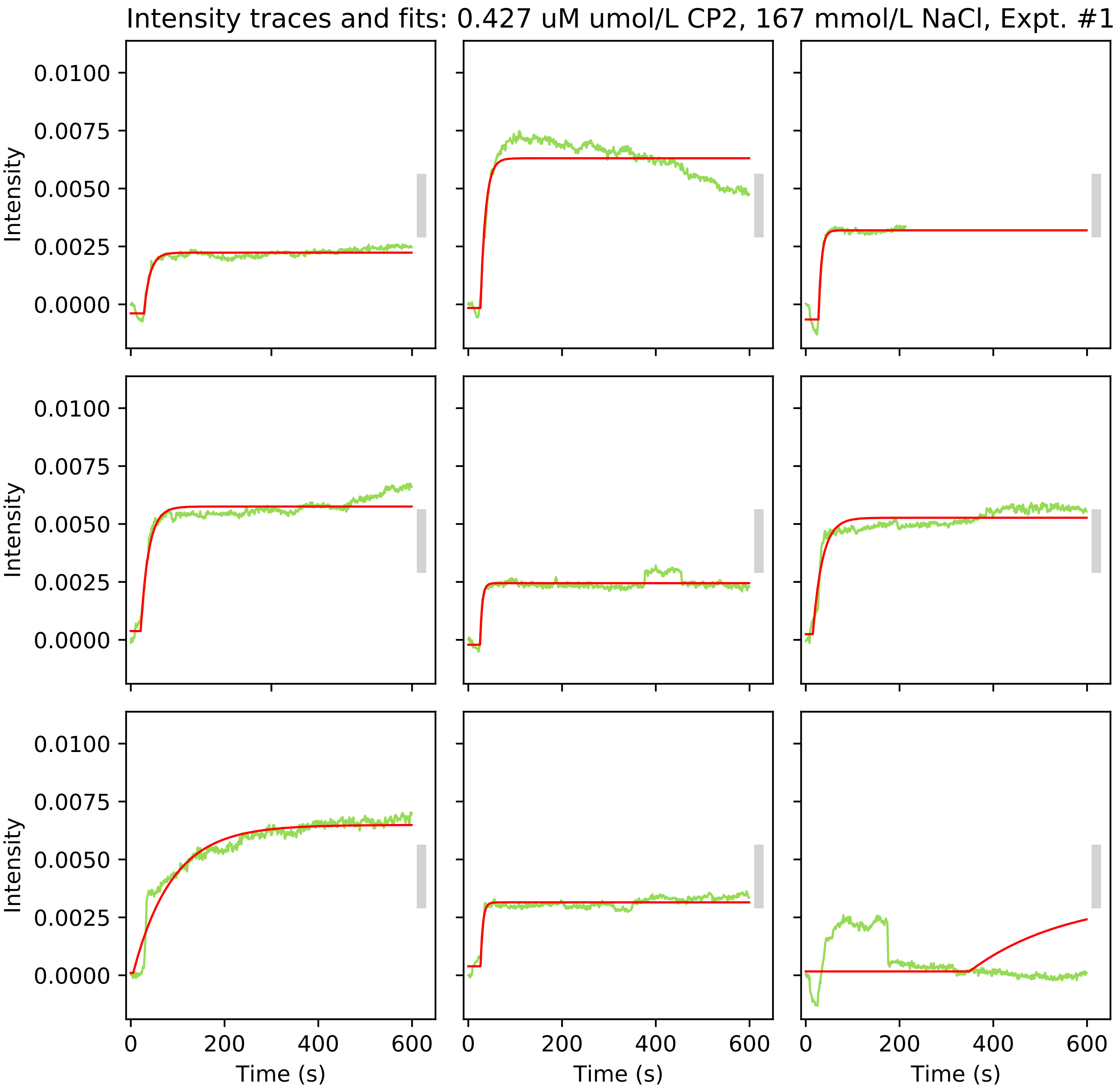
Intensity traces and fits: 0.427 uM umol/L CP2, 167 mmol/L NaCl, Expt. #1
Intensity
0.0100
0.0075
0.0050
0.0025
0.0000
0
200
400
600
Time (s)

Intensity traces and fits: 0.427 uM umol/L CP2, 167 mmol/L NaCl, Expt. #1

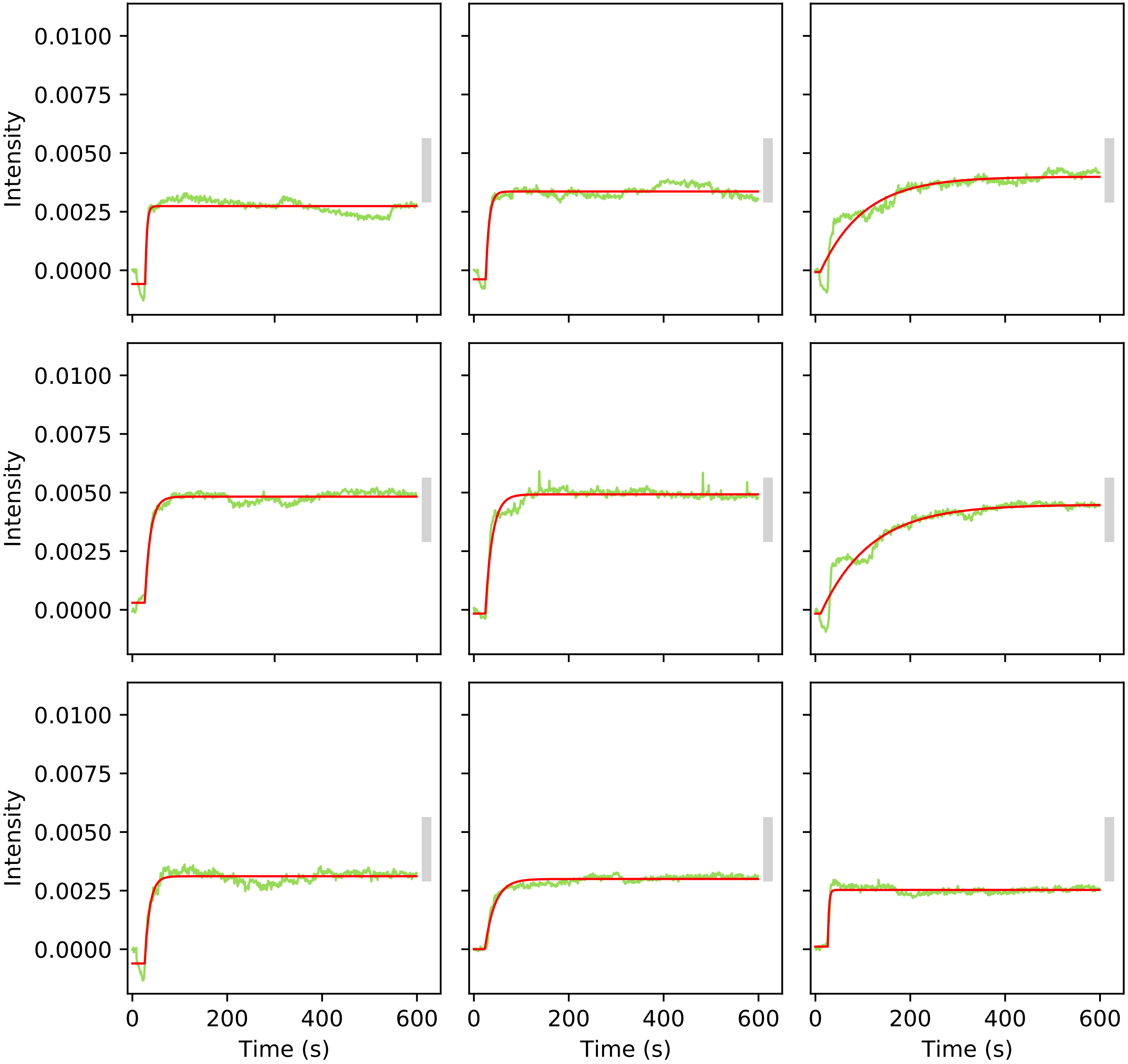

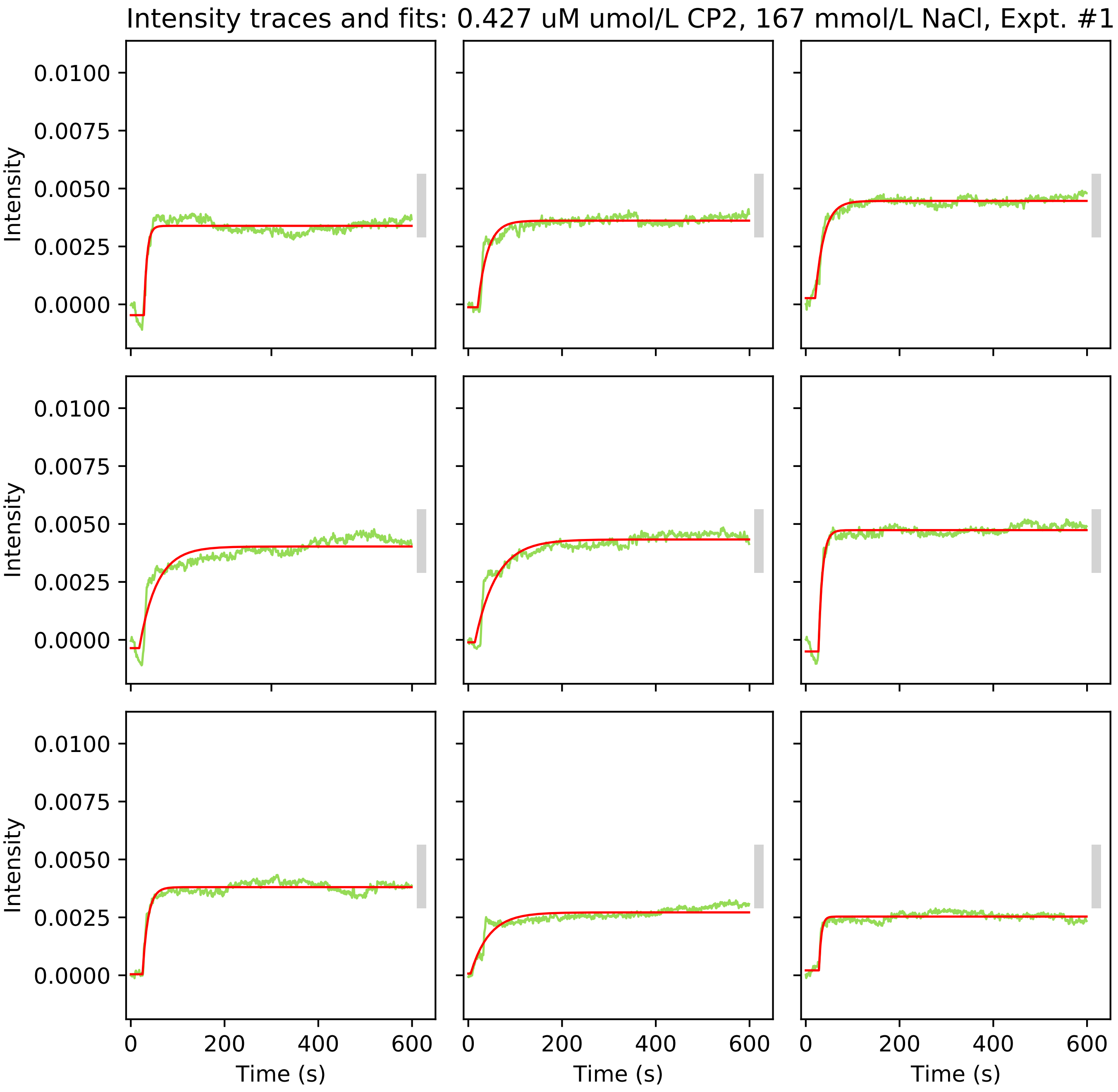

Intensity traces and fits: 0.427 uM umol/L CP2, 167 mmol/L NaCl, Expt. #1
Intensity
0.0100
0.0075
0.0050
0.0025
0.0000
0
200
400
600
Time (s)

Intensity traces and fits: 0.427 uM umol/L CP2, 167 mmol/L NaCl, Expt. #1

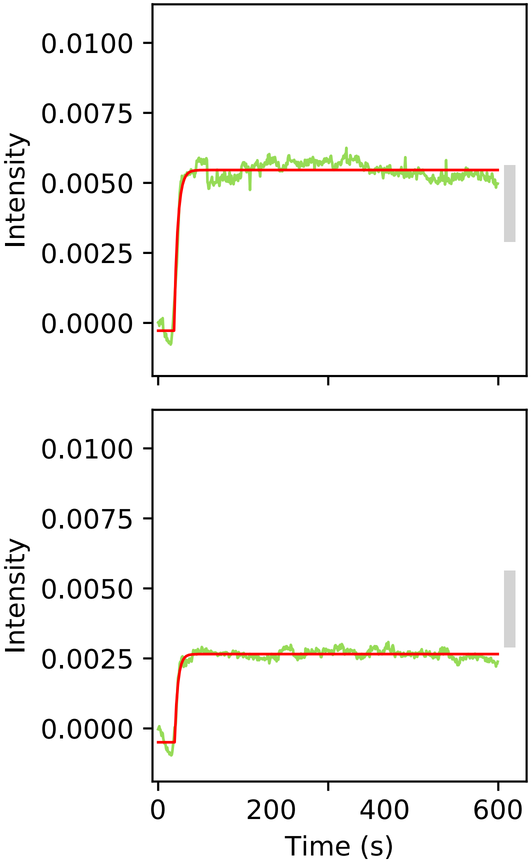

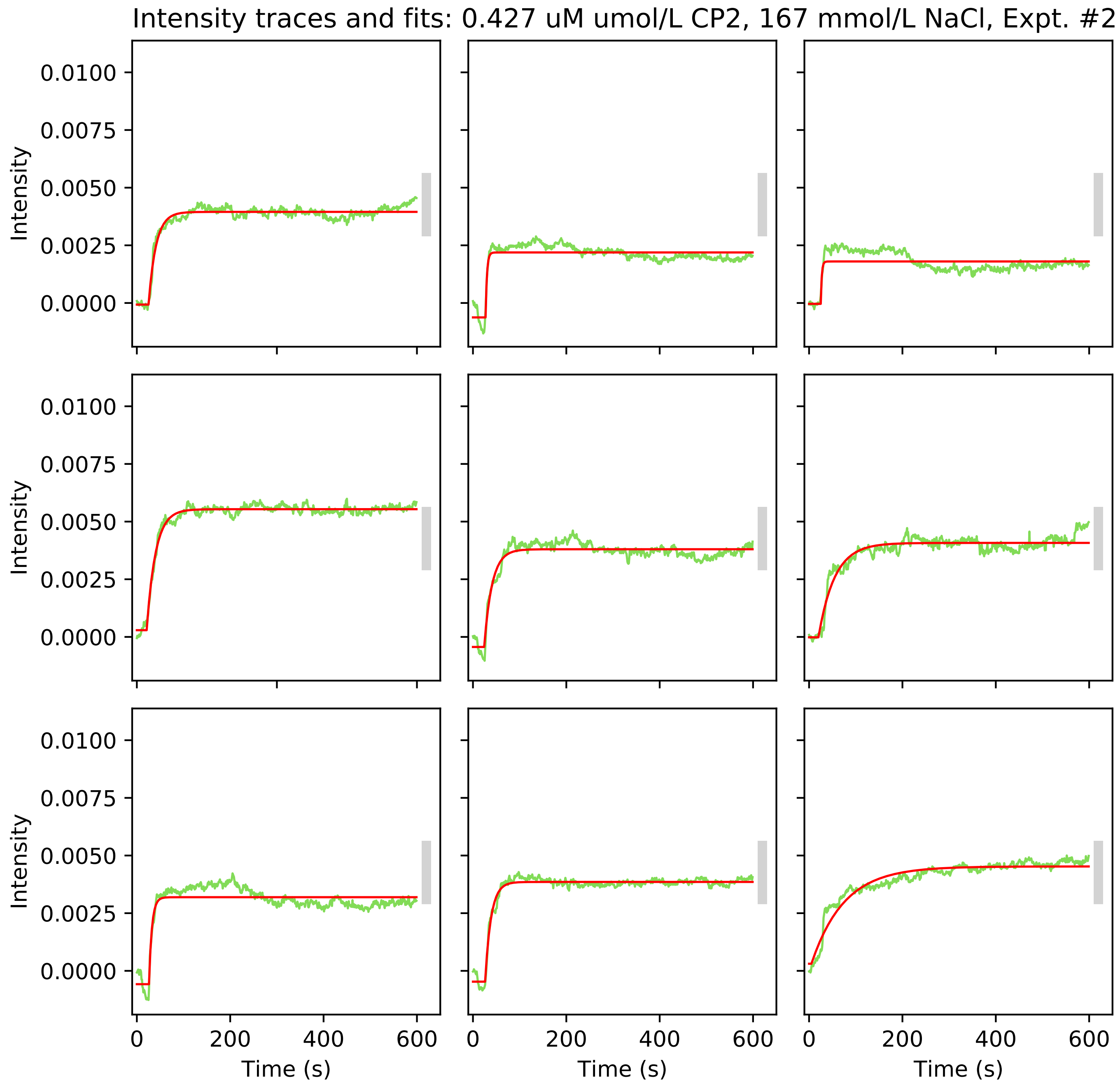

Intensity traces and fits: 0.427 uM umol/L CP2, 167 mmol/L NaCl, Expt. #2
Intensity
0.0100
0.0075
0.0050
0.0025
0.0000
Time (s)
0
200
400
600

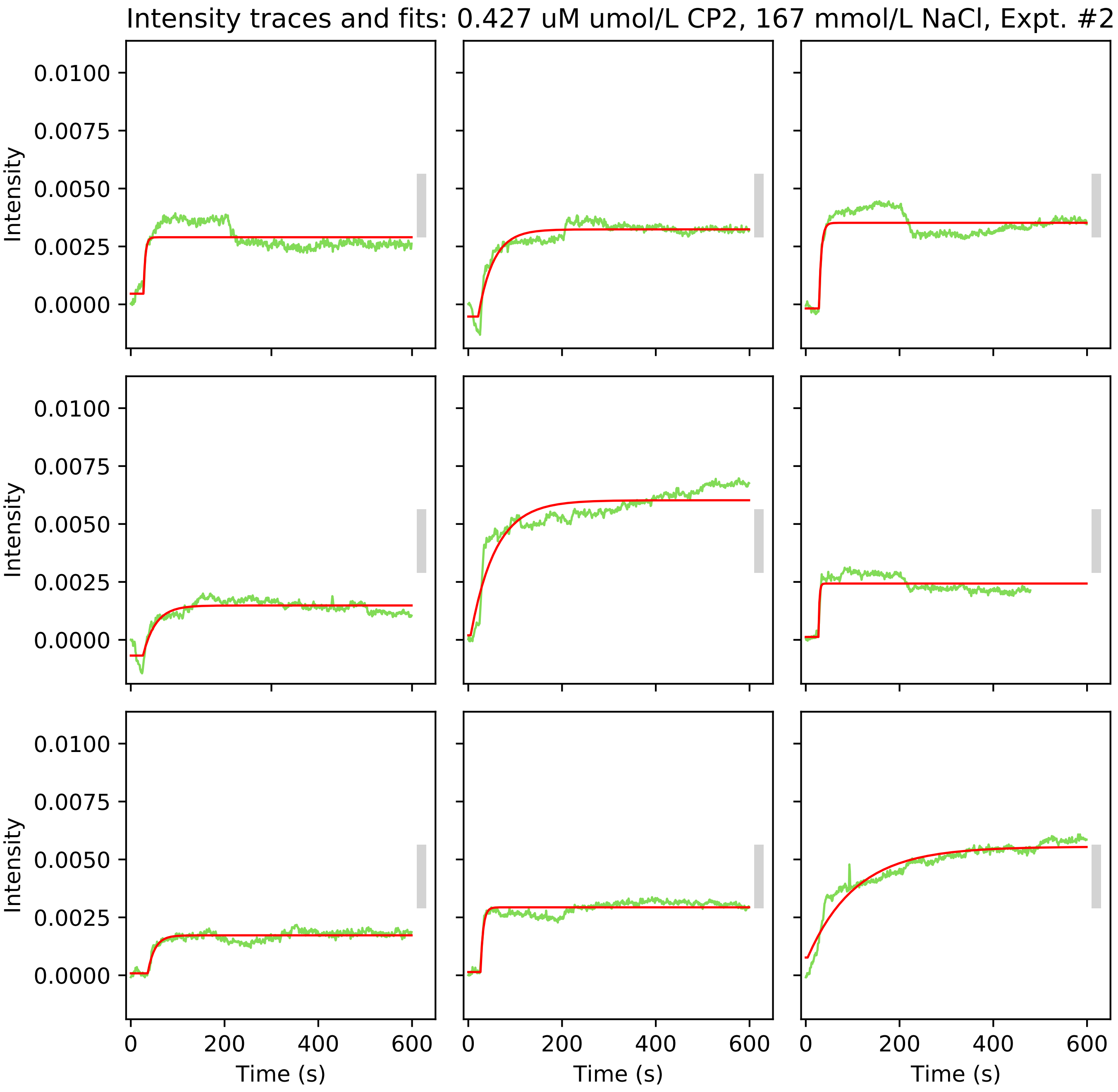
Intensity traces and fits: 0.427 uM umol/L CP2, 167 mmol/L NaCl, Expt. #2
Intensity
0.0100
0.0075
0.0050
0.0025
0.0000
0
200
400
600
Time (s)

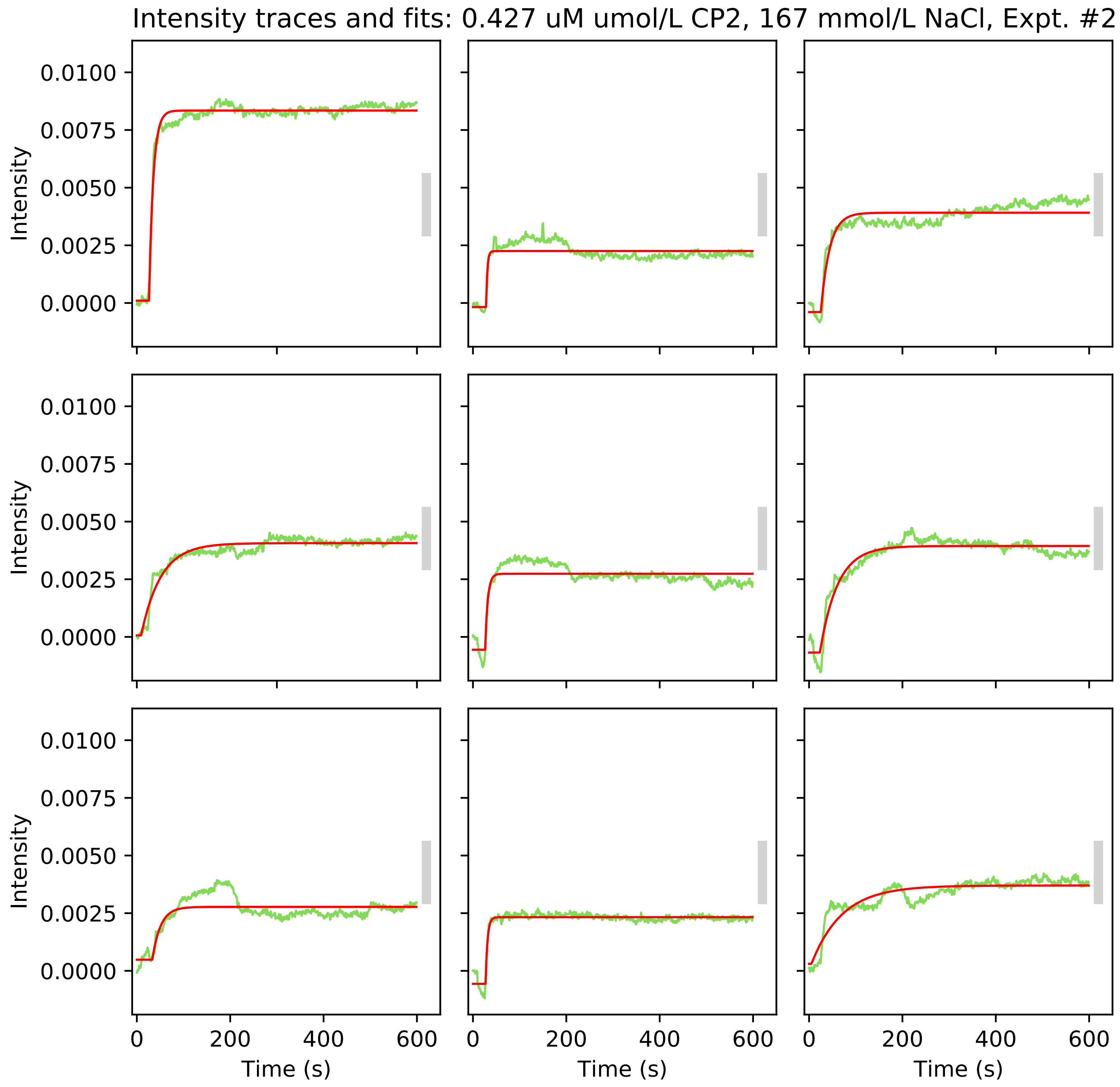
Intensity traces and fits: 0.427 uM umol/L CP2, 167 mmol/L NaCl, Expt. #2
Intensity
0.0100
0.0075
0.0050
0.0025
0.0000
Time (s)
0
200
400
600

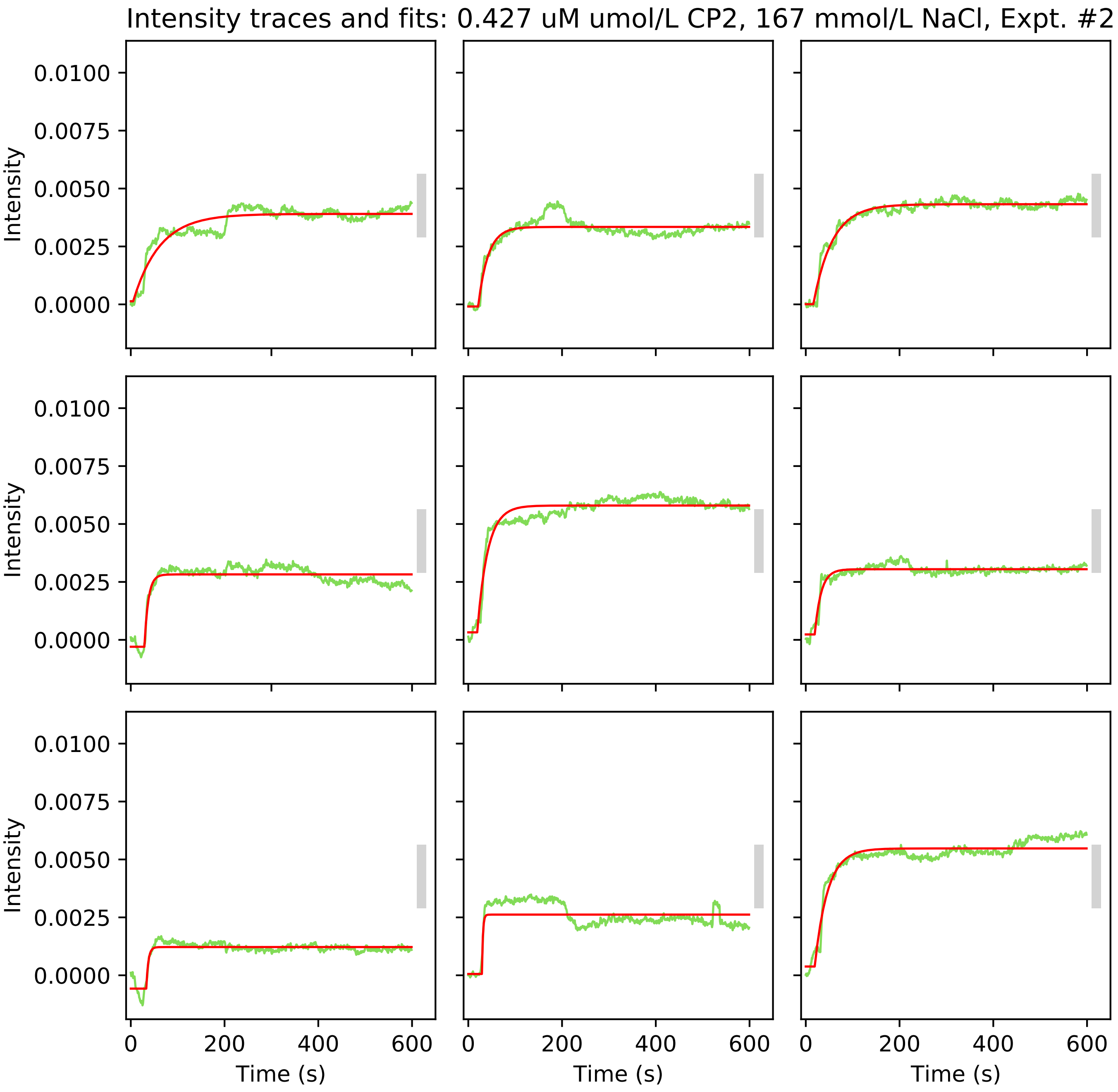

Intensity traces and fits: 0.427 uM umol/L CP2, 167 mmol/L NaCl, Expt. #2
Intensity
0.0100
0.0075
0.0050
0.0025
0.0000
0
200
400
600
Time (s)

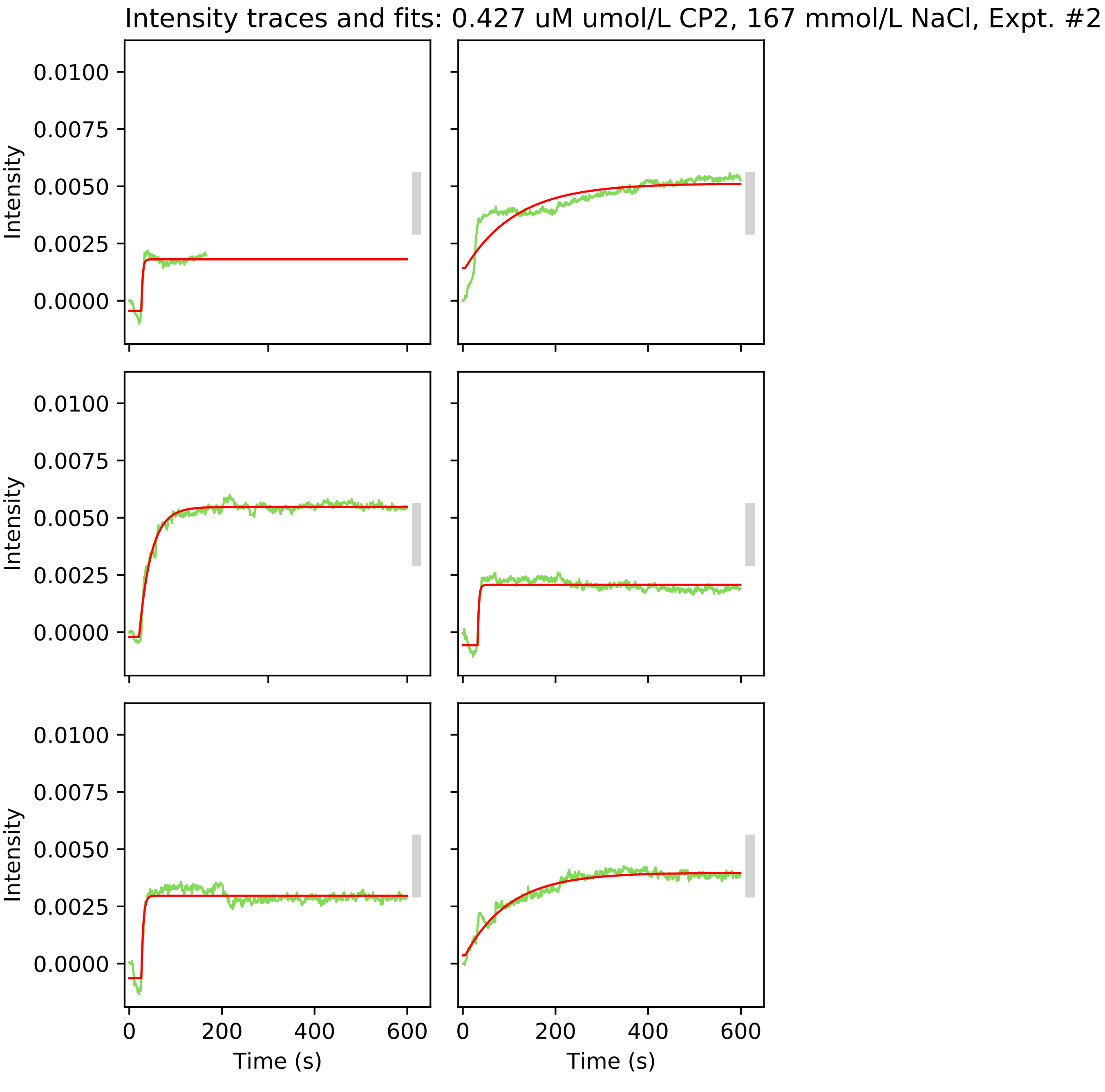
Intensity traces and fits: 0.427 uM umol/L CP2, 167 mmol/L NaCl, Expt. #2
Intensity
0.0100
0.0075
0.0050
0.0025
0.0000
0
200
400
600
Time (s)

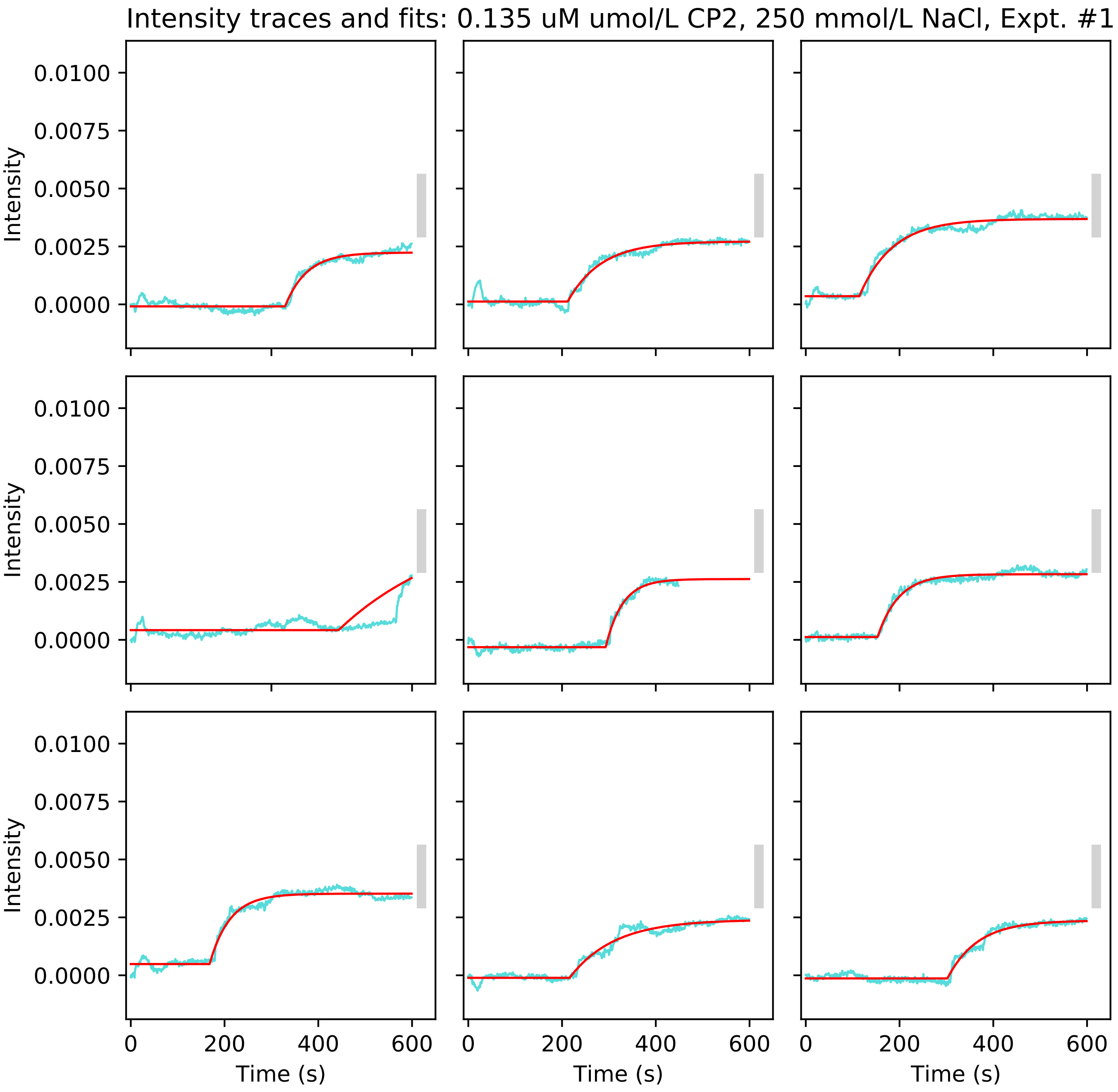

Intensity traces and fits: 0.135 uM umol/L CP2, 250 mmol/L NaCl, Expt. #1
Intensity
0.0100
0.0075
0.0050
0.0025
0.0000
Time (s)
0
200
400
600

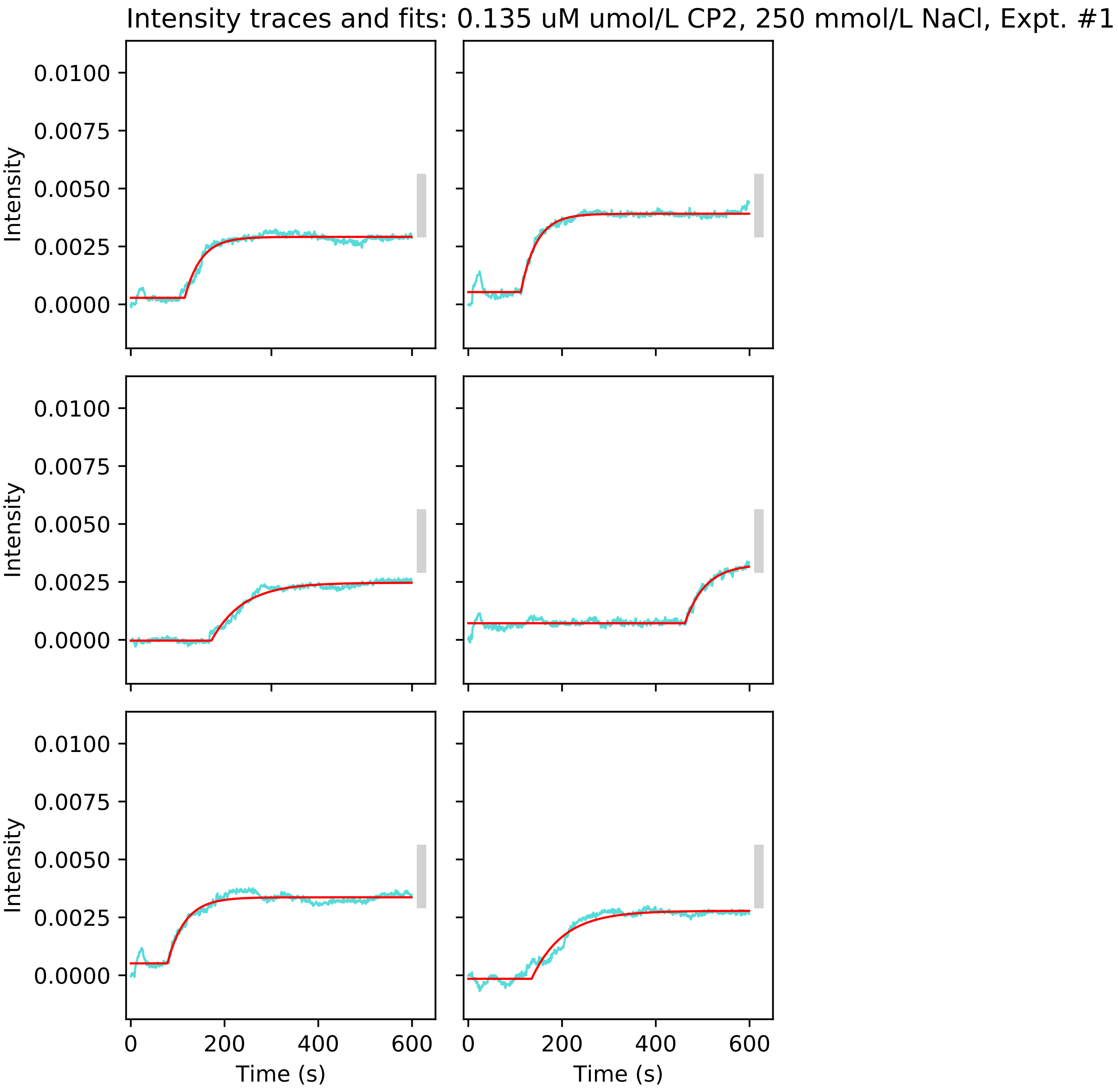
Intensity traces and fits: 0.135 uM umol/L CP2, 250 mmol/L NaCl, Expt. #1
Intensity
0.0100
0.0075
0.0050
0.0025
0.0000
0
200
400
600
Time (s)

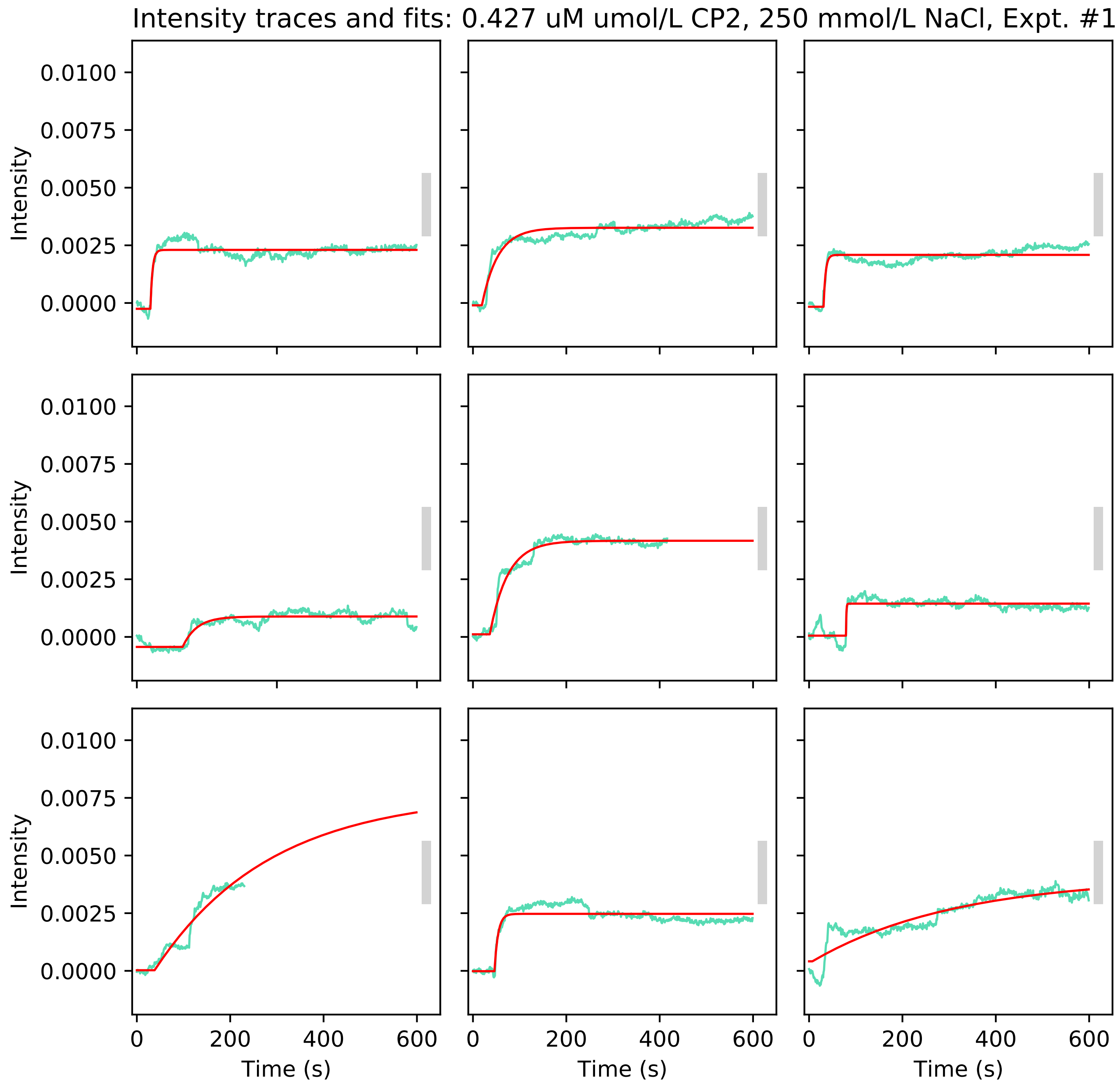

Intensity traces and fits: 0.427 uM umol/L CP2, 250 mmol/L NaCl, Expt. #1
Intensity
0.0100
0.0075
0.0050
0.0025
0.0000
0
200
400
600
Time (s)

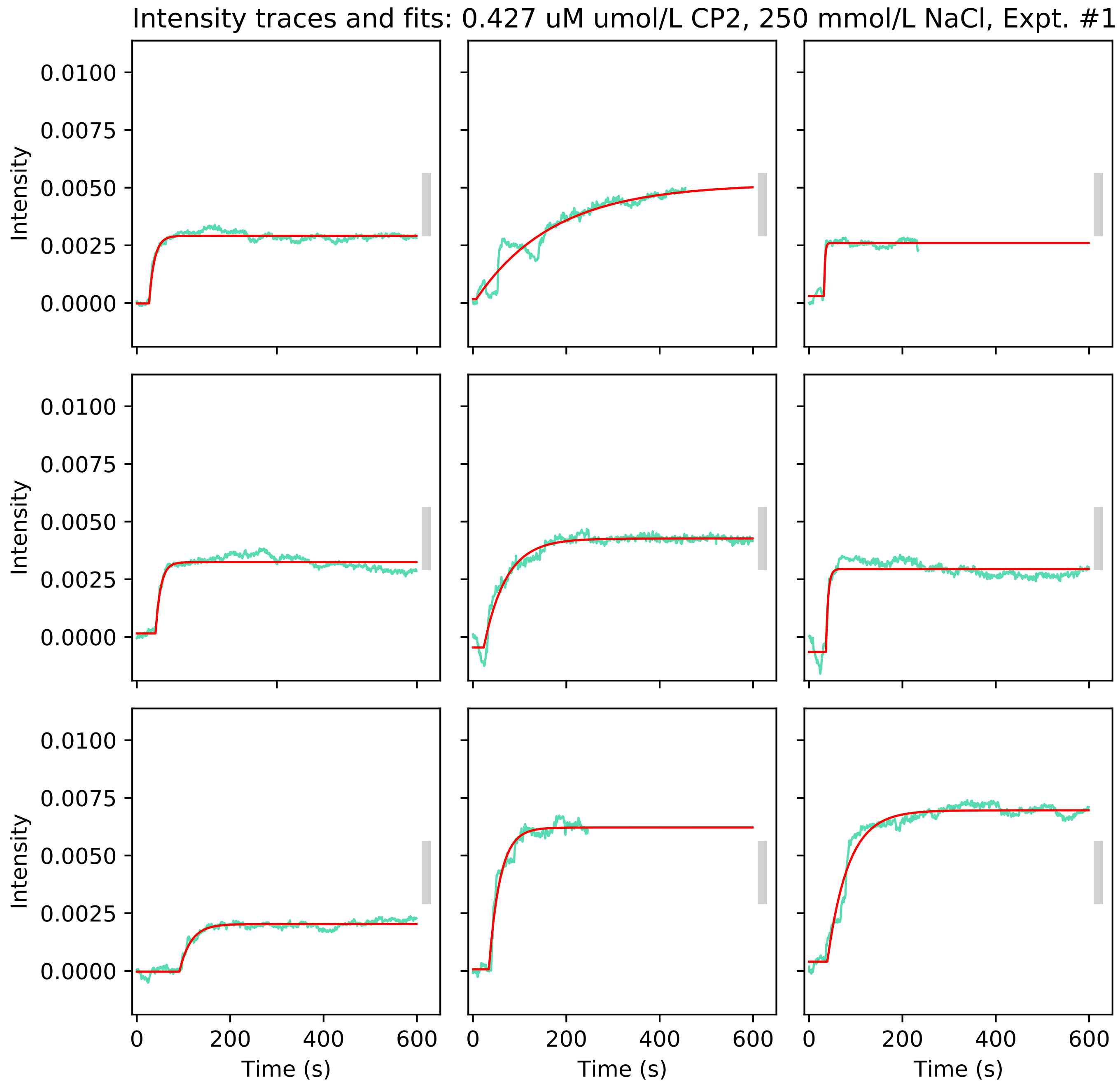

Intensity traces and fits: 0.427 uM umol/L CP2, 250 mmol/L NaCl, Expt. #1
Intensity
0.0100
0.0075
0.0050
0.0025
0.0000
0
200
400
600
Time (s)

Intensity traces and fits: 0.427 uM umol/L CP2, 250 mmol/L NaCl, Expt. #1

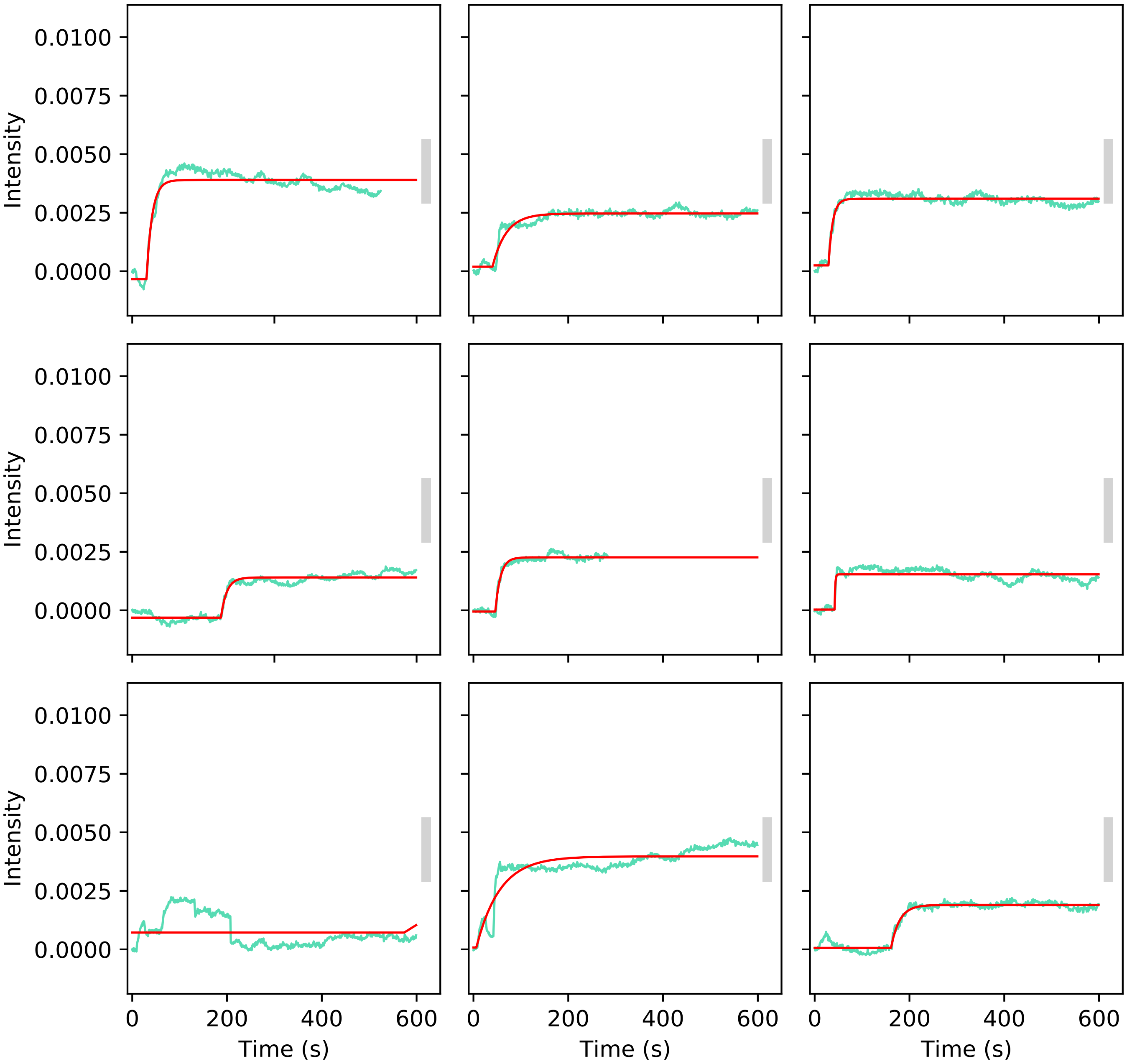

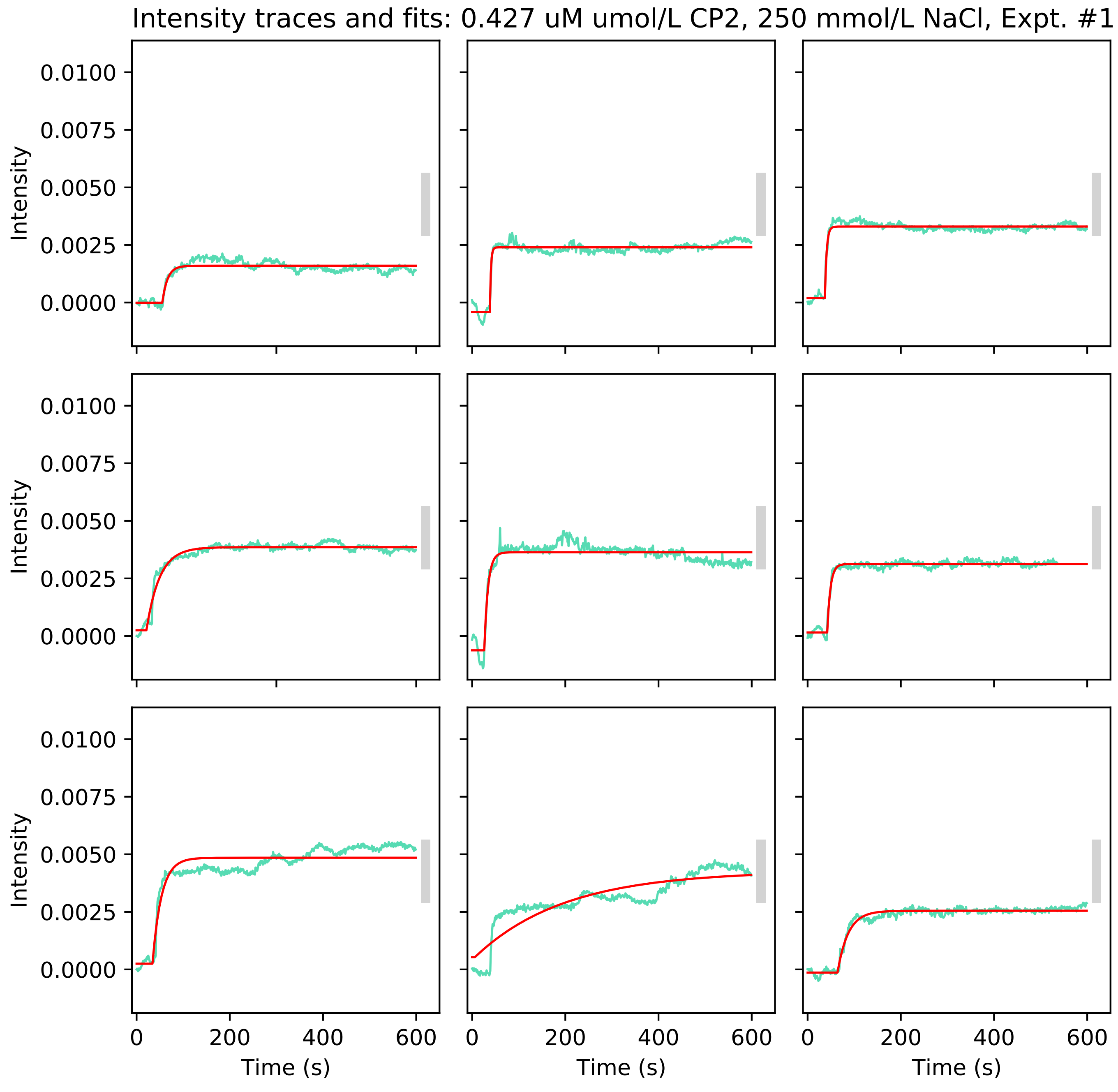
Intensity traces and fits: 0.427 uM umol/L CP2, 250 mmol/L NaCl, Expt. #1
Intensity
0.0100
0.0075
0.0050
0.0025
0.0000
0
200
400
600
Time (s)

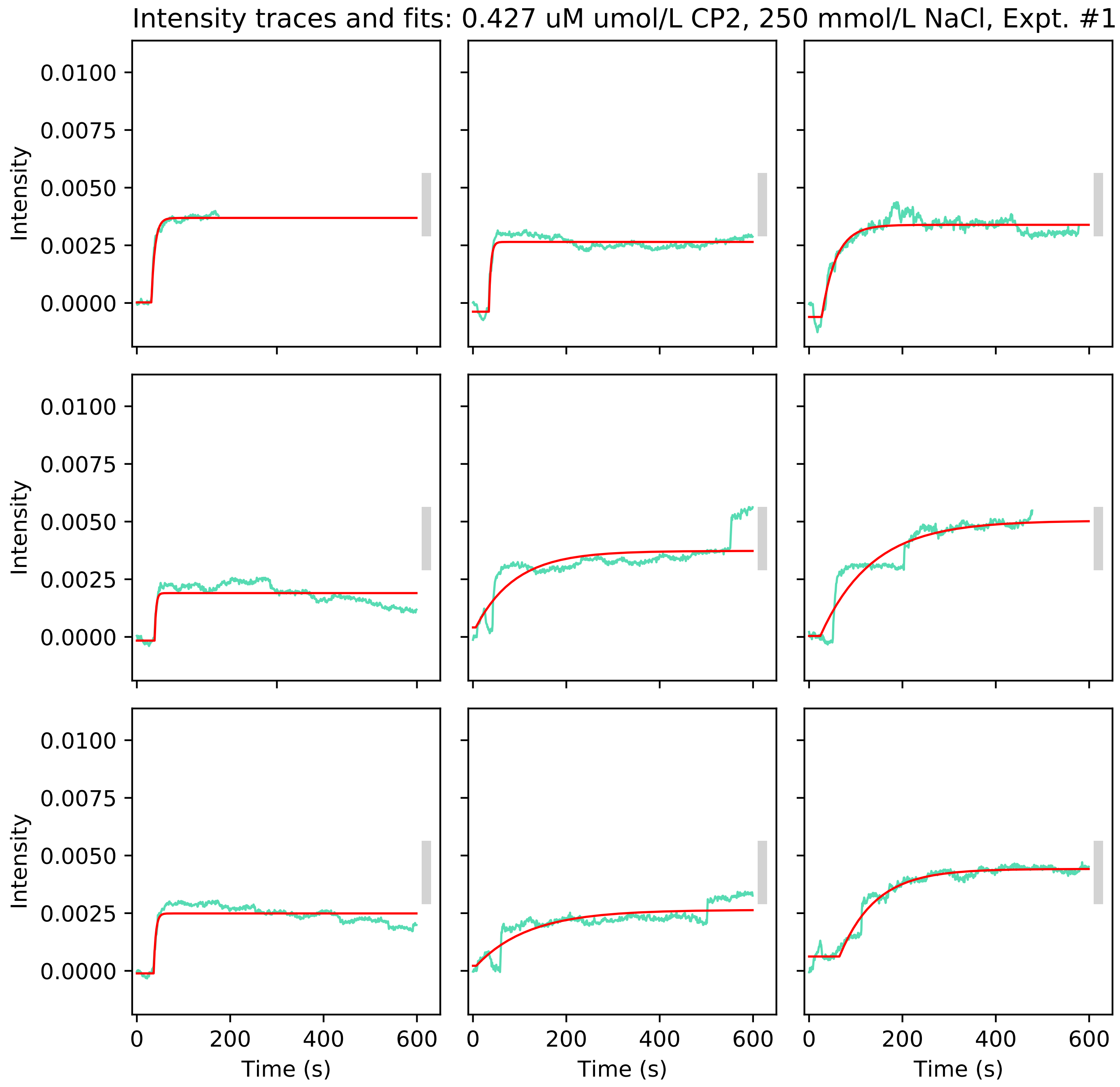

Intensity traces and fits: 0.427 uM umol/L CP2, 250 mmol/L NaCl, Expt. #1
Intensity
0.0100
0.0075
0.0050
0.0025
0.0000
0
200
400
600
Time (s)

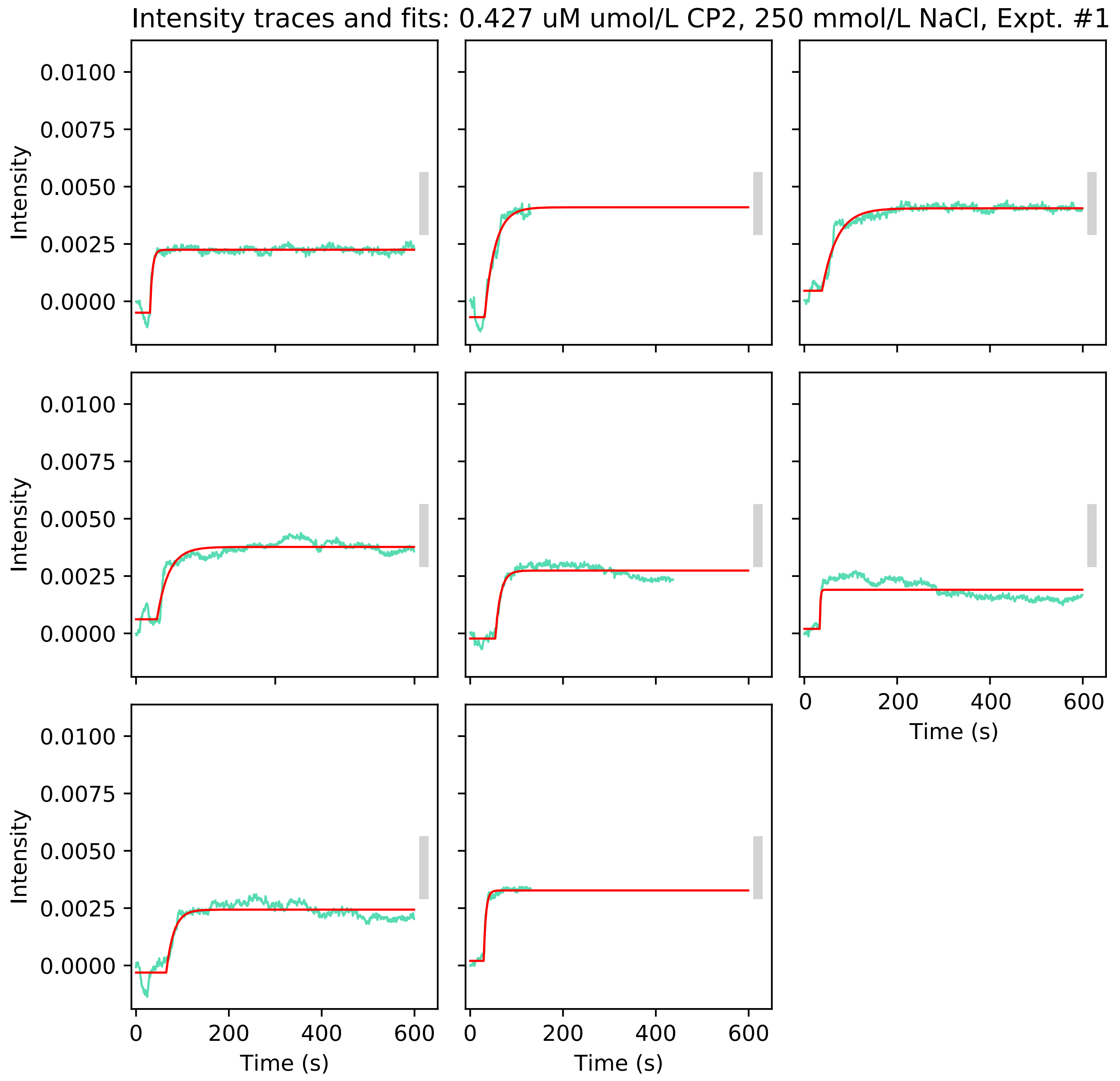

Intensity traces and fits: 0.427 uM umol/L CP2, 250 mmol/L NaCl, Expt. #1
Intensity
0.0100
0.0075
0.0050
0.0025
0.0000
Time (s)
0
200
400
600

Intensity traces and fits: 0.427 uM umol/L CP2, 250 mmol/L NaCl, Expt. #2

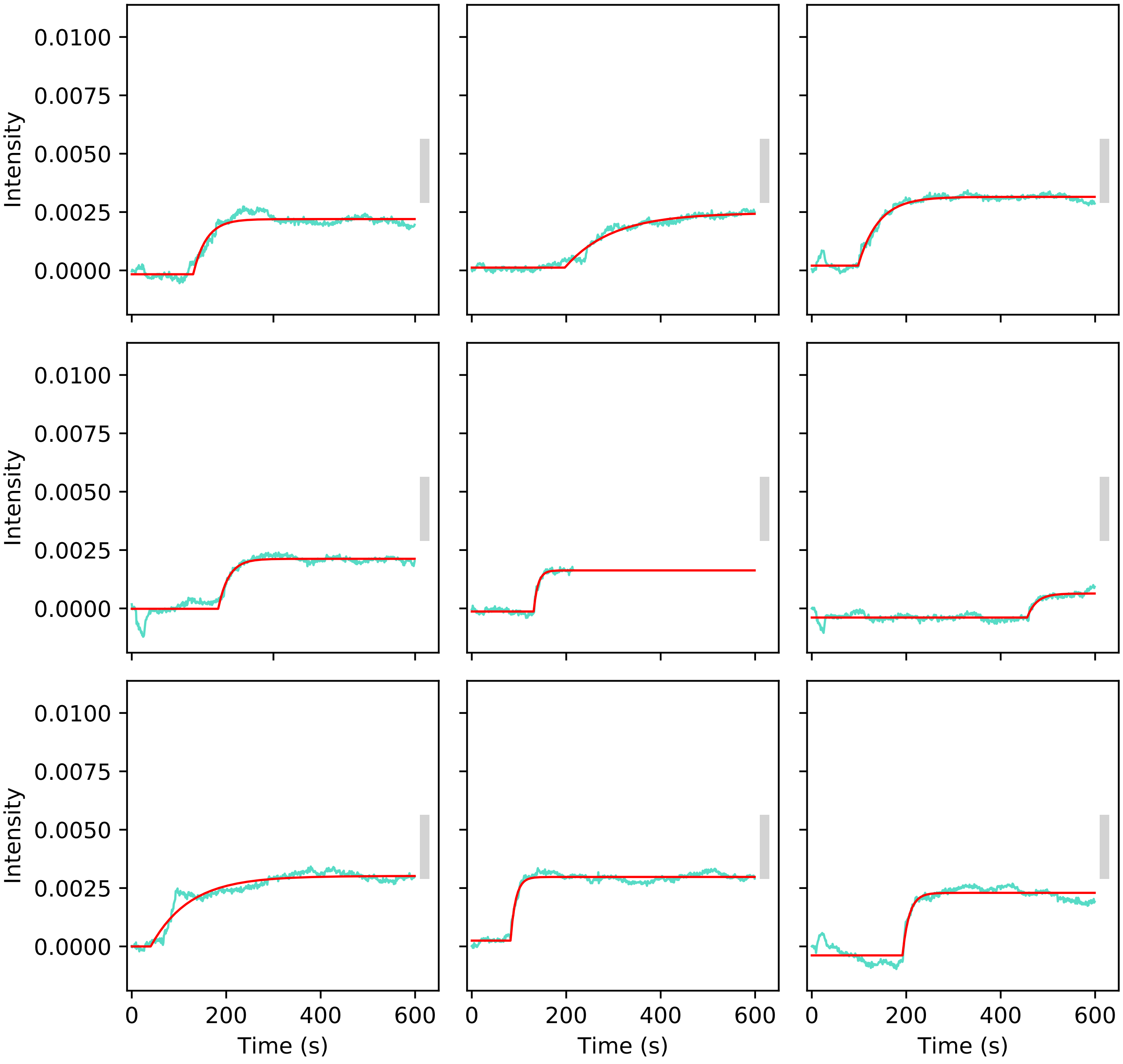

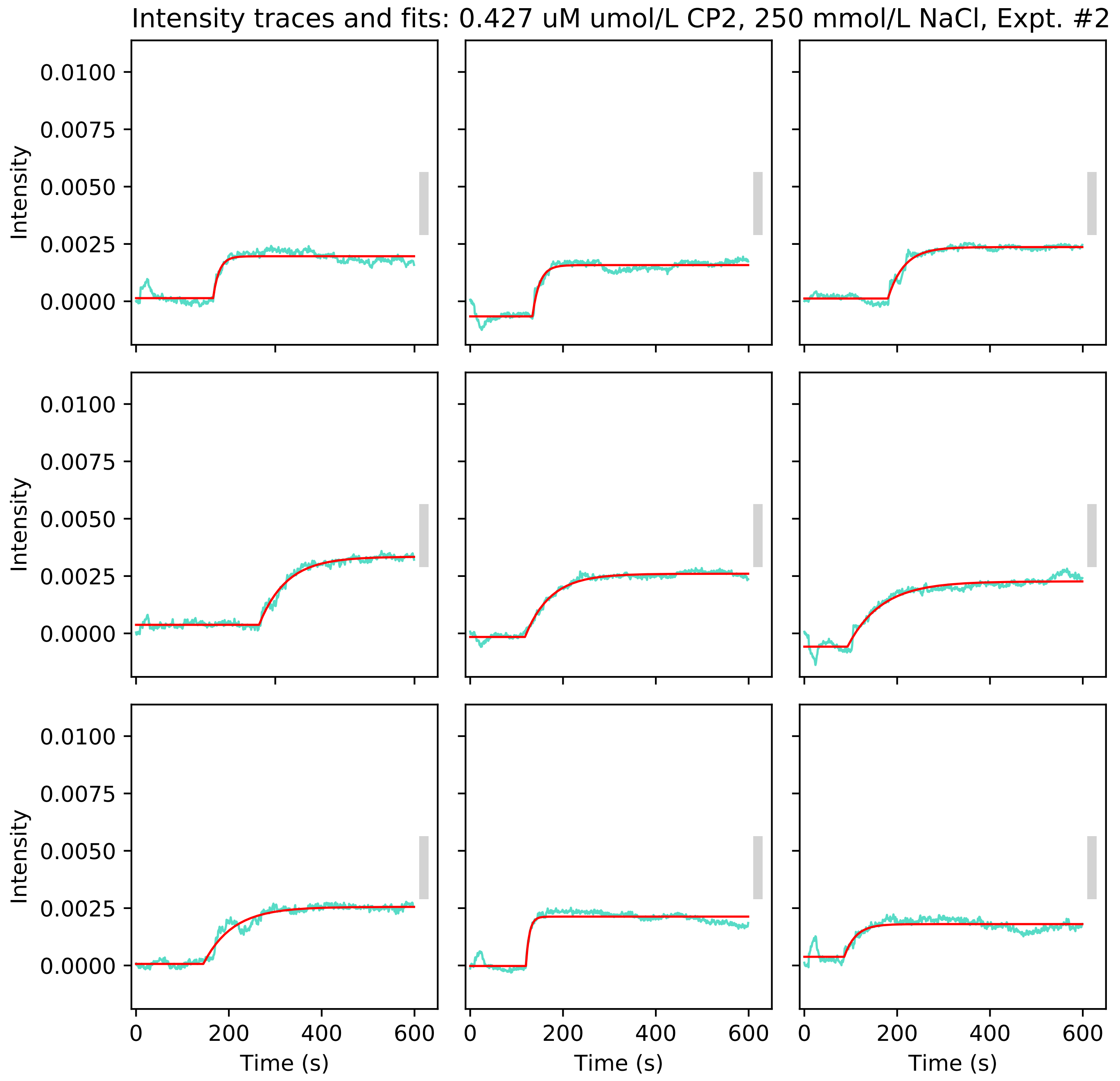

Intensity traces and fits: 0.427 uM umol/L CP2, 250 mmol/L NaCl, Expt. #2
Intensity
0.0100
0.0075
0.0050
0.0025
0.0000
0
200
400
600
Time (s)

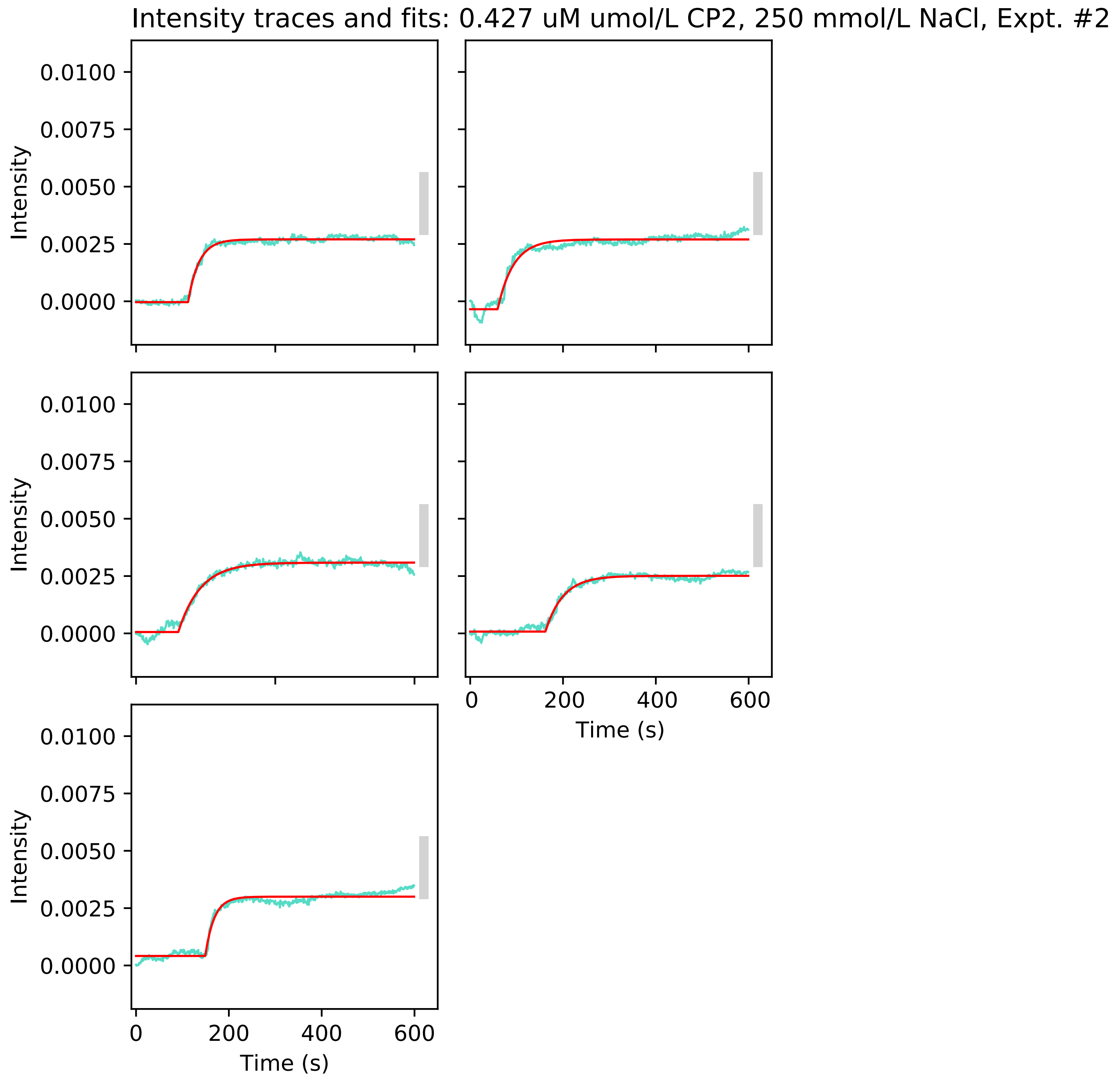

Intensity traces and fits: 0.427 uM umol/L CP2, 250 mmol/L NaCl, Expt. #2
Intensity
0.0100
0.0075
0.0050
0.0025
0.0000
0
200
400
600
Time (s)
